# Low membrane fluidity triggers lipid phase separation and protein segregation *in vivo*

**DOI:** 10.1101/852160

**Authors:** Marvin Gohrbandt, André Lipski, James W. Grimshaw, Jessica A. Buttress, Zunera Baig, Brigitte Herkenhoff, Stefan Walter, Rainer Kurre, Gabriele Deckers-Hebestreit, Henrik Strahl

## Abstract

All living organisms adapt their membrane lipid composition in response to changes in their environment or diet. These conserved membrane-adaptive processes have been studied extensively. However, key concepts of membrane biology linked to regulation of lipid composition including homeoviscous adaptation maintaining stable levels of membrane fluidity, and gel-fluid phase separation resulting in domain formation, heavily rely upon *in vitro* studies with model membranes or lipid extracts. Using the bacterial model organisms *Escherichia coli* and *Bacillus subtilis*, we now show that inadequate *in vivo* membrane fluidity interferes with essential complex cellular processes including cytokinesis, envelope expansion, chromosome replication/segregation and maintenance of membrane potential. Furthermore, we demonstrate that very low membrane fluidity is indeed capable of triggering large-scale lipid phase separation and protein segregation in intact, protein-crowded membranes of living cells; a process that coincides with the minimal level of fluidity capable of supporting growth. Importantly, the *in vivo* lipid phase separation is not associated with a breakdown of the membrane diffusion barrier function, thus explaining why the phase-separation process induced by low fluidity is biologically reversible.

## Introduction

Biological membranes are complex arrangements predominantly composed of lipids and both integral and surface-attached proteins (Nicholson, 2104). The primordial function of biological membranes was likely to act as a simple, semipermeable diffusion barrier separating the cell from its environment, and genomes from each other (Chen & Valde, 2010). Later, membranes and membrane proteins evolved to fulfil a multitude of cellular functions including transport, respiration, and morphogenesis. Since the physicochemical state of biological membranes is highly sensitive to changes in the environment including temperature, osmolarity, salinity, pH or diet (Ernst *et al*, 2016; Hazel, 1995; Razin, 1967), careful homeostatic regulation of key membrane parameters such as thickness or fluidity is vital for cell function (Ernst *et al*, 2016; Harayama & Riezman, 2018; Levental *et al*, 2020; Parsons & Rock, 2013).

Arguably, the best studied membrane-adaptive process is homeoviscous adaptation that acts upon changes in temperature (Ernst *et al*, 2016; Hazel, 1995; Parsons & Rock, 2013). With increasing temperature, lipid bilayers exhibit reduced head group packing, increased fatty acid disorder and increased fluidity in terms of increased rotational and lateral diffusion of molecules (Chapman, 1975; Heimburg, 2007). While all of these membrane parameters are closely interconnected, membrane fluidity is thought to be the key property actively maintained at stable levels that optimally support vital membrane functions through homeoviscous adaptation processes (Ernst *et al*, 2016; Hazel, 1995; Parsons & Rock, 2013). All living organisms achieve this by actively adapting their lipid composition. Most commonly, this is achieved by altering the content of lipids carrying fluidity-promoting unsaturated fatty acids (UFA) or branched chain fatty acids (BCFA) and fluidity-reducing saturated fatty acids (SFA), respectively, thereby counteracting shifts in membrane fluidity (Ernst *et al*, 2016; Hazel, 1995; Diomandé *et al*, 2015).

While adaptive changes in lipid fatty acid composition as well as the regulatory processes involved are increasingly well characterised (Ernst *et al*, 2018; Mansilla *et al*, 2004; Ballweg *et al*, 2020), the cellular consequences of inadequate membrane fluidity are significantly less understood. Sufficiently high membrane fluidity has been implicated in promoting folding, catalytic activity, and diffusion of membrane proteins (Andersen & Koeppe, 2007; Lee, 2004). Too high membrane fluidity, in turn, has been shown to increase proton permeability *in vitro* (Rossignol *et al*, 1982; van de Vossenberg, 1999), thus potentially hampering with efficient ion homeostasis and energy conservation (Valentine, 2007) while too low membrane fluidity impedes respiration due to reduced ubiquinone diffusivity (Budin *et al*, 2018). However, our understanding of the behaviour of biological membranes upon changing fluidity is predominantly based on *in vitro* and *in silico* studies with simplified model lipids, or *in vitro* studies with either natural lipid extracts or isolated membranes (Baumgart *et al*, 2007; Nickels *et al*, 2017; Schäfer *et al*, 2011).

One of the fascinating properties of lipids is their ability to undergo phase transitions between distinct configurations that differ in terms of ability to form bilayers, membrane thickness and degree of lipid packing (Chapman, 1975). The biologically relevant bilayer-forming lipid phases are: (*i*) the liquid-disordered phase characterised by low packing density and high diffusion rates that forms the regular state of biological membranes, (*ii*) the cholesterol/hopanoid-dependent liquid-ordered phase that forms nanodomains (lipid rafts) found in biological membranes, both representing different fluid phases and (*iii*) the gel or solid phase characterised by dense lipid packing with little lateral or rotational diffusion, which is generally assumed to be absent in biologically active membranes (Schmid, 2017; Veatch, 2007; Sáenz *et al*, 2015). In fact, the temperature associated with gel phase formation has been postulated to define the lower end of the temperature range able to support vital cell functions (Burns *et al*, 2017; Drobnis *et al*, 1993; Ghetler *et al*, 2005), and maintaining biological membranes in the correct phase (homeophasic regulation) has been suggested as an alternative rationale behind temperature-dependent lipid adaptation (Hazel, 1995). Finally, lipid phases can co-exist, resulting in separated membrane areas exhibiting distinctly different composition and characteristics (Baumgart *et al*, 2007; Nickels *et al*, 2017; Elson *et al*, 2010; Heberle & Feigenson, 2011; Shen *et al*, 2017). This principal mechanism of lipid domain formation is best studied in the context of lipid rafts (Lingwood & Simons, 2010; Shaw *et al*, 2021). Here the co-existence of fluid liquid-disordered and liquid-ordered phases, and the associated protein segregation, has been demonstrated in membranes of living eukaryotic cells (Toulmay & Prinz, 2013; Shelby *et al*, 2021). In contrast, comprehensive *in vivo* studies on gel-fluid lipid phase separation in live cells have been challenging due to the tendency of cholesterol/hopanoids to suppress gel-fluid phase transitions (Heberle & Feigenson, 2011), and due to the difficulty to modify the membrane fatty acid composition and, thus, fluidity without inducing lipotoxicity (Shen *et al*, 2017; Budin *et al*, 2018).

While *in vitro* and *in silico* approaches with simplified lipid models have provided detailed insights into the complex physicochemical behaviour of lipid bilayers, testing the formed hypotheses and models in the context of protein-rich biological membranes is now crucial. Bacteria tolerate surprisingly drastic changes in their lipid composition and only possess one or two membrane layers as part of their cell envelope. Consequently, bacteria are both a suitable and a more tractable model to study the fundamental biological process linked to membrane fluidity and phase separation *in vivo*.

We analysed the biological importance of membrane homeoviscous adaptation in *Escherichia coli* (phylum *Proteobacteria*) and *Bacillus subtilis* (phylum *Firmicutes*), respectively. These organisms were chosen due to their prominence as Gram-negative and Gram-positive model organisms, and the different archetypes of membrane fatty acid composition (straight vs. branched chain fatty acids) they represent. We have established protocols that allow the fatty acid composition of both organisms to be progressively altered and the cellular consequences to be directly monitored in growing cells. This approach allowed us to address three central questions linked to homeostatic regulation of membrane composition and fluidity: (*i*) what are the cellular consequences of an inadequate level of membrane fluidity that necessitate the extensive and conserved homeostatic regulatory processes, (*ii*) how do changes in lipid fatty acid composition translate to changes in membrane fluidity of living cells and (*iii*) what is the lipid phase behaviour in living cells with protein-crowded membranes and intact lipid domain organization?

Our results demonstrate that too low membrane fluidity results in growth arrest in both organisms, which is accompanied by severe disturbances of the cell morphogenesis and ion homeostasis. Furthermore, too low fluidity triggers a striking, large-scale lipid phase separation into liquid-disordered and gel phase membranes, accompanied by segregation of otherwise disperse membrane proteins such as ATP synthase and glucose permease. Our results revealed that phase separation between liquid-disordered and gel state membranes is associated with loss of essential membrane functions, thereby limiting the range of membrane fluidity able to support life. At last, our findings demonstrating that gel-liquid phase separation and associated membrane protein segregation indeed occurs in protein-crowded, native plasma membranes of living cells, are fully consistent with the comparable phenomena observed in *in vitro* and *in silico* model systems (Baumgart *et al*, 2007; Veatch, 2007; Schäfer *et al*, 2011; Lingwood & Simons, 2010; Domański *et al*, 2012). Thus, the results provide strong *in vivo* support for the general validity of the respective models.

## Results

### Depletion of BCFAs in *B. subtilis*

To modify the fatty acid composition in *B. subtilis*, we constructed a Δ*bkd* Δ*des* deletion strain (Table S1). The *bkd* operon encodes enzymes catalysing the conversion of branched chain amino acids into intermediates for BCFA synthesis (Debarbouille *et al*, 1999). The lack of this activity can be complemented by supplementation with precursors 2-methylbutyric acid (MB) or isobutyric acid (IB) (Boudreaux *et al*, 1981; Kaneda, 1977). This provides the experimental means to control the lipid *iso*- and *anteiso*-BCFA composition (Figure S1A) normally responsible for the homeostatic adaptation of membrane fluidity in response to environmental changes (Diomandé *et al*, 2015). In addition, the strain is deficient for the lipid desaturase Des to prevent rapid adaptation of membrane fluidity by converting SFAs or BCFAs into UFAs (Diomandé *et al*, 2015). In the remaining text, the *B. subtilis* strain is labelled “Δ*bkd*” for simplicity.

We compared growth of *B. subtilis* 168 used as wild type (WT) and Δ*bkd* cells at 37°C upon supplementation with BCFA precursors MB or IB (Figure 1A). While BCFA precursors had little impact on growth of WT cells, the auxotrophic Δ*bkd* strain only grew in the presence of either of the precursors. Corresponding fatty acid analyses revealed large shifts in the composition of the Δ*bkd* strain depending on the supplied precursor (Figures 1B and S1B). As expected, cells supplemented with MB exhibited a high content (77%) of *anteiso*-BCFAs, whereas cells grown with IB showed a high content of *iso-*BCFAs (77%). To obtain cells depleted for both BCFA types, cells were grown in the presence of IB, followed by wash and incubation in <u>p</u>recursor-<u>f</u>ree (PF) medium. This precursor depletion leads to growth arrest after about 90 min (Figure S1B), corresponding to an accumulated SFA content of ∼50% (Figure 1B).

**Figure 1.**
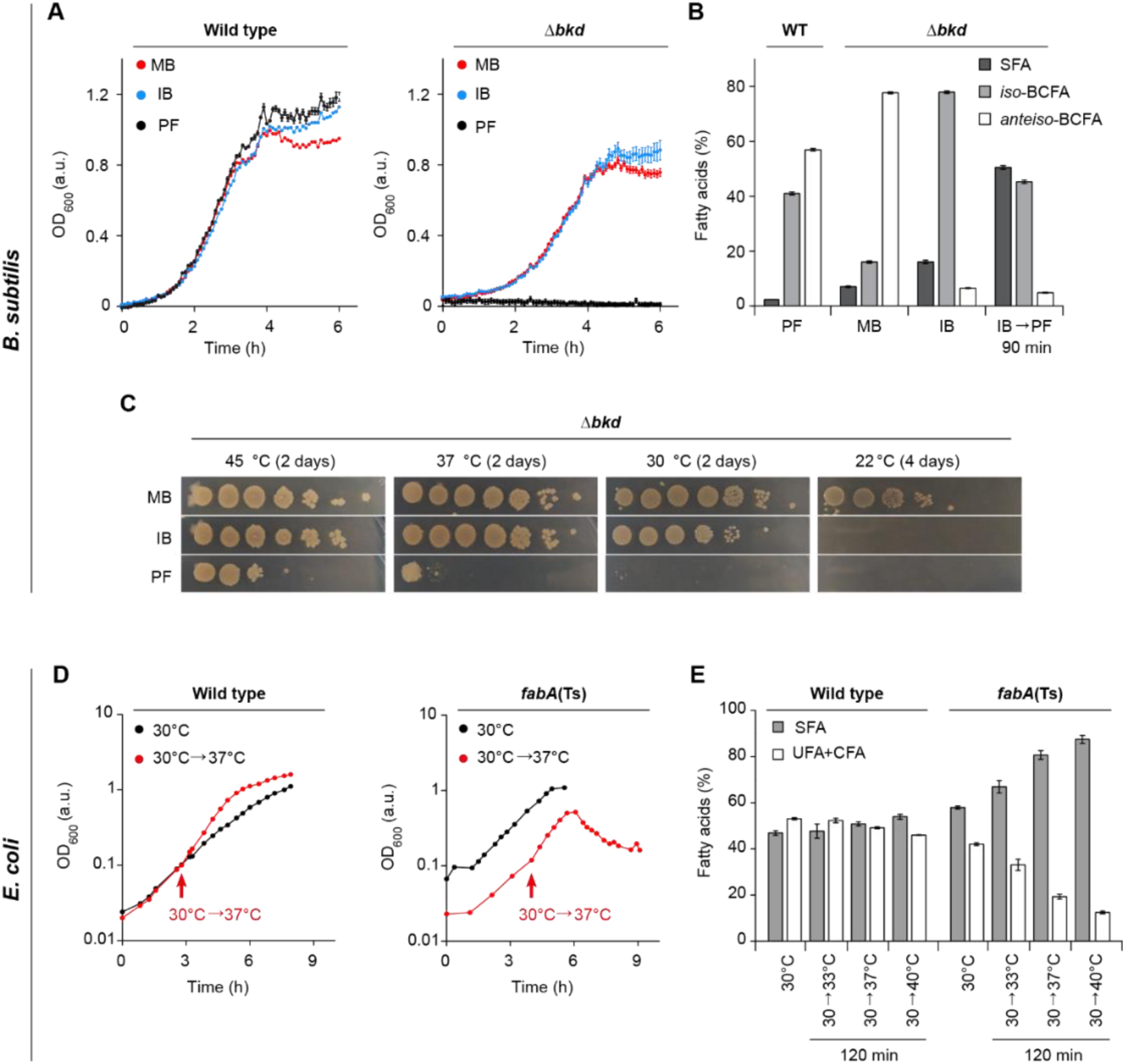
Membrane fatty acid composition-dependent growth of *B. subtilis* and *E. coli*. (**A**) Growth of *B. subtilis* WT and fatty acid precursor-auxotrophic Δ*bkd* cells in medium supplemented with precursor 2-methyl butyric acid (MB), isobutyric acid (IB) or grown precursor-free (PF). (**B**) Fatty acid composition of *B. subtilis* WT cells grown in PF medium, and Δ*bkd* grown with MB, IB or depleted for precursor for 90 min (IB→PF). For detailed analyses, see Figure S1B. (**C**) Temperature-dependent growth of *B. subtilis* Δ*bkd* on solid medium in serial 10-fold dilutions. For comparison between WT, Δ*des* and Δ*bkd* Δ*des* cells, see Figure S1C. (**D**) Temperature-dependent growth behaviour of *E. coli* WT and *fabA*(Ts), including a shift from 30°C to 37°C as non-permissive temperature of *fabA*(Ts). (**E**) Fatty acid composition of *E. coli* WT and *fabA*(Ts) cells grown at 30°C and shifted to different temperatures for 120 min. For detailed analyses, see Figures S2B and S2C. **Data information**: (**A**) The diagram depicts mean and <u>s</u>tandard <u>d</u>eviation (SD) of technical triplicates for each strain. (**B, E**) The histograms depict mean and SD of biological triplicates. (**C, D**) The experiments are representative of three independent repeats. CFA, <u>c</u>yclopropane <u>f</u>atty <u>a</u>cid. Strains used: (**A**-**C**) *B. subtilis* 168, HS527; (**D, E**) *E. coli* MG1, MG4 (strains Y-Mel and UC1098, respectively, additionally encoding fluorescent ATP synthase (F_O_F_1_ *a*-mNG)).

Analyses of Δ*bkd* cells grown at different growth temperatures with different BCFA precursors (Figures 1C and S1C) indicated that only MB, the precursor for *anteiso-*BCFAs, is capable for supporting robust growth at low temperatures (22°C). At 30°C and 37°C, growth was comparable in the presence of either MB or IB, while no growth was observed in the absence of precursors, demonstrating that a high BCFA level is essential for growth under these conditions. At 45°C, Δ*bkd* could grow at low dilutions even in the absence of BCFA precursors (Figures 1C and S1C). For these reasons, we chose growth with IB at 37°C, as the reference condition for *B. subtilis* Δ*bkd*.

### Depletion of UFAs in *E. coli*

In *E. coli*, membrane fluidity is modulated by UFAs (Parsons & Rock, 2013) (Figure S2A). While synthesis of UFAs is essential for *E. coli* growth, a temperature-sensitive *fabF fabA*(Ts) mutant (Table S1), in which a shift to non-permissive growth temperatures leads to UFA depletion, has been isolated (Cronan & Gelmann, 1973). DNA sequencing of *fabF* and *fabA* from this rather old isolate revealed that FabF (β-ketoacyl-ACP synthase II) is non-functional due to S291N and G262S substitutions. FabA (β-hydroxyacyl-ACP-dehydratase/isomerase) carries a single G101D substitution (Rock *et al*, 1996). Based on the structure of the FabA head-to-tail homodimer (Nguyen *et al*, 2014), G101 is positioned at the border of the dimerisation interface. Consequently, the G101D substitution could plausibly cause thermosensitivity by destabilising the essential dimer structure of FabA at elevated temperatures, thereby provoking the gradual depletion of UFA (Cronan & Gelmann, 1973). Thus, this strain provides the experimental tool to control the UFA/SFA balance in growing cells. Throughout the text, the strain is labelled “*fabA*(Ts)” for simplicity.

While the growth of *fabA*(Ts) at 30°C is comparable to *E. coli* Y-Mel used as WT, transfer to non-permissive temperatures such as 37°C only supported growth for about 120 min, followed by growth arrest and onset of cell lysis (Figure 1D). Corresponding fatty acid analyses confirmed a strong, temperature-dependent decrease in UFAs (Figures 1E and S2). In agreement with Cronan & Gelmann (1973), a minimal amount of 10-15% UFAs appeared to be essential to support growth (Figures 1E and S2B). In comparison, WT cells showed only minor, temperature-dependent changes in fatty acid composition caused by homeoviscous adaptation towards increased SFA content at higher temperatures (Figures 1E and S2C).

### Reduced membrane fluidity in cells depleted of UFAs or BCFAs

To confirm that changes in fatty acid composition translate to shifts in *in vivo* membrane fluidity, we monitored changes in steady-state fluorescence anisotropy of 1,6-<u>d</u>i<u>p</u>henyl-1,3,5-<u>h</u>exatriene (DPH), the rotational freedom of which is sensitive to acyl chain disorder and, thus, indirectly to the fluidity of lipid bilayers (Lentz, 1993). DPH anisotropy measurements with *B. subtilis* Δ*bkd* revealed the highest membrane fluidity for cells with the highest *anteiso-*BCFA content (Figure 2A). Cells with high *iso-*BCFA content exhibited membrane fluidity levels slightly lower than those found for WT. These results confirm that *anteiso-*BCFAs promote higher membrane fluidity than the corresponding *iso-*forms *in vivo*; a difference previously based on *in vitro* evidence only (Lewis *et al*, 1987). The changes observed upon depletion of BCFAs, which is accompanied by accumulation of SFAs, were more drastic and resulted in a gradual reduction of membrane fluidity, ultimately leading to growth arrest (Figures 2A and S1B).

**Figure 2.**
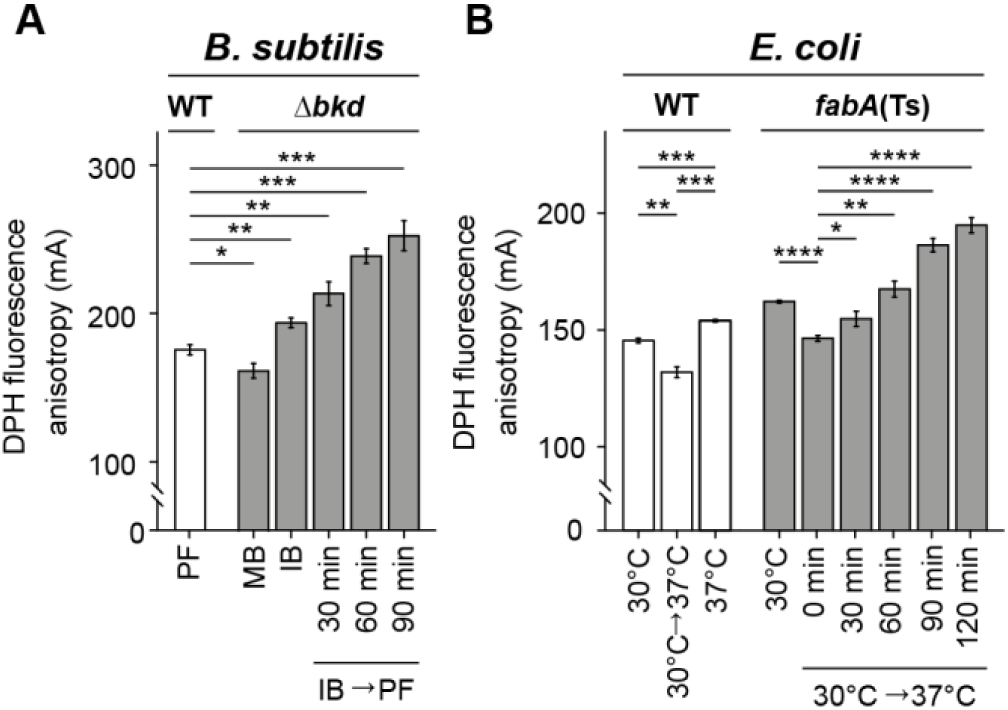
Reduced membrane fluidity in cells depleted for UFAs or BCFAs. (**A**) DPH anisotropy of *B. subtilis* WT or <u>Δ</u>*bkd* cells supplemented either with MB, IB or depleted for precursor (IB→PF) for the times indicated. High DPH anisotropy indicates low membrane fluidity. (**B**) DPH anisotropy of *E. coli* WT cells grown steady state at 30°C, 37°C or shifted from 30°C to 37°C followed by immediate measurement. In addition, DPH anisotropy of *fabA*(Ts) cells grown steady state at 30°C or shifted from 30°C to 37°C followed by measurement at the times indicated. **Data information**: (**A, B**) The experiments are representative of three independent repeats. The histograms depict means and SD of technical triplicates, together with P values of an unpaired, two-sided t-test. Significance was assumed with **** p < 0.0001, *** p < 0.001, ** p < 0.01, * p < 0.05, n.s., not significant. Strains used: (**A**) *B. subtilis* 168, HS527; (**B**) *E. coli* Y-Mel, UC1098.

DPH anisotropy measurements conducted with *E. coli* followed a similar trend (Figure 2B). Both *E. coli* WT and *fabA*(Ts) cells grown at 30°C exhibited an expected, immediate increase in membrane fluidity upon a shift to 37°C; a phenomenon that in WT cells is overtime counteracted by homeoviscous adaptation restoring membrane fluidity close to pre-shift levels. However, in *fabA*(Ts) continued growth at 37°C resulted in a gradual increase in DPH anisotropy, thus confirming a substantial reduction of membrane fluidity.

In conclusion, the established fatty acid depletion procedures allow membrane fluidity to be controllably lowered to a point incapable of supporting growth in both organisms. In the following chapters, we use this approach to analyse which cellular processes are impaired by inadequate levels of membrane fluidity.

### Consequences of low membrane fluidity on membrane diffusion barrier function

The prevalence of adaptive mechanisms maintaining membrane fluidity (Hazel, 1995) might indicate its importance for preserving the fundamental membrane barrier function. To analyse the consequences of too low membrane fluidity on membrane leakiness, we used the combination of two fluorescent dyes. Sytox Green is a membrane-impermeable, DNA-intercalating dye used to assess the integrity of bacterial plasma membranes in terms of permeability (Roth *et al*, 1997). DiSC_3_(5), a voltage-sensitive dye accumulating in cells with high membrane potential (te Winkel *et al*, 2016), indicates changes in either membrane ion conductivity or respiration.

Growing *B. subtilis* Δ*bkd* cells, irrespectively of the supplied BCFA precursor, exhibited DiSC_3_(5) fluorescence signals comparable to those observed for WT (Figures 3A, 3B and S3A-C). This indicates that the corresponding changes in the membrane fatty acid composition and fluidity had surprisingly little impact on membrane potential. In contrast, depletion of BCFAs triggered a gradual membrane depolarisation that was, in a mild form, already detectable after 30 min. A complete membrane depolarisation was observed after 90 min coinciding with growth arrest (Figure S1B). However, membranes remained impermeable for Sytox Green (Figure 3A) demonstrating that the gradual membrane depolarisation was not caused by a simple disruption of membrane continuity. In contrast, even the severely BCFA-depleted membranes were fully capable of forming a continuous, tight diffusion barrier.

**Figure 3.**
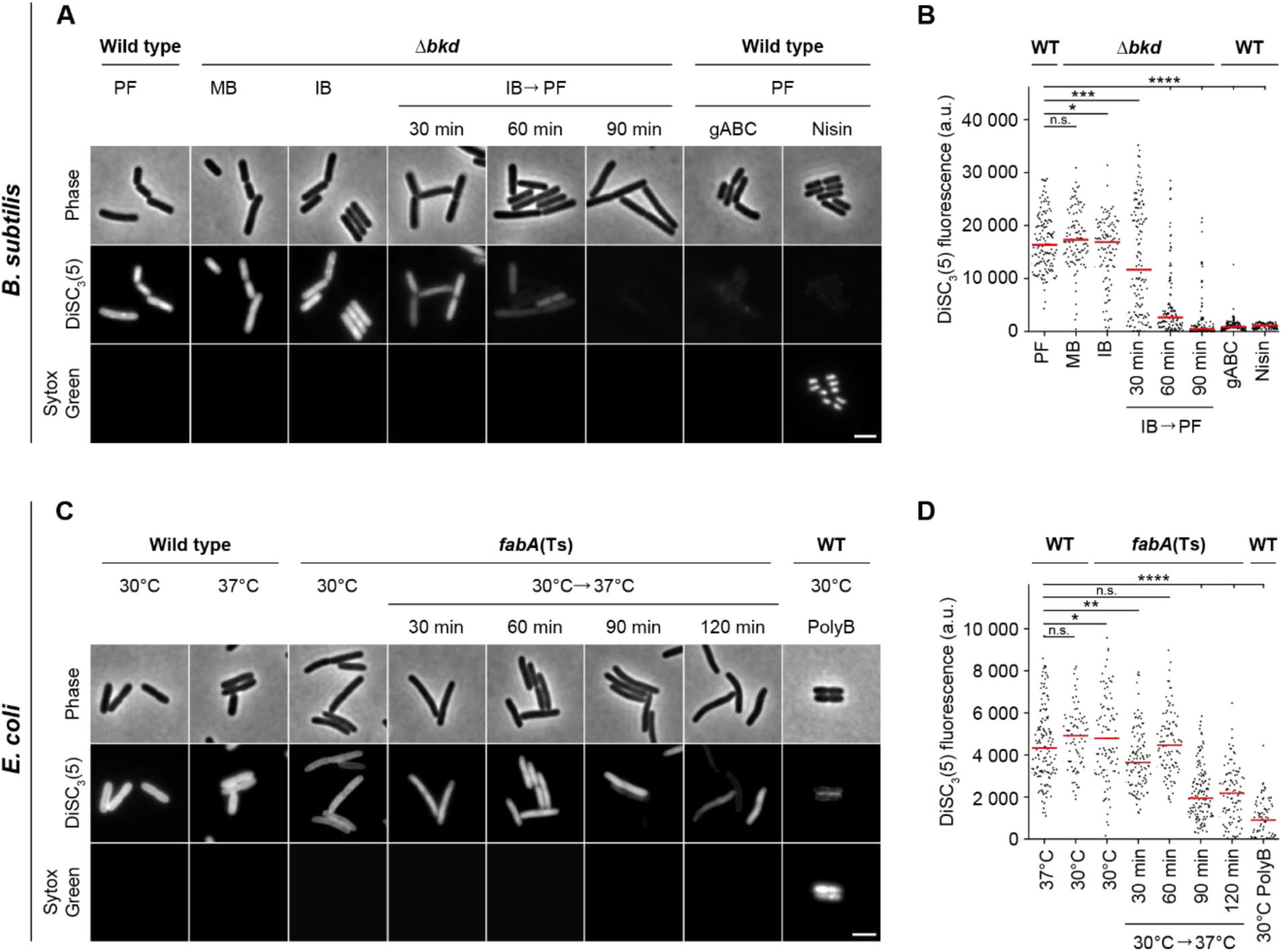
Consequences of low membrane fluidity on membrane diffusion barrier function. (**A**) Images of *B. subtilis* WT and Δ*bkd* cells co-labelled with the membrane potential-sensitive dye DISC_3_(5) and the membrane permeability indicator Sytox Green. Membrane properties were assessed for Δ*bkd* cells grown in the presence of MB, IB or washed precursor-free (IB→PF) for the times indicated. As controls, WT cells were measured in the presence of depolarising antimicrobial peptide gramicidin ABC (gABC) or pore-forming lantibiotic Nisin. For cross-correlation between membrane depolarisation and membrane permeabilisation, see Figures S3A-C. (**B**) Quantification of DISC_3_(5) fluorescence for cells (n = 100-142) depicted in panel A. Median represented by red line. (**C**) Images of *E. coli* WT and *fabA*(Ts) cells co-labelled with the same indicator dyes as in panel A. Membrane properties were assessed for *fabA*(Ts) at 30°C and upon transfer to non-permissive 37°C for the times indicated. As controls, WT cells were incubated with the pore-forming antibiotic Polymyxin B (PolyB). For cross-correlation between membrane depolarisation and membrane permeabilisation, see Figures S3D and S3E. The integrity of the diffusion barrier function was additionally studied via ONPG permeability in a Δ*lacY* background (see Figure S4). (**D**) Quantification of DISC_3_(5) fluorescence for cells (n = 76-141) depicted in panel **C**. Median represented by red line. **Data information**: (**A**-**D**) The experiments are representative of three independent repeats. (**B, D**) Red lines indicate the median. P values represent the results of unpaired, two-sided t-tests. Significance was assumed with **** p < 0.0001, *** p < 0.001, ** p < 0.01, * p < 0.05, n.s., not significant. (**A, C**) Scale bar, 3 µm. Strains used: (**A, B**) *B. subtilis* 168, HS527; (**C, D**) *E. coli* Y-Mel, UC1098.

High DiSC_3_(5) fluorescence signals and, thus, high membrane potential levels were also observed both for *E. coli* WT and *fabA*(Ts) grown at the permissive temperature of 30°C (Figures 3C, 3D, S3D and S3E), whereas depletion of fluidity-promoting UFA in *E. coli fabA*(Ts) triggered a gradual loss of membrane potential. However, compared to *B. subtilis* the loss was delayed and incomplete (Figure 3D). Again, the lack of Sytox Green staining revealed that the membranes were not impaired in their general diffusion barrier function (Figure 3C). To confirm this important finding, membrane permeability was also monitored by following LacZ–dependent hydrolysis of the chromogenic substrate ONPG (*<u>o</u>rtho*-<u>n</u>itro<u>p</u>henyl β-D-<u>g</u>alactopyranoside) in cells deficient for the uptake system LacY. While we were able to detect low level, LacZ-dependent ONPG hydrolysis in *lacY* deletion background, no difference was observed between cells with native fatty acid composition and cells strongly depleted for UFAs undergoing lipid phase separation. Hence, the results provide an independent control for the lack of membrane permeabilisation (Figure S4).

In summary, while membrane depolarisation is observed due to too low membrane fluidity in both organisms, the core permeability function of the plasma membrane is not compromised even upon conditions unable to support growth. This is consistent with a more subtle effect of low fluidity on membrane-associated biological processes maintaining ion homeostasis such as respiration.

### Consequences of low membrane fluidity on cell morphogenesis

In rod-shaped bacteria, cell growth and morphogenesis are predominantly driven by two membrane-associated multiprotein complexes, the elongasome responsible for envelope expansion and rod shape determination (Typas *et al*, 2012), and the divisome responsible for cytokinesis (Adams & Errington, 2009). The main scaffold proteins for these prominent cellular machineries are the tubulin homolog FtsZ (Adams & Errington, 2009) and the actin homolog MreB (Typas et al, 2012). To assess the functionality of these key cellular machineries, we determined the localisation of corresponding GFP fusions upon depletion of fluidity-promoting fatty acids. Furthermore, by use of GFP-fused DNA-binding protein Hu (*B. subtilis*) (Köhler & Marahiel, 1997) or DNA staining with intercalating dye DAPI (*E. coli*), we analysed the cells for potential defects in chromosome replication and segregation.

In *B. subtilis*, depletion of BCFAs had no effect on nucleoid prevalence or morphology indicating the presence of largely functional DNA replication, segregation, and compaction mechanisms (Figures 4A and S5A). While no DNA-free cells indicative for defects in DNA replication and segregation were observed in *E. coli* either, a clear de-condensation of the nucleoid was evident at later stages of UFA depletion (Figures 4B and S5B). Intriguingly, this process coincides with partial dissociation of RNA degradosome from the membrane as indicated by an increasingly cytoplasmic localisation of the key scaffold and membrane anchor protein RNase E in cells exhibiting very low fluidity (Figures S6A-D). These two processes appear to be causally linked since expression of a cytoplasmic variant of RNase E, which lacks an amphipathic helix essential for membrane binding, leads to comparable decondensation of the nucleoid (Figures S7A and S7B).

**Figure 4.**
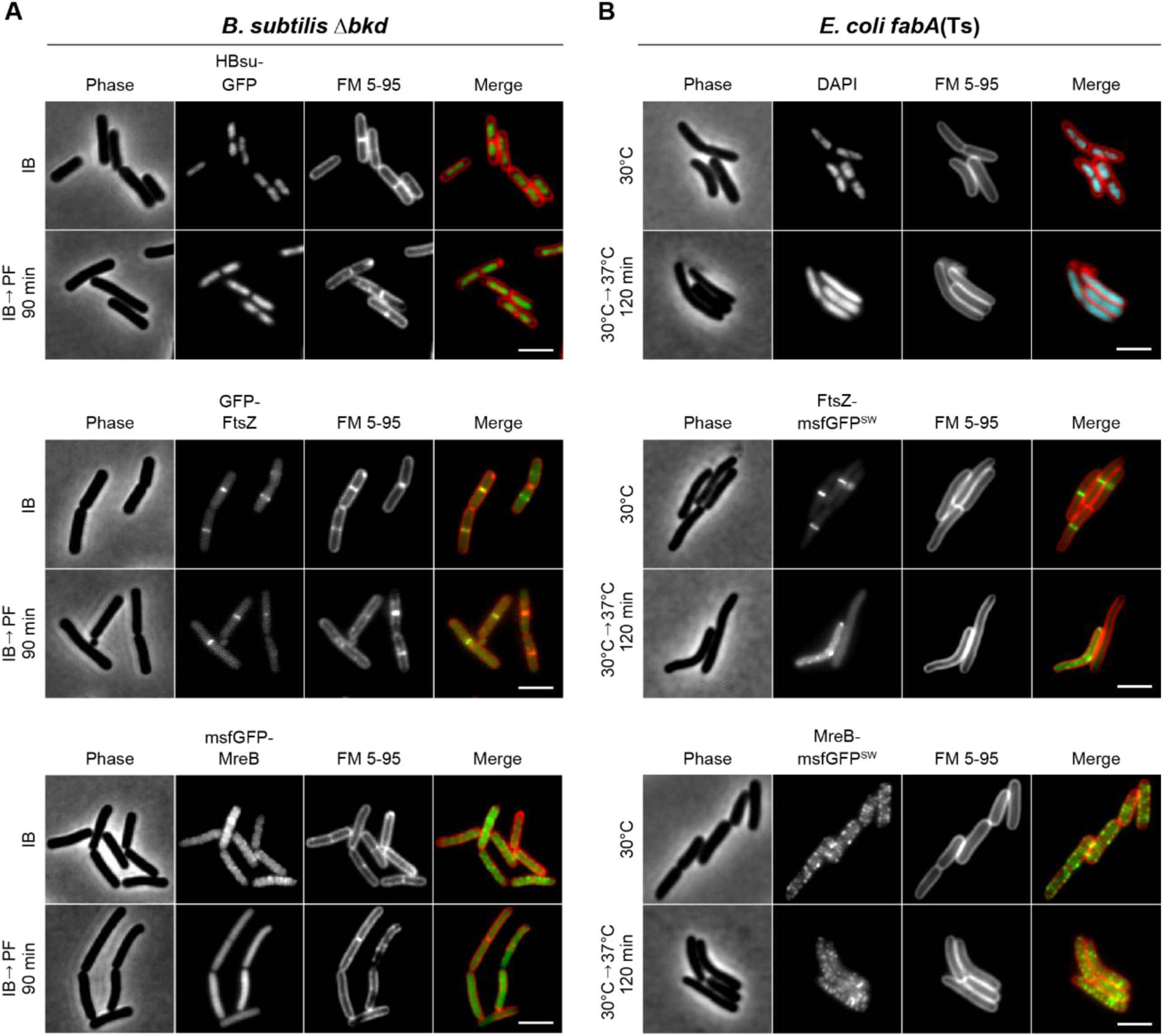
Consequences of low membrane fluidity on and cell morphogenesis. (A) Images of *B. subtilis* Δ*bkd* cells stained with membrane dye FM 5-95 and expressing GFP fusions of DNA-binding protein HBsu (top), cell division protein FtsZ (middle) or cell elongation protein MreB (bottom). Cells were grown with IB or depleted for precursors for 90 min (IB→PF). For further examples and additional time points, see Figure S5A. (B) Images of *E. coli fabA*(Ts) cells stained with FM 5-95 for the outer membrane and with DNA-intercalating dye DAPI (top), or expressing GFP <u>s</u>and<u>w</u>ich (SW) fusions to the cell division protein FtsZ (middle) and the cell elongation protein MreB (bottom), respectively. Depicted are cells grown at 30°C or with a temperature shift to 37°C for 120 min. For a more detailed view on the influence of low membrane fluidity on membrane dissociation of RNase E as well as on the cell division machinery, see Figures S6, S7 and S8. For further examples and additional time points, see Figure S5B. **Data information**: (A, B) The experiments are representative of biological triplicates. Scale bar, 3 µm. Strains used: (A) *B. subtilis* HS541, HS548, HS549; (B) *E. coli* UC1098, BHH500, BHH501.

The cell division machinery, indicated by mid-cell localisation of FtsZ, turned out to be robust towards changes in membrane fluidity in *B. subtilis*, with only a weakening of the fluorescent mid-cell signal observed upon BCFA depletion (Figures 4A and S5A). In contrast, a clear defect in divisome assembly was observed upon depletion of UFA in *E. coli* (Figures 4B and S5B). To confirm that the *E. coli* cell division machinery indeed is stressed by low membrane fluidity, we combined the UFA depletion with a deletion of cell division regulator MinC (Hu *et al*, 1999). Indeed, in the absence of MinC, the *E. coli* cell division process became the growth-limiting factor upon UFA depletion and even a slight shift in temperature from 30°C to 35°C resulted in hypersensitivity towards low fluidity and loss of viability for strain *fabA*(Ts) Δ*minC* (Figures S8A-C).

An inverse sensitivity was observed for the cell elongation machinery using localisation of MreB as proxy. In this case, depletion of BCFA triggered a complete disassembly of the MreB cytoskeleton in *B. subtilis* (Figures 4A and S5A), whereas the *E. coli* counterpart was largely unaffected from UFA depletion in terms of membrane association and filament formation (Figures 4B and S5B).

In conclusion, very low membrane fluidity incapable to support growth indeed affects membrane-associated cellular machineries responsible for bacterial growth and division. It is important to note, however, that the changes in membrane fluidity required to disturb cell morphogenesis are rather extreme and go well beyond those observed upon normal changes in growth temperature.

### Consequences of low membrane fluidity on membrane homogeneity

In the microscopic experiments described above (Figures 4A and 4B), cells were stained with FM 5-95. This hydrophobic fluorescent dye allows visualization of the plasma membrane of *B. subtilis* (Sharp & Pogliano, 1999), or the outer membrane in the case of *E. coli* (Pilizota & Shaevitz, 2012). While no changes in FM 5-95 staining were observed in *E. coli* cells, the smooth staining observed in Δ*bkd* cells supplemented with IB transitioned into a distinctly irregular pattern upon BCFA depletion (Figures 4A and S5A). This suggests that the more homogeneous membrane, present under normal growth conditions, segregates into areas with different local physicochemical properties upon low membrane fluidity. Intriguingly, the development of membrane irregularities coincides with growth arrest (Figure 5A and Movie S1). It is worth emphasising, however, that the changes in lipid fatty acid composition needed to trigger this response are rather extreme compared to changes observed upon normal homeoviscous adaptation (Figures S9A and S9B).

**Figure 5.**
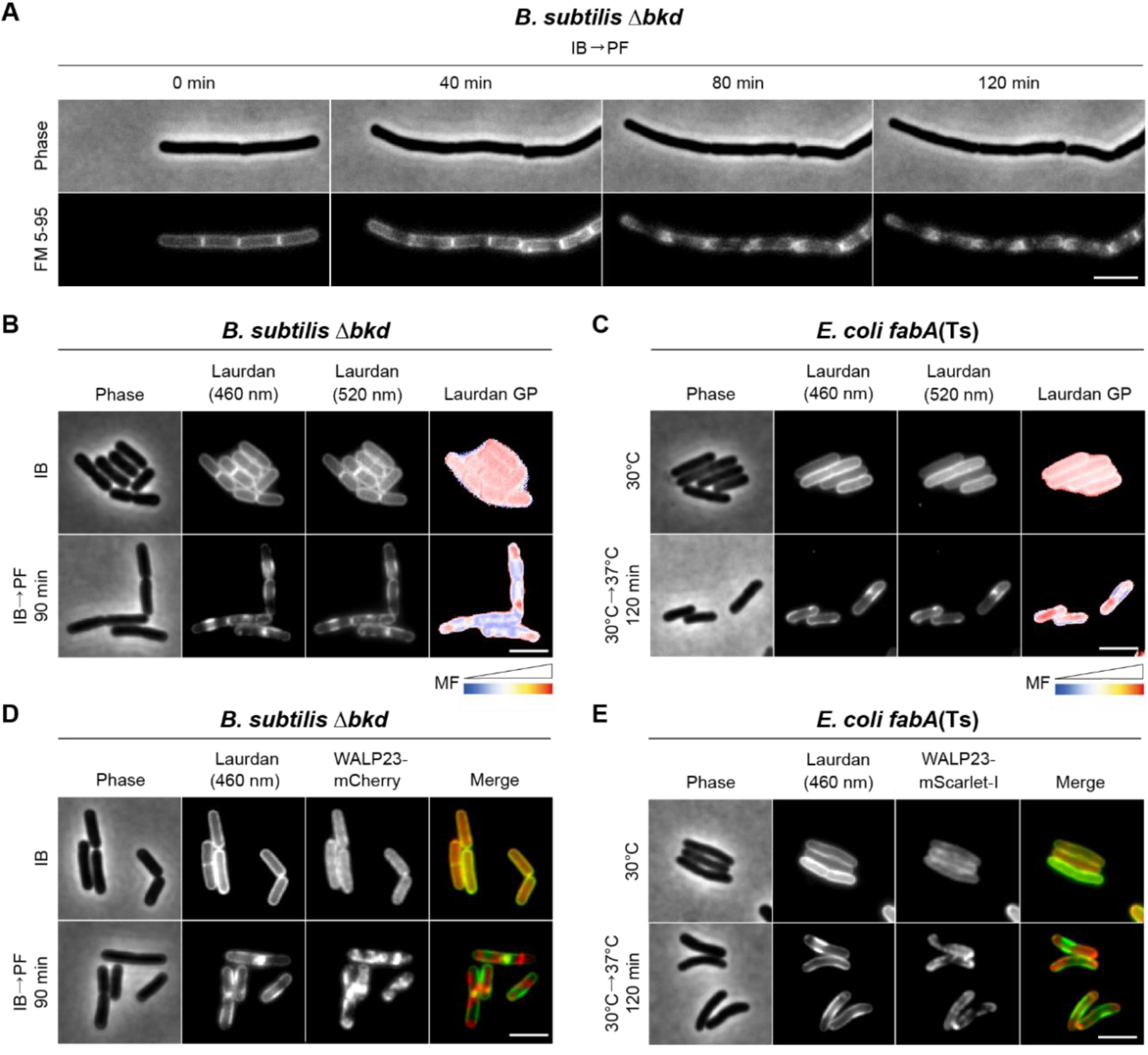
Consequences of low membrane fluidity on membrane homogeneity. (**A**) Time lapse images of *B. subtilis* Δ*bkd* cells stained with FM 5-95 and grown in PF medium. See Movie S1 for full time lapse. Figure S9 shows the changes observed in relation to natural homeoviscous adaptation. (**B**) Images of *B. subtilis* Δ*bkd* cells grown with IB or without precursor (IB→PF). Cells were stained with membrane fluidity-sensitive dye Laurdan and imaged at 460 nm, 520 nm and as the corresponding colour-coded Laurdan GP map. (**C**) Images of *E. coli fabA*(Ts) cells grown at 30°C or shifted to non-permissive 37°C for 120 min. Cells were stained and imaged as described in panel B. For a more pronounced view on domain formation associated with local differences in membrane fluidity, see Figure S10, showing *E. coli fabA*(Ts) grown in LB instead of M9 minimal medium with glucose/casamino acids. (**D**) Images of *B. subtilis* Δ*bkd* cells grown and stained with Laurdan (imaged at 460 nm) as in panel B, but additionally expressing WALP23-mCherry. For corresponding fluorescence intensity correlation between images, see Figure S11A. Co-localisation of membrane dye FM 5-95 and transmembrane peptide WALP23 is shown in Figures S12 and S11C. (**E**) Images of *E. coli fabA*(Ts) cells grown and stained as in panel C, but additionally expressing WALP23-mScarlet-I. For corresponding fluorescence intensity correlation between images, see Figure S11B. **Data information**: (**A**-**E**) The experiments are representative of biological triplicates. Scale bar, 3 µm. MF, <u>m</u>embrane <u>f</u>luidity. Strains used: (**A, B**) *B. subtilis* HS527; (**C**) *E. coli* UC1098; (**D**) *B. subtilis* HS547; (**E**) *E. coli* UC1098/pBH501.

*In vitro*, lipid mixtures of low fluidity undergo phase transition into a more tightly packed gel state. We speculated that the lipid de-mixing observed in *B. subtilis* (Figures 4A and 5A) could therefore represent large-scale lipid phase separation between gel and liquid-disordered phases. Therefore, we analysed the local membrane fluidity of both *B. subtilis* and *E. coli* with the fluidity-sensitive membrane dye Laurdan. Laurdan exhibits shifts in its fluorescence emission spectrum that depends on lipid packing-linked penetration of H_2_O to the membrane interior and, thus, the immediate environment surrounding the fluorophore. The interconnected nature of lipid packing and membrane fluidity allows local membrane fluidity to be estimated as Laurdan <u>g</u>eneralized <u>p</u>olarization (GP) (Parasassi *et al*, 1990; Wenzel *et al*, 2018). Indeed, when *B. subtilis* and *E. coli* cells were depleted for fluidity-promoting fatty acids, domain formation associated with both differential staining (Laurdan fluorescence intensity) and differences in local fluidity (Laurdan GP) was observed (Figures 5B, 5C and S10). To verify these findings with an independent dye-free method, we used cells expressing helical transmembrane peptide WALP23, which preferentially accumulate in liquid-disordered membrane areas (Schäfer *et al*, 2011; Scheinpflug *et al*, 2017b). Indeed, co-labelling of cells with Laurdan clearly demonstrated that the observed lipid phase separation results in segregation of WALP23 in membrane areas of low Laurdan fluorescence (Figures 5D, 5E, S11A and S11B). At last, co-staining with FM 5-95 demonstrated that FM 5-95 and WALP23 share the same preference for higher fluidity areas in de-mixed membranes (Figures S11C and S12).

In conclusion, by exhibiting de-mixing into distinct areas of high and low membrane fluidity, respectively, the observed *in vivo* domain formation shares the core characteristic of lipid phase separation between fluid and gel state membranes (Baumgart *et al*, 2007; Domanski *et al*, 2012; Mostofian *et al*, 2019).

### Segregation of membrane proteins into fluid domains of phase-separated plasma membranes

As indicated by WALP23 (Figures 5D and 5E), the observed lipid phase separation might have broader consequences on membrane protein localisation. To test this, we focused on *E. coli* for two reasons. Firstly, the membrane depolarisation caused by low membrane fluidity, which itself can affect membrane protein localisation (Strahl & Hamoen, 2010), is less extensive in *E. coli* (Figure 3). Secondly, *E. coli* does not exhibit delocalisation of MreB (Figures 4B and S5B), which we have previously shown to induce membrane protein clustering (Strahl *et al*, 2014). As a model protein of choice, we focused on ATP synthase (F_O_F_1_), an abundant polytopic membrane protein complex (Junge & Nelson, 2015).

To visualise the localisation of F_O_F_1_, fluorescent protein was C-terminally fused to membrane-integral F_O_-*a* yielding a stable and active enzyme (Figures S13A and S13B). Upon UFA depletion, F_O_F_1_ showed clear segregation behaviour (Figure 6A and Movie S2). When co-expressed, F_O_F_1_ and WALP23 showed clear co-segregation into the fluid areas of phase-separated membranes (Figures 6B and S11D). This property was also confirmed by co-staining with Laurdan showing an anti-correlation of the fluorescent signals (Figures 6C and S11E). In agreement with the smooth outer membrane staining using FM 5-95 (Figures 4B and S5B), no segregation was observed for the major outer membrane protein OmpA. Co-labelling of the inner and outer membrane with fluorescent proteins analysed by super resolution structured illumination microscopy demonstrated that the inner membrane marker F_O_F_1_ *a*-mNG showed segregation into the fluid areas at non-permissive 37°C, while the pattern of the outer membrane marker OmpA-mCherry remains homogeneous, thus supporting the view that the outer membrane does not participate in the phase separation process (Figures S14A-D).

**Figure 6.**
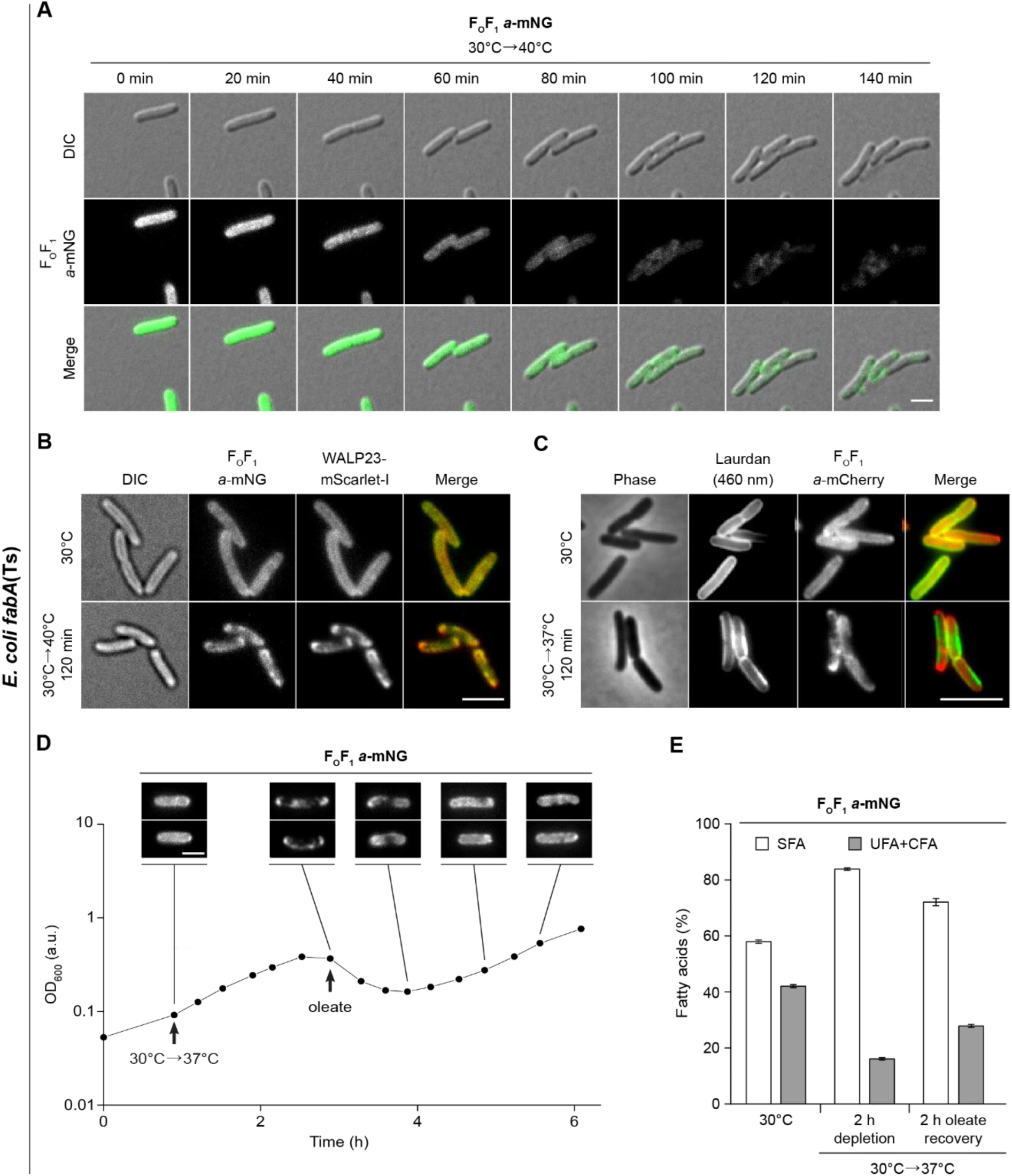
Partitioning of membrane proteins into fluid domains of phase-separated plasma membranes. (**A**) Time lapse images of *E. coli fabA*(Ts) cells expressing F_O_F_1_ *a*-mNG. For corresponding controls, see Figure S13. Cells were grown at 30°C and shifted from 30°C to non-permissive 37°C. See Movie S2 for full time lapse. (**B**) Images of *E. coli fabA*(Ts) co-expressing WALP23-mScarlet-I and F_O_F_1_ *a*-mNG grown at permissive 30°C or shifted to non-permissive 40°C. For fluorescence intensity correlations, see Figure S11D. For corresponding images of *E. coli fabA*(Ts) cells co-expressing inner membrane marker F_O_F_1_ *a*-mNG and outer membrane marker OmpA-mCherry, see Figure S14. (**C**) Images of F_O_F_1_ *a*-mCherry-expressing *E. coli fabA*(Ts) grown at 30°C or shifted to non-permissive 37°C and stained with Laurdan. For fluorescence intensity correlations, see Figure S11E. (**D**) Growth behaviour of F_O_F_1_ *a*-mNG expressing *E. coli fabA*(Ts) after shift from 30°C to non-permissive 37°C and upon recovery through exogenous supplementation with UFA oleate (*cis*-Δ9-C18:1). The corresponding reversible segregation of F_O_F_1_ *a*-mNG is shown above the growth curve. For further controls, see Figure S15A. For a comparable fluorescence time lapse analysis of mNG-labelled glucose permease PtsG, see Figure S16. (**E**) Fatty acid composition of *E. coli fabA*(Ts) cells (same cell batch as in D) upon growth at 30°C, upon depletion of UFA by incubation at 37°C for 120 min and upon recovery by oleate supplementation for additional 120 min. For detailed analyses, see Figures S15B-D. **Data information**: (**A**-**D**) The experiments are representative of biological triplicates. (**E**) The histogram depicts mean and SD of biological triplicates. DIC, <u>d</u>ifferential <u>i</u>nterference <u>c</u>ontrast. Strains used: (**A, D, E**) *E. coli* MG4; (**B**), *E. coli* MG4/pBH501; (**C**), *E. coli* LF6.red.

Depletion of UFAs had no substantial influence on the DCCD-sensitive ATPase activity of F_O_F_1_ (Figure S13B), thus arguing that high viscosity of the surrounding lipids does not significantly hinder F_O_F_1_ in its rotation-based catalytic cycle (Junge & Nelson, 2015). In a wider context, this indicates that the remaining fluid phase, to which F_O_F_1_ partitions upon UFA depletion, can retain robust bioactive properties. Motivated by this finding, we analysed whether the lipid phase separation and the associated growth arrest is reversible. Indeed, when severely UFA-depleted *fabA*(Ts) cells exhibiting both growth arrest and lipid phase separation were exogenously supplied with oleate (*cis*-Δ9-C18:1), incorporation of oleate into phospholipids, together with growth recovery and restoration of the dispersed distribution of F_O_F_1_ was observed (Figures 6D, 6E and S15A-D). Comparable experiments performed with fluorescently labelled glucose permease (PtsG-mNG) confirmed that the observed protein segregation and its recovery (Figures S16A-C) is not unique for ATP synthase.

In summary, our results demonstrate that lipid phase separation occurring under conditions of low membrane fluidity has a profound effect on membrane protein distribution, triggering segregation of integral membrane proteins into the remaining liquid-disordered phase areas.

### Restricted diffusion of membrane proteins in UFA-depleted *E. coli* membranes

To analyse the consequences of UFA depletion on protein diffusion and, thus, membrane fluidity directly, we followed mNG-labelled F_O_F_1_ (F_O_F_1_ *a*-mNG) by *in vivo* single molecule tracking. Consistent with the lack of a specific localisation pattern, F_O_F_1_ *a*-mNG complexes exhibited free diffusion within the plasma membrane plane of *E. coli* WT cells (Figures 7A, S17 and Movie S3). The observed lateral mobilities and jump sizes were largely independent of the growth temperature (Figures 7B and S17), as expected for cells with active homeoviscous adaptation mechanisms. F_O_F_1_ *a*-mNG expressed in *fabA*(Ts) cells at 30°C also showed unrestricted lateral mobility comparable to WT. Under conditions of UFA depletion (33-40°C), however, a gradual, temperature-dependent reduction of lateral displacement and median jump sizes was observed (Figures 7A, 7B, S17 and Movie S4). This is consistent with a local confinement of F_O_F_1_ caused by co-occurring lipid phase separation (compare Figures 6A-C). Calculation of apparent lateral diffusion coefficients (D_app_) revealed that the lateral mobility was reduced up to 9-fold (Figure 7C).

**Figure 7.**
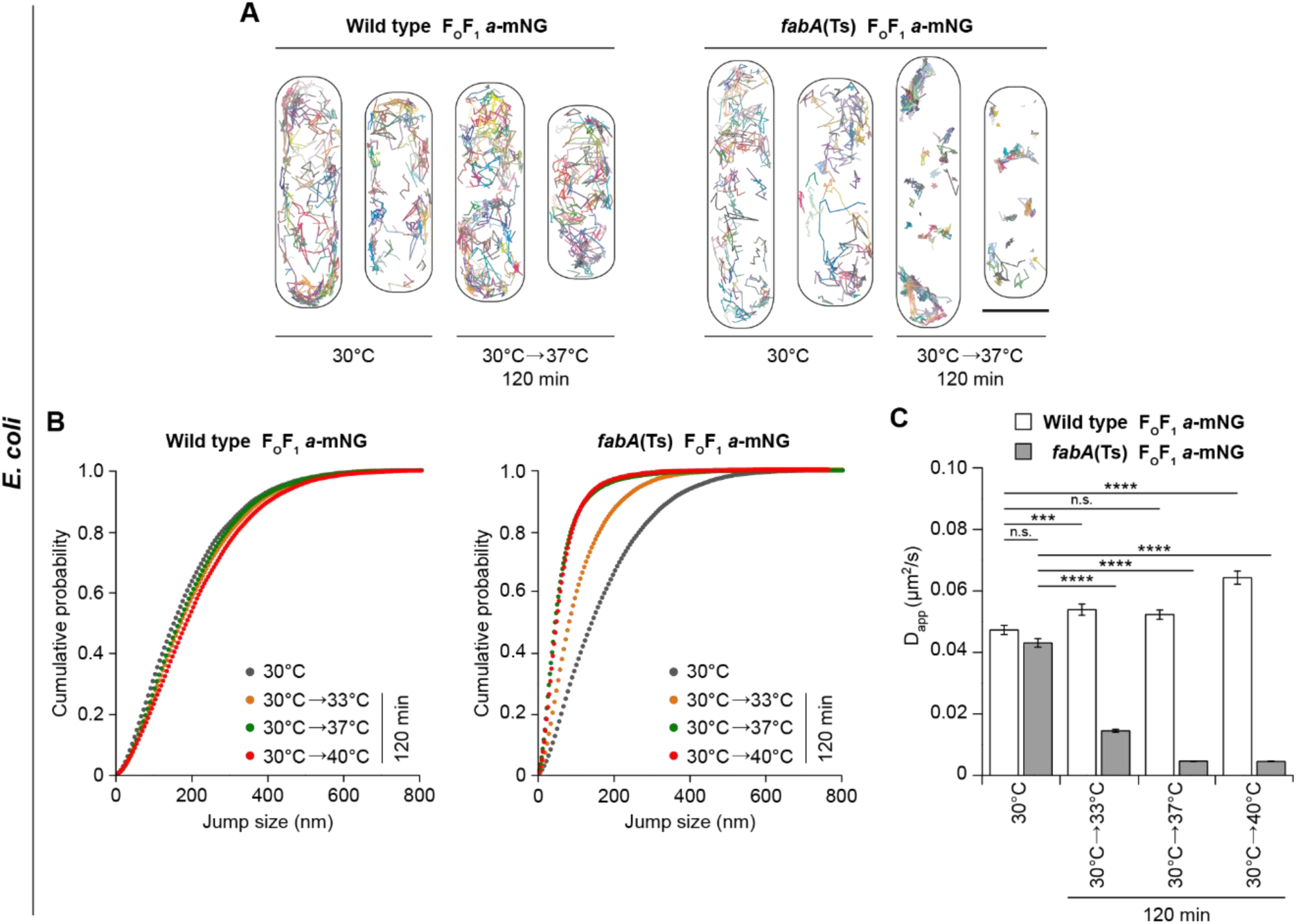
Restricted diffusion of membrane proteins in UFA-depleted *E. coli* membranes. (**A**) Representative trajectory maps of individual F_O_F_1_ *a*-mNG molecules in *E. coli* WT and *fabA*(Ts) cells grown at 30°C and upon shift to non-permissive 37°C for 120 min. See Movie S3 (WT) and Movie S4 (*fabA*(Ts)) for accumulating trajectory maps of F_O_F_1_ *a*-mNG molecules. (**B**) Cumulative probability plots of F_O_F_1_ *a*-mNG jump sizes in *E. coli* WT and *fabA*(Ts) cells. For an analysis of cell-to-cell homogeneity of median jump sizes of F_O_F_1_ *a*-mNG, see Figure S17. (**C**) Apparent lateral diffusion coefficients (D_app_) of F_O_F_1_ *a*-mNG analysed in panel **B**. For a comparable single molecule tracking analysis of transmembrane peptide WALP23-mNG, see Figure S18. A detailed analysis of F_O_F_1_ *a*-mNG in an osmotically stabilized *fabB1*5(Ts) mutant (2% KCl) at non-permissive 40°C is shown in Figures S19 and S20. **Data information**: (**A**) Representative for 3-5 biological replicates. (**B**) Trajectories with ≥5 consecutive frames for each growth condition and strain were pooled from 3-5 biological replicates (n=2345-4468). (**C**) Median and SD from 3-5 biological replicates, together with P values of a two-sided Wilcoxon rank sum test. Significance was assumed with **** p < 0.0001, *** p < 0.001, ** p < 0.01, * p < 0.05, n.s., not significant. (**A**) Scale bar, 1µm. Strains used: (**A**-**C**), *E. coli* MG1, MG4.

In comparison, single molecule tracking of mNG-labelled WALP23, expressed in *fabA*(Ts) cells at 30°C also exhibited rapid, unconfined diffusion. As expected, due to its single transmembrane helix (Lucena *et al*, 2018; Ramadurai *et al*, 2009), the median jump sizes and D_app_ of WALP23 (Figures S18A-C) were higher than those observed for F_O_F_1_. Upon lipid phase separation caused by UFA depletion, WALP23 exhibited confined mobility comparable to that observed for F_O_F_1_, again supporting the notion that both proteins co-segregate into the remaining fluid phase (Figures 6B and S11D).

Osmotic stabilisation has been described for some UFA auxotrophic *E. coli* strains (Akamatsu, 1974; Broekman & Steenbakkers, 1973; 1974). Whereas osmotic stabilisation does not restore the viability of *fabA*(Ts) strain UC1098 in non-permissive temperatures (Akamatsu, 1974), a strain carrying a *fabB15*(Ts) mutation is able to grow at non-permissive 40°C as long as an osmotic stabilizer (e.g. 2% KCl) is present (Akamatsu, 1974) (Figure S19A). Fluorescence microscopy of *fabB15(Ts)* cells expressing F_O_F_1_ *a*-mNG revealed that protein partitioning observed in the presence of KCl is detectable but less severe (Figure S19B), while single molecule tracking of F_O_F_1_ *a*-mNG showed a lesser reduction of lateral diffusion compared to *fabA*(Ts) (Figure S20A). Crucially, fatty acid analyses (Figures S20B-D) showed that depletion of UFA is significantly less pronounced in this strain upon osmotic stabilisation with KCl. Hence, osmotic stabilisation of this strain does not rescue the cells from low UFA content but rather acts by partially restoring UFA synthesis.

In summary, depletion of UFA in *E. coli* results in a strong reduction of membrane fluidity that severely restricts lateral diffusion of membrane proteins (summarised in Figure 8). As a complementary, dye-independent method, the tracking experiments also confirm the lipid phase separation phenomenon, resulting in integral membrane proteins segregated and confined into the remaining fluid membrane areas.

**Figure 8.**
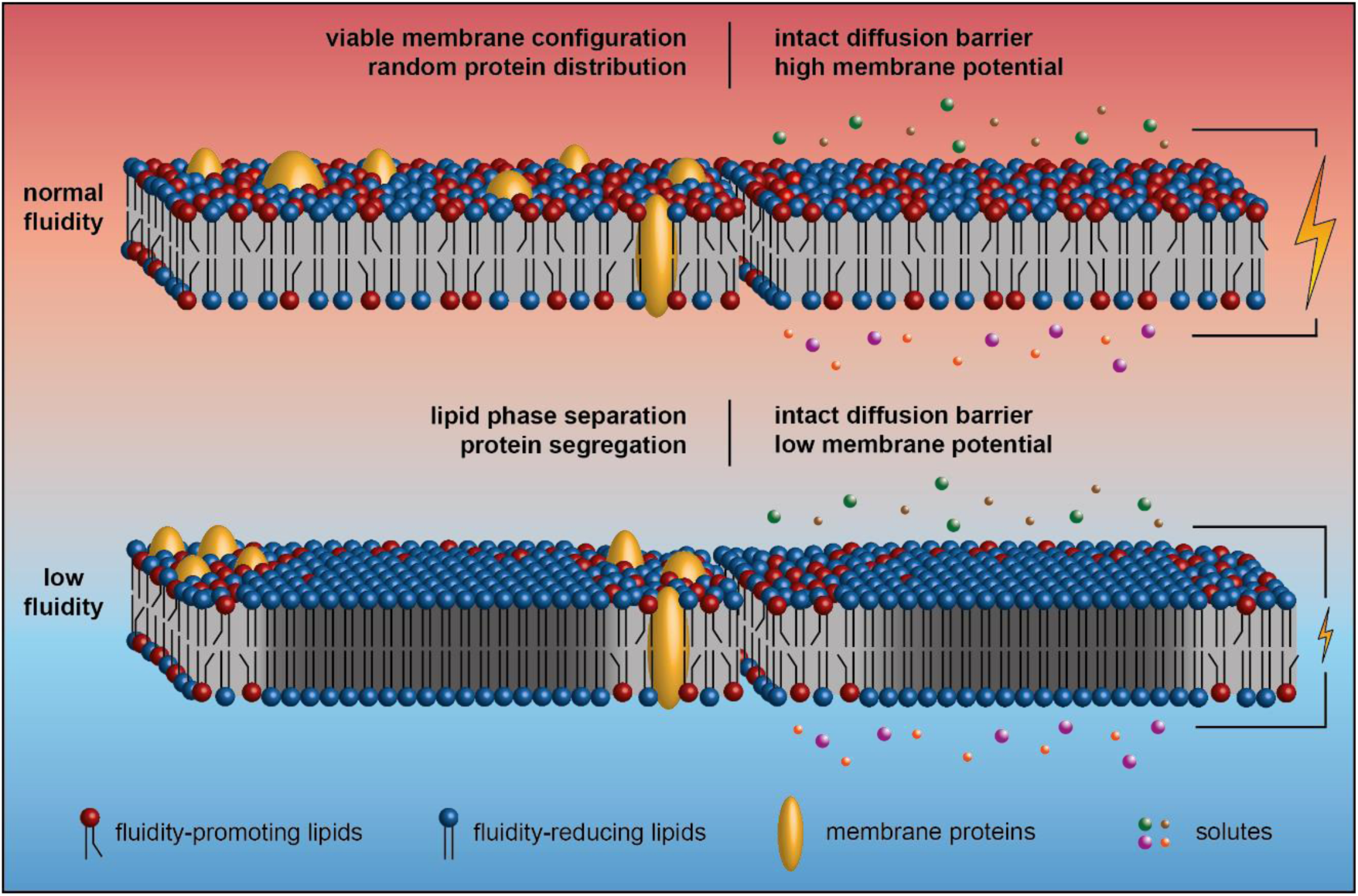
Low membrane fluidity triggers large-scale gel-liquid lipid phase separation *in vivo*. The cartoon illustrates how too low membrane fluidity triggers large-scale lipid phase separation associated with both segregation and confined diffusion of membrane-integral proteins (left part). Too low membrane fluidity also results in dissipation of membrane potential likely due to a reduced electron transport chain activity (Budin *et al*, 2018). The core diffusion barrier function of the membrane, however, is maintained (right part).

## Discussion

Consistent with early studies (Boudreaux *et al*, 1981; Kaneda, 1977; Cronan & Gelmann, 1973; Willecke & Pardee, 1971), the ability to maintain membrane fluidity through synthesis of fluidity-promoting lipid species is indeed essential for the growth of both *E. coli* and *B. subtilis*. However, the magnitude of changes in composition and fluidity which the cells can tolerate is surprising (summarised in Figure 8). While *B. subtilis* cells finely balance the ratio of *iso*- and *anteiso*-BCFA in response to changes in temperature (Klein *et al*, 1999; Suutari & Laakso, 1992), even the massive changes in the *iso*/*anteiso*-ratio obtained through precursor supplementation had no significant effect on growth behaviour. Similarly, changes in fatty acid composition and membrane fluidity needed to impair growth of *E. coli* are much more drastic than those observed as part of the normal homeoviscous adaptation upon temperature shifts (Mansilla *et al*, 2004; Marr & Ingraham, 1962; Sinensky, 1974). Consequently, while both *E. coli* and *B. subtilis* adapt their membrane composition and fluidity even upon subtle changes in temperature, the failure to do so is not associated with immediate growth-inhibitory consequences. Thus, at least for these organisms, the biological reason for the homeoviscous adaptation process is not directly evident.

A more drastic reduction of membrane fluidity, however, does have severe, growth-inhibitory consequences. Whereas low membrane fluidity associated with lipid phase separation is accompanied by a substantial increase in membrane leakiness *in vitro* (Cordeiro, 2018; Heimburg, 2007; Papahadjopoulos *et al*, 1973), our results suggest that, at least in the context of the *in vivo* plasma membranes of *B. subtilis* and *E. coli*, such an effect does not play a significant role. Even the very low fluidity membranes incapable for supporting growth are able to retain robust diffusion barrier function. Rather than indicating ion leakage, the observed gradual and partial membrane depolarisation is fully consistent with previous reports demonstrating that the membrane fluidity influences the <u>e</u>lectron <u>t</u>ransport <u>c</u>hain (ETC) both in *E. coli* and mitochondria (Budin *et al*, 2018; Torres *et al*, 2018). While the enzyme complexes of the ETC maintain their function (as we also observed for F_O_F_1_), the diffusivity of ubiquinone is reduced in membranes of low fluidity and thus controls the electron transfer rate in the ETC (Budin *et al*, 2018). Therefore, maintaining robust ETC activity may well be one of the biological reasons why a fine, homeostatic balance of membrane fluidity is important.

In addition to changes in membrane potential, we also observed severe effects on the machineries responsible for cell morphogenesis. The reduction of membrane fluidity in *B. subtilis* is associated with rapid delocalisation of MreB indicating disturbance of lateral cell wall synthesis. Conversely, in *E. coli* both the cell division machinery and the nucleoid morphology were disturbed. However, both cell division and cell wall synthesis machineries are also influenced by membrane depolarisation (Strahl & Hamoen, 2010; Strahl *et al*, 2014). It therefore remains to be determined whether the observed changes are a direct consequence of low membrane fluidity, or secondarily caused by the gradual membrane depolarisation. It is worth highlighting though, that the changes in membrane fluidity required to disturb cell morphogenesis are quite extreme and go well beyond those observed upon normal changes in growth temperature. Consequently, rather than as a sign for sensitivity, these findings provide an indication for the relative robustness of bacterial morphogenetic systems towards normally encountered changes in membrane fluidity.

The most striking phenomenon caused by severe reduction of membrane fluidity is the lipid de-mixing associated with membrane protein segregation (summarised in Figure 8). Molecular dynamic simulations with membrane models composed of lipids with SFAs and BCFAs revealed increased ordering of the lipid bilayer when the SFA content was systematically increased. In the presence of approximately 20% SFAs (compared to 5-7% in *B. subtilis* WT cells), a sharp transition representing phase separation between liquid and gel phase was observed (Mostofian *et al*, 2019). These findings are fully consistent with the *in vivo* lipid de-mixing we observe upon depletion of BCFAs in favour of SFAs accumulation in *B. subtilis*, albeit with a slightly higher SFA content needed. In *E. coli*, we observe *in vivo* lipid de-mixing upon accumulation of SFA to a maximal content of 80% (compared to about 50% in WT (Zhu *et al*, 2009)), which again is consistent with previous *in vitro* studies regarding lipid phase separation of SFA/UFA mixtures (Letellier *et al*, 1977; Morein *et al*, 1996; Suárez-Germà *et al*, 2011). For these reasons, we argue that the observed de-mixing represents lipid phase separation between liquid-disordered and gel state membranes, here occurring in intact, living cells. This phenomenon is likely comparable to that observed in eukaryotic endoplasmic reticulum, characterized by low cholesterol levels, upon metabolism of SFA C18:0 (Shen *et al*, 2017). While the phase separation phenomenon unarguably affects the plasma membranes both in Gram-positive *B. subtilis* and Gram-negative *E. coli*, we suggest that the process is limited to the plasma membrane and does not encompass the Gram-negative outer membrane. This notion is based on an even distribution of a major outer membrane protein OmpA and smooth FM 5-95 outer membrane staining in cells with phase-separated inner membranes.

The specificity of dyes to label different lipid phases is still a matter of debate. Small hydrophobic dyes may themselves alter the composition and ordering of coexisting phases, even when used in trace amounts (Veatch, 2007). However, we are convinced that (*i*) by using the combination of two chemically distinct fluorescent membrane dyes Laurdan and FM 5-95 exhibiting opposing phase preferences, (*ii*) by combining dye-based approaches with localisation of WALP23 peptide previously shown to exhibit liquid-disordered phase preference *in vivo, in vitro* and *in silico* (Ridder *et al*, 2004; Schäfer *et al*, 2011; Scheinpflug *et al*, 2017b) and (*iii*) by following the reversible phase separation through its consequences on lateral membrane protein diffusion, an approach completely independent of dyes but based on intrinsic chromosomal expression of proteins, we have exhausted the possibility that the observed phase separation is an artefact caused by the labelling techniques used.

Importantly, by demonstrating lipid liquid-gel phase separation and the associated membrane protein segregation occurring in protein-crowded, native membranes of living cells, our results are fully consistent with comparable phenomena observed in simplified *in vitro* and *in silico* model systems (Domański *et al*, 2012; Picas *et al*, 2010; Suárez-Germà *et al*, 2011; Shaw *et al*, 2021), thus providing strong, complementary *in vivo* support for the general validity of the respective membrane models.

It is perhaps not surprising that the observed lipid phase separation coincides with growth arrest. Transmembrane segments of integral membrane proteins are embedded within the hydrophobic interior of lipid bilayers. Consequently, lipid bilayer thickness, which acutely changes with membrane fluidity and phase, is both important for membrane protein activity, and drives partitioning of proteins between different phases (Lee, 2004; Lenaz, 1987; Lorent *et al*, 2017; Nickels *et al*, 2019). Peripheral membrane proteins, which establish membrane association through bilayer-intercalating domains such as amphipathic helices, in turn rely on sufficiently low packing density/high fluidity for efficient membrane association (Bigay & Antonny, 2012; Drin & Antonny, 2010; Strahl & Errington, 2017). At last, the severe restriction of lateral diffusion caused by phase separation is likely interfering with localisation and activity of many membrane-associated cellular processes relying on diffusion and capture mechanism (Rudner *et al*, 2002). In conclusion and as suggested earlier based on indirect evidence (Burns *et al*, 2017; Drobnis *et al*, 1993; Ghetler *et al*, 2005), we argue that it is indeed the lipid phase separation process and the formation of gel phase areas that determines the lower end of membrane fluidity capable of supporting viable cell functions.

## Materials and Methods

### *E. coli* strains and growth conditions

*E. coli* strains (Appendix Table S1) were grown in M9 minimal medium composed of Na_2_HPO_4_•2H_2_O (0.85% w/v), KH_2_PO_4_ (0.3% w/v), NaCl (0.3% w/v), NH_4_Cl (0.05% w/v), MgSO_4_•7H_2_O (0.25% w/v), CaCl_2_•2H_2_O (0.015% w/v) and supplemented with thiamine (0.01% w/v), casamino acids (0.1% w/v) and glucose (0.4% w/v) at 30°C, unless stated otherwise. For induction of the temperature-sensitive phenotype *fabA*(Ts), pre-cultures of *E. coli* strain UC1098 and its derivatives were diluted from an overnight culture grown at 30°C to OD_600_ of 0.025 and grown to OD_600_ of 0.5 at 30°C, and again diluted to OD_600_ of 0.025 in pre-warmed, fresh medium for complete removal of cells in the stationary growth phase. At OD_600_ of 0.1-0.2, cells were transferred for 120 min to growth temperatures of 33°C, 37°C or 40°C as indicated. WT strains were handled accordingly. For induction of the thermosensitive phenotype of *fabB15*(Ts) cells were grown essentially as described (Akamatsu, 1974). Precultures of *E. coli* strain BHH87 were diluted from an overnight culture, grown at 30°C in the presence or absence of 2% (w/v) KCl, to OD_600_ of 0.025 and grown at permissive 30°C to OD_600_ of 0.4-0.5 or grown at non-permissive 40°C for 180 min, each with or without 2% KCl. The corresponding WT strain was handled accordingly. For recovery from UFA depletion and corresponding phase separation, UC1098 derivatives were subsequently supplemented with potassium oleate (100 µg/ml; Sigma Aldrich) dissolved in Brij^®^58 (0.1% w/v; Sigma Aldrich). Fluorescently labelled ATP synthase (F_O_F_1_ *a*-mNG or F_O_F_1_ *a*-mCherry; C-terminal fusion to F_O_-*a*), fluorescently labelled glucose permease (PtsG-mNG; C-terminal fusion) and msfGFP sandwich fusions of FtsZ and MreB were expressed from their own locus under control of their native promoter. Fluorescently labelled WALP23 (WALP23-mScarlet-I or WALP23-mNG; amino acid sequence: AWW(LA)_8_LWWA) was expressed plasmid-encoded under control of the *B. subtilis Pxyl* promoter (Appendix Table S3) resulting in constitutive expression in *E. coli*. Fluorescently labelled RNase E-YFP (C-terminal fusion) was expressed plasmid-encoded under control of its own promoter. RNase E and ΔAH-RNase E (RNase E lacking the membrane-binding amphipathic helix) were expressed plasmid-encoded under control of the *ParaBAD*-inducible promoter. Details for strain construction can be found in the Appendix.

### *B. subtilis* strains and growth conditions

For strain construction, *B. subtilis* (Appendix Table S1) was grown either in LB (lysogeny broth), Nutrient Broth or Nutrient Agar (Oxoid). If necessary, these media were supplemented with either <u>i</u>so<u>b</u>utyric acid (IB) (100 µM; Sigma Aldrich) or 2-<u>m</u>ethyl<u>b</u>utyric acid (MB) (100 µM; Sigma Aldrich). All other experiments were carried out with fortified Spizizen minimal medium composed of (NH_4_)_2_SO_4_ (0.2% w/v), K_2_HPO_4_ (1.4% w/v), KH_2_PO_4_ (0.6% w/v) Na_3_-citrate•2H_2_O (0.1% w/v), MgSO_4_ (0.09% w/v), ferric ammonium citrate (1.1 µg/ml) supplemented with glucose (0.96% w/v), L-tryptophan (20 µg/ml), and casamino acids (0.02% w/v). In our hands, the precursor isovaleric acid, which is the primer for the synthesis of *iso-*C15:0 and *iso-*C17:0, neither supported growth nor resulted in synthesis of the expected *iso-*BCFAs, thus implying that this precursor cannot be supplied exogenously in the *B. subtilis* 168 strain background. All cultures were inoculated by 1:100 dilution of an LB overnight culture supplemented with the corresponding precursor. Depletion of BCFAs was carried out for cells initially grown in the presence of IB (100 µM), followed by washing, pelleting and resuspension in pre-warmed, <u>p</u>recursor-<u>f</u>ree medium (PF). Unless stated otherwise, all experiments were carried out at 37 °C. Fluorescently labelled WALP23 peptides (WALP23-mCherry or WALP23-msfGFP), msfGFP-MreB as well as GFP-FtsZ were expressed ectopically (*amyE* locus) under control of the *Pxyl* promoter and induced by xylose (1% w/v in case of WALP23; 0.3% w/v in case of GFP-FtsZ). Additional details for strain construction can be found in theAppendix.

### DPH anisotropy measurements

Steady state DPH fluorescent anisotropy measurements were carried out with 1,6-Diphenyl-1,3,5-hexatriene (DPH)-labelled cells using a BMG Clariostar multimode plate reader (BMG Labtec). For *B. subtilis*, cells taken from cultures at time points of interests were diluted to an OD_600_ of 0.25 in a pre-warmed medium, followed by addition of DPH (Sigma Aldrich) dissolved in dimethyl formamide (DMF) to a final concentration of 10 µM DPH and 1% (v/v) DMF. Samples were shaken in dark at 37°C for 5 min, followed by a wash, resuspension in dye-free medium to an OD_600_ of 0.5, and transfer to pre-warmed, black, polystyrene 96-well microtiter plates (Labsystems) for measurement. Following 1 min incubation under shaking in the pre-warmed plate reader to homogenise the sample, the fluorescence anisotropy was measured at 37°C using excitation wavelength of 360-10 nm, emission wavelength of 450-10 nm, and a dichroic mirror set at 410 nm. The fluorescence anisotropy (A) was calculated with MARS Data Analysis software (BMG Labtec) using the equation (I_parallel_-I_perpendicular_)/(I_parallel_+2xI_perpendicular_).

The corresponding measurements for *E. coli* were carried out using the same protocol with following modifications. Staining was carried out with cells grown in the presence of non-growth inhibitory concentrations (30 µg/ml; Sigma Aldrich) of the outer membrane-permeabilising agent Polymyxin B nonapeptide, which is required for good staining of *E. coli* with DPH. The measurements at 30°C and 37°C without temperature shifts were carried out with all media, plastic ware and the plate reader pre-warmed to the corresponding temperatures. The rapid temperature shift from 30°C to 37°C was carried out with cells grown, stained and washed at 30°C, followed by final resuspension in buffer pre-warmed to 37°C and measurement with microtiter plate as well as plate reader pre-warmed to 37°C.

### Determination of fatty acid composition

The fatty acid composition of *E. coli* and *B. subtilis* was determined from 50-100 mg (wet weight) of bacterial cells grown as described in the main text. Fatty acids were extracted as methyl esters after saponification and methylation as described (Sasser, 1990). For saponification, cell pellets were mixed with 15% (w/v) NaOH in 50% (v/v) methanol (1 ml), incubated at 100°C for 5 min, vortexed and further incubated for 25 min. After cooling, acid methylation with 6 N HCl in 50% (v/v) methanol (2 ml) was performed for 10 min at 80°C followed by immediate cooling on ice. Methylated fatty acids were extracted by addition of hexane/methyl tert-butylether in a 1:1 ratio (1.25 ml), followed by end-over-end incubation for 10 min. After phase separation by centrifugation, the lower phase was discarded. The organic phase was washed with 1.2% (w/v) NaOH (3 ml) by 5 min end-over-end incubation and centrifugation. The upper phase of the phase-separated sample was used for further analysis.

The <u>f</u>atty <u>a</u>cid <u>m</u>ethyl <u>e</u>sters (FAME) were separated and identified by gas chromatography-mass spectrometry (GC-MS) with a gas chromatograph (model 7890A; Agilent Technologies) equipped with a 5% phenylmethyl silicone capillary column and a mass spectrometer (model 5975C; Agilent Technologies). Helium was used as carrier gas, injection volume was 1 µl, injector temperature was 250°C, the column temperature was increased from 120 to 240°C at a rate of 5°C/min, and the GC-MS line transfer temperature was 280°C. FAME were separated by their retention times and identified by their equivalent chain lengths and their mass spectra. Equivalent chain length values were calculated from linear interpolation of unknown peaks’ retention time between two saturated straight chain FAME of a standard.

For the analysis of the fatty acid composition of *B. subtilis* wild type cells grown at different temperatures (Figure S9), the cells were grown in LB medium and collected when the cultures reached an OD_600_ of approximately 0.5. The fatty acids were analysed as fatty acid methyl esters with GS-MS as described above. However, these specific analyses were carried out by the Identification Service of the DSMZ, Braunschweig, Germany.

### Glycerophospholipid analysis by MALDI-TOF mass spectrometry

Extraction of lipids from bacterial cells was performed according to Gidden *et al*. (2009) as follows. 10^10^ cells (assuming that 1 ml of cell culture with OD_600_ of 1.0 contains 10^9^ cells (Neidhardt *et al*, 1990) were harvested, washed twice with cooled water and extracted with 450 µl of dichloromethane:ethanol:water 1:1:1 (v:v:v) overnight at 4°C. 1 μl of the lipid-containing lower organic phase was spotted on a MALDI target plate (Prespotted AnchorChip 96 Set for Proteomics II; Bruker Daltonics/Eppendorf; washed peptide-free with 2-propanol) pre-spotted with 1 µl of 9-aminoacridine (10 mg/ml dissolved in acetone:water 9:1 (v/v)) and air-dried. Mass spectra were obtained on an ultrafleXtreme MALDI-TOF/TOF mass spectrometer equipped with a smartbeam™ solid state laser (Bruker Daltonics) operating in the negative ion mode. The laser was fired with a frequency of 500 Hz with 4×250 laser shots per spot. MS/MS spectra were obtained using the ‘LIFT’ technique implemented in the mass spectrometer with an increased laser power. Samples from three independent biological replicates were measured per condition each as technical triplicates, compared with corresponding standard lipids PE(16:0)(18:1) and PG(16:0)(18:1) (Avanti polar lipids) in MS as well as MS/MS spectra, and analysed with FlexAnalysis 3.4 (Bruker Daltonics).

### ONPG membrane permeability assay

For measuring passive ONPG membrane permeability in *E. coli* Δ*lacY* strains carrying plasmid-encoded *lacZ*, overnight cultures were grown at 30°C in LB medium followed by 1:100 dilution in M9-casamino acid medium and growth to an OD_600_ of 0.3. For maintaining the plasmid, media were supplemented with 50 µg/ml ampicillin. Cells at OD_600_ of 0.3 were transferred to clear 96-well microtiter plates followed by addition of 2-nitrophenyl β-D-galactopyranoside (ONPG) dissolved in PBS (Oxoid) to a final concentration of 2 mg/ml. LacZ-driven hydrolysis of ONPG was monitored by measuring absorbance at 420 nm using a BMG Clariostar multimode plate reader, every 2 min for up to 30 min at 37°C with vigorous shaking in between. Meanwhile, the culture was transferred to 37°C for 120 min in a pre-warmed water bath. Cells were then adjusted to OD_600_ 0.3 and ONPG measurements were repeated as above. The relative ONPG permeability was calculated as the rate of ONPG conversion, which is permeability-limited in intact Δ*lacY* cells.

### Fluorescence microscopy

Regular wide field fluorescence microscopy was carried out with cells immobilised on Teflon-coated multi-spot microscope slides (Hendley-Essex) with 1.2% (w/v) agarose/H_2_O (te Winkel *et al*, 2016). In brief, after the agarose solidified within 10 min at room temperature, 0.5 µl of cell culture were applied to the exposed agarose surface, air-dried until the liquid-drop was soaked in, covered with a coverslip and immediately used for microscopy. For staining with various fluorescent dyes, cells were incubated upon shaking at the growth temperature for 5 min with following concentrations: FM 5-95 (2 µg/ml; Thermo Fisher Scientific), DiSC_3_(5) (2 µM; Sigma Aldrich), Sytox Green (50 ng/ml; Thermo Fisher Scientific), DAPI (200 ng/ml; Sigma Aldrich). Membrane depolarisation of *B. subtilis* was achieved by 5 min incubation with small cation specific channel-forming antimicrobial peptide Gramicidin ABC (10 µM; gABC; Sigma Aldrich), and membrane permeabilisation by 5 min incubation with pore-forming lantibiotic Nisin (10 μM; Sigma Aldrich). In the case of *E. coli*, membrane depolarisation and permeabilisation was achieved by 15 min incubation in the presence of pore-forming antibiotic Polymyxin B (10 µg/ml; Sigma Aldrich), and nucleoid staining was achieved by 15 min incubation with 500 ng/ml DAPI (Severn Biotech). Laurdan microscopy was carried out with cells stained with 100 µM Laurdan (Sigma Aldrich; dissolved in 1% (v/v) DMF) as described (Scheinpflug *et al*, 2017a; Wenzel *et al*, 2018). As in case of DPH, staining of *E. coli* was carried out with Polymyxin B nonapeptide outer membrane-permeabilised cells. The time lapse microscopy of *B. subtilis* was carried out with the fortified Spizizen minimal medium (with glucose, tryptophan and casamino acids) diluted to one tenth and supplemented with 1.4% (w/v) low-melting point agarose. The slide preparation was carried out as described (de Jong *et al*, 2011). Fluorescence microscopy of *B. subtilis* and *E. coli* cells stained with FM 5-95, DiSC_3_(5), Sytox Green, Laurdan or expressing fluorescent protein fusions was performed at 37°C with a Nikon Eclipse Ti fluorescence microscope. Structured illumination microscopy (2D-SIM) was carried out with Nikon N-SIM. Additional details on fluorescence microscope equipment can be found in the Appendix. All images were analysed using Fiji. Laurdan GP maps were calculated and generated using the ImageJ-macro described by Wenzel *et al*, 2018). The localisation correlation analysis was carried out with the Fiji plugin Coloc 2, using a 3-pixel wide line following the cell periphery as a region of interest.

For *E. coli fabA*(Ts) time lapse microscopy as well as wide field fluorescence, microscope slides were coated with a thin film of 1% (w/v) agarose dissolved in M9 minimal media supplemented with glucose/casamino acids. Cells (3 µl) were immobilized and imaged with a DeltaVision Elite microscope. Z-Stacks of 300 nm in 5 optical slices were acquired of cells growing on the microscope slide with the microscope tempered to the corresponding temperature and imaged for 2.5 h with 5 min intervals using an exposure time per frame of 50 ms. Image processing, including generation of movies, was performed with Fiji. Additional details can be found in the Appendix.

### Single molecule tracking

Single molecule imaging of *E. coli* cells expressing F_O_F_1_ *a*-mNG was performed using a total internal reflection fluorescence (TIRF) microscope. Cells were pre-bleached for 2.5 s at 10 % laser intensity (approx. 1.5 µW/µm^2^) to obtain single molecule fluorescence level. Subsequently, single emitter signals were imaged at 30 frames per second for 1200 frames (40 s) with 5 % of laser intensity. All experiments were carried out at room temperature for comparability. Tracking of single molecules and data analysis were carried out with well-established localisation and tracking algorithms, implemented in a software package called ‘SLIMfast’ (kindly provided by C.P. Richter (Osnabrück)) written in Matlab (Appelhans *et al*, 2018; Richter *et al*, 2017). Localisation precision was typically about 20-25 nm. Between 900 and 1000 frames per image series were used for further step length and diffusion constant analysis. Step length analysis is based on trajectories exhibiting at least five sequential frames (excluding deflation loops and frame gaps). Typically, the population of all trajectories (step size of ≥1) is approximately twice as high as those taken into account (step size of ≥5).

Analysis of lateral mobility was performed via cumulative probability plots with jump sizes of pooled trajectories from 3-5 separately grown cell batches and via boxplots to determine the median of all trajectories present within individual cells. Apparent two-dimensional diffusion coefficients *D*_*app*_ were estimated by the <u>m</u>ean-<u>s</u>quared <u>d</u>isplacement (MSD) ⟨(Δ*r*(*τ*))^2^⟩ =4*D*_*app*_*τ* considering a linear free diffusion model. Here, *τ* = Δ*t*, 2Δ*t*, …, *n*Δ*t* is the lag time defined by multiples of time interval Δ*t* of the image series. For all trajectories with at least ≥5 five sequential frames, MSD was averaged and the diffusion coefficient calculated by the slope of a linear fit based on the first four data points of the MSD. For determination of standard error, the statistical resampling method of bootstrapping was used, evaluating data sets with N data points 1000-times (Bradley, 1981).

### Quantification and statistical analysis

Statistical analysis was performed using a two-sided Wilcoxon rank sum test (MATLAB) or an unpaired two-sided t-test (GraphPad Prism or Microsoft Excel). Error bars represent SD from three independent biological replicates, unless stated otherwise. Significance was assumed with **** p < 0.0001, *** p < 0.001, ** p < 0.01, * p < 0.05, n.s., not significant.

## Data Availability

Additional methods, figures, tables, and movies can be found in the Supplemental information. Correspondence and requests for material should be addressed to H.S. or G.D.-H. This study includes no data deposited in external repositories.

## Acknowledgements

We thank E. Garner (Harvard), Tanneke den Blaauwen (Amsterdam), S. Lee (Newcastle), and K. Jahreis (Osnabrück) for providing strains/plasmids; E. Becker, L. Fellner, A. Bogdanowski (Osnabrück), and Kenneth H. Seistrup (Newcastle) for aid in generating plasmids/strains and for generating wild type fatty acid profiles used in comparisons; H. Winkelmann, H. Arlt for introducing MG into fluorescence microscopy; C.P. Richter, T. Appelhans for technical discussions (Osnabrück). This work was supported by CRC944 (to G.D.-H.), ‘Incentive Award’ of the faculty of Biology/Chemistry of Osnabrück University (to G.D.-H.) and BBSRC New Investigator Award BB/S00257X/1 (to H.S.). MG is supported by a stipend of the Hans Mühlenhoff-Stiftung (Osnabrück).

## Author contributions

G.D.-H. and H.S. designed research; G.D.-H. and H.S. coordinated the collaborative research; M.G., A.L., J.W.G., J.A.B., B.H., Z.B. and H.S. performed the experiments; M.G., A.L., J.W.G., J.A.B., R.K., S.W., G.D.-H. and H.S. analysed data; H.S., G.D.-H., and M.G. wrote the paper.

