## Supplemental information for "Low membrane fluidity triggers lipid phase separation and protein segregation *in vivo*"

**Supplemental Methods, Tables, Figures and Movies**

### Construction of *E. coli* strains

All *E. coli* strains used are listed in Table S1. *E. coli* strains MG1 or EB8.1 carrying a C-terminal mNeonGreen (mNG) or mCherry fusion via a Gly-Ser linker to the membrane-integral $F_O$-*a* subunit, respectively, were generated using the phage λ Red recombinase to replace a chromosomal sequence (Datsenko & Wanner, 2000). Briefly, the kanamycin resistance cassette of strain EB4 (Δ*atpBE::*FRT-*kan*-FRT) was exchanged by the HindIII/AseI fragment of plasmids pBH189 (MG1) or pEB21.2 (EB8.1), followed by growth on M9 minimal medium with succinate (0.4% (w/v)) as sole carbon source for selection. As expected, both strains show a $Succ^+$ $Kan^S$ phenotype. In detail, EB4 cells transformed with temperature-sensitive plasmid pKD46 encoding λ Red recombinase genes under control of the *ParaBAD* promoter were grown at 30°C to mid-logarithmic phase in lysogenic broth (LB) composed of yeast extract (0.5% w/v), tryptone (1% w/v), NaCl (1% w/v) and supplemented with arabinose (0.2% w/v) for induction. Competent cells were prepared by wash with ice-cold water for removal of salts and medium components, electroporated in the presence of corresponding DNA fragments, grown for 1 h in LB at 37°C for phenotypic expression and plated on selective solid medium.

For generation of *E. coli* strain UC1098.PtsG-mNG, a 75 bp linker from plasmid pBLP2 (encoding the amino acids EFTMVPAAPAPAAAAPAAAPTPASR) and the open reading frame (ORF) encoding mNG (pNCS-mNeonGreen) were inserted into the chromosomally encoded *ptsG* gene prior to its stop codon using λ Red mutagenesis (Datsenko & Wanner, 2000). Briefly, the kanamycin resistance cassette of strain UC1098.Δ*ptsG* (Δ*ptsG::*FRT-*kan*-FRT) (see below) was exchanged by a PCR product harbouring the *ptsG-linker-mNG* fusion gene flanked by upstream/downstream chromosomal regions of *ptsG* using growth on M9 minimal medium with glucose (0.4% (w/v)) for selection. The PCR product was generated by a two-step PCR overlap extension method. Firstly, four individual PCR products were generated using (*i*) oligonucleotides 1/2 with lysed cells of *E. coli* strain Y-Mel as template, (*ii*) oligonucleotides 3/4 with pBLP2 as template, (*iii*) oligonucleotides 5/6 with pNCS-mNeonGreen as template and (*iv*) oligonucleotides 7/8 again with lysed cells of Y-Mel as template. Secondly, the four PCR products and oligonucleotides 1/8 were used for the second amplification step. Oligonucleotides used are listed in Table S2.

Several *E. coli* strains were obtained by P1 transduction (Thomason *et al*, 2007). In detail, P1 liquid lysate was generated by growing the donor strain to optical density at 600 nm ($OD_{600}$) of 0.1 in LB (3 ml), adding $CaCl_2$ (330 µl 50 mM) and P1 lysate (20 µl of ~$10^{-9}$ phages/ml) and further growth with good aeration until lysis occurred. 5 drops of chloroform were added to lyse remaining cells, centrifuged twice to pellet debris, and the supernatant was stored with 50 µl of chloroform in the dark at 4°C. For transduction, an overnight culture of the recipient strain (200 µl) was mixed with $CaCl_2$ (28 µl 50 mM) and P1 lysate (50 µl) of the donor strain and incubated for 20 min at 37°C. After addition of LB (0.7 ml) and $Na_3$-citrate (100 µl 1 M) and further incubation for 40 min, cells were plated on selective solid medium containing $Na_3$-citrate (20 mM).

Strains LF4 and AB1 were obtained using EB4 as donor, UC1098 (kindly provided by J.E. Cronan, Jr. (Illinois)) and Cy288 as recipients, respectively, using LB with kanamycin (50 µg/ml) for selection leading to a $Succ^+$ $Kan^R$ phenotype. Subsequently, MG4 and LF6.red were generated by P1 transduction using MG1 and EB8.1 as respective donor strains, LF4 as recipient and M9 minimal medium with succinate (0.4% (w/v)) for selection. Strain BHH87 was generated in the same way as described for strain MG4. For generation of strains BHH100 and BHH101 by P1 transduction, MG1655.mreB-msfGFP and KC555, respectively, were used as respective donor strains, UC1098 as recipient and LB with kanamycin (50 µg/ml) or chloramphenicol (12.5 µg/ml) for selection. For generation of strain UC1098.Δ*ptsG*, strain JW1087-2 was used as donor, UC1098 as recipient and LB with kanamycin for selection. For generation of strains Y-Mel.Δ*lacY* and UC1098.Δ*lacY*, strain JW0334-1 was used as donor, Y-Mel or UC1098 as recipients and LB medium with kanamycin for selection. For generation of UC1098.Δ*minC*, strain JW1165-1 was used as donor, UC1098 as recipient and LB medium with kanamycin for selection. In all strains, the genes of interest were verified by colony PCR and DNA sequencing.

**Construction of *E. coli* plasmids**

For construction of plasmids pBH189 and pEB21.2, a BamHI site (encoding a Gly-Ser linker) and genes encoding mNG (pBH189) and mCherry (pEB21.2), respectively, were inserted into the *atp* operon of plasmid pBWU13 prior to the stop codon of *atpB* using a two-step PCR overlap extension method. Firstly, three individual PCR products were generated using (*i*) oligonucleotides 9/10 with pSD166 as template, (*ii*) oligonucleotides 11/12 with pNCS-mNeonGreen as template for pBH189 or with pQW58 as template for pEB21.2 and (*iii*) oligonucleotides 13/14 with pSTK3 as template. Secondly, the three PCR products and oligonucleotides 9/14 were used for the second amplification step. HindIII/AseI-digested PCR products were cloned into correspondingly digested pBH4.

For construction of plasmids pBH500 and pBH501, a linker encoding SGSGSG, and the ORFs encoding mNeonGreen and mScarlet-I, respectively, were fused with WALP23-ORF by two-step PCR. Briefly, for plasmid pBH500, two different PCR products were obtained using oligonucleotides 15/16 with pL030 as template and oligonucleotides 18/19 with pNCS-mNeonGreen as template. For the second PCR step, oligonucleotides 15/19 were used. For plasmid pBH501, two PCR products were obtained using oligonucleotides 15/17 with pL030 as template and oligonucleotides 20/21 with synthesized mScarlet-I-encoding DNA as template. For the second PCR step, oligonucleotides 15/21 were used. In both cases, AvrII/SpeI-digested PCR products were cloned into correspondingly digested pL030.

For construction of plasmids pJG130 and pJG131, plasmid pBAD322 was linearised with oligonucleotides 26 and 27. Full length *rne* was amplified using oligonucleotides 28/29, while inserts for construction of *rne*-ΔAH were amplified using oligonucleotides 28/30 and 29/31. These fragments were fused using NEBuilder® HiFi DNA Assembly Cloning Kit (New England Biolabs).

All constructs were verified by DNA sequencing and are listed in Table S3. Oligonucleotides used are listed in Table S2.

**Construction of *B. subtilis* strains**

All *B. subtilis* strains used are listed in Table S1. For construction of a *B. subtilis* strain expressing

WALP23 fused to monomeric superfolder GFP (msfGFP), the plasmid pBH500 was linearised with oligonucleotides 22 and 23, msfGFP amplified using oligonucleotides 24 and 25 (Table S2) and the fragments fused using NEBuilder® HiFi DNA Assembly Cloning Kit (New England Biolabs). The resulting plasmid was transformed into *B. subtilis* 168, thus resulting in strain JG054. All other *B. subtilis* strains were constructed by transforming the respective recipient strains with chromosomal DNA from the donor strains or corresponding plasmid DNA. Transformations were carried out as described (Hamoen *et al*, 2002).

**Fluorescence microscopy**

The fluorescence microscopy of *B. subtilis* and *E. coli* cells stained with FM 5-95, DiSC$_3$(5), Sytox Green, Laurdan or expressing fluorescent protein fusions was performed at 37°C with Nikon Eclipse Ti equipped with either Sutter Instruments Lambda LS Xenon-arc light source or CoolLed pE-300white LED light source, CoolLed pE-4000 LED light source, Photometrics Prime sCMOS camera, Photometrics BSI sCMOS camera and either Nikon Plan Fluor 100x/1.30 NA Oil Ph3, Nikon CFI Plan Apo VC 100x/1.40 NA or Nikon Plan Apo 100×/1.40 NA Oil Ph3 objectives. The used filter sets were Chroma 49000 (for DAPI and Laurdan 460 nm), Chroma 49002 (for GFP and Sytox Green), Chroma 49003 (for YFP), Chroma 49008 (for mCherry, mScarlet-I and FM5-95), Semrock Cy5-4040C (for DiSC$_3$(5)), a custom filter set consisting of Chroma AT350/50x excitation filter, Chroma T400lp beam splitter, or Chroma ET525/50m (for Laurdan 520 nm). The microscopy images shown in Figure S10 were carried out with Applied Precision DeltaVision RT equipped with Photometrics Coolsnap HQ2 camera, Zeiss Plan-APOCHROMAT 100x objective and the standard DeltaVision filters sets. Dual-color 2D-SIM was carried out with Nikon N-SIM equipped with Nikon CFI APO TIRF 100x/1.49 oil objective, 488 nm (Coherent Sapphire) and 561 nm (Cobolt Jive 100) solid-state lasers and Andor Xion X3 EMCCD camera. The used filter sets were Chroma 49904 (for mNG) and 49909 (for mCherry). Image capture and reconstruction of high-resolution 2D-SIM images were performed with NIS elements 5.11 (Nikon). All images were analysed using Fiji (Schindelin *et al*, 2012). Laurdan GP maps were calculated and generated using the ImageJ-macro as described (Wenzel *et al*, 2018). The localisation correlation analysis was carried out with the Fiji plugin Coloc 2, using a 3-pixel wide line following the cell periphery as a region of interest.

For *E. coli fabA*(Ts) time lapse microscopy as well as widefield fluorescence microscopy microscope slides were coated with a thin film of 1% (w/v) agarose dissolved in M9 minimal media supplemented with glucose/casamino acids as described in the main text. Cells (1 or 3 µl) were immobilized and imaged with a DeltaVision Elite microscopy system (Applied Precision, GE Healthcare) equipped with an inverted microscope (IX-71, Olympus), a 100x oil immersion objective (UAPON 100x TIRF, Olympus) or an extended apochromat phase-contrast objective (UPLXAPO100XOPH, Olympus), solid state illumination system (Insight SSI, Applied Precision), a sCMOS camera (pco.edge 4.2, PCO) and acquisition software (softWoRx 5.5, Applied Precision). Fluorescence of mNeonGreen and mScarlet-I was excited using a polychromic beamsplitter (405 nm/488 nm/590 nm/650 nm) as well as either a GFP/FITC bandpass filter (461-489 nm) or a mCherry/Alexa594 bandpass filter (562-588 nm).

Fluorescence detection was achieved using a GFP/FITC bandpass emission filter (501-559 nm) for mNG and a mCherry/Alexa594 bandpass emission filter (602-648 nm) for mScarlet-I.

Single molecule imaging of *E. coli* cells expressing $F_OF_1$ *a*-mNG or WALP23-mNG was performed using a total internal reflection fluorescence (TIRF) microscopy system equipped with an inverted microscope (IX-83, Olympus), a motorized four-line TIRF condenser (cellTIRF, Olympus), an 150x oil immersion objective (UAPON 150x/1.45 NA TIRF, Olympus), an EMCCD camera (iXON Ultra 897, Andor) and the acquisition software CellSens 2.3 (Olympus). Fluorescence of mNG was excited by a 488 nm laser diode (LuxX 488-200, Omicron) using a TIRF pentaband polychroic mirror (zt405/488/561/640/730rpc, Chroma). Fluorescence detection and efficient TIRF laser blocking was achieved by a pentabandpass emission filter (BrightLine HC 440/521/607/694/809, Semrock) and an additional single bandpass emission filter (BrightLine HC 525/35, Semrock).

#### Membrane vesicles and DCCD-sensitive ATPase activity

Inverted membrane vesicles were prepared as previously described (Brandt *et al*, 2013) using 850 ml of cell culture grown in M9 minimal medium with 0.4% (w/v) glucose and 0.1% (w/v) casamino acids. After harvest, cell pellets were resuspended in 25 ml 50 mM Tris-HCl, pH 7.5, 10 mM $MgCl_2$, 10% (v/v) glycerol and disrupted in the presence of 10 µg/ml DNaseI (Sigma) with a constant cell disruptor system (Daventry) at 4°C and 1.35 kbar. For removal of cell debris, lysates were centrifuged at 35 000x*g* for 30 min at 4°C. After ultracentrifugation of the supernatant at 250 000x*g* at 4°C for 60 min, membranes were resuspended with a marten paint brush in a small aliquot (0.3-0.5 ml) of the same buffer and stored in liquid nitrogen.

ATPase activities of inverted membrane vesicles were determined using an automated continuous assay enabling a direct recording of the substrate turnover (Arnold *et al*, 1976). For inhibition of ATPase activities with 80 µM *N*,*N'*-dicyclohexylcarbodiimide (DCCD; Sigma Aldrich; stock solution 40 mM in ethanol), membranes were incubated in 1 ml of 50 mM Tris-HCl, pH 8.0 for 20 min at 37°C prior to measurement (Deckers-Hebestreit & Altendorf, 1992).

#### SDS-PAGE and Western blot

Protein concentrations were determined with the BCA assay as recommended by the supplier (Pierce). Proteins (20 µg/lane) were separated by SDS-PAGE using 10% tris-tricine gels (10% T, 3% C) (Schägger & von Jagow, 1987) with PageRuler™ prestained protein ladder (Fermentas) as standard. For immunoblotting, separated proteins were transferred to nitrocellulose membranes (0.45 µm) via wet blotting in carbonate buffer (10 mM $NaHCO_3$, 3 mM $Na_2CO_3$, pH 8.9 with NaOH, 20% (v/v) methanol) for 45 min at 1.8 A with cooling (Hilbers *et al*, 2013). Membranes were blocked with 5% (w/v) skimmed milk powder in TBS buffer (50 mM Tris/HCl pH 7.4, 0.9% (w/v) NaCl), decorated with monoclonal mouse antibodies specific for $F_O$-*a* (GDH 14-5C6 (Jäger *et al*, 1998)) or mNeonGreen (32F6, ChromoTek) and secondary IRDye™800DX-labelled goat-anti-mouse IgG (H+L) (LI-COR Biosciences), and detected using a two-channel Odyssey infrared imaging System (LI-COR Biosciences).

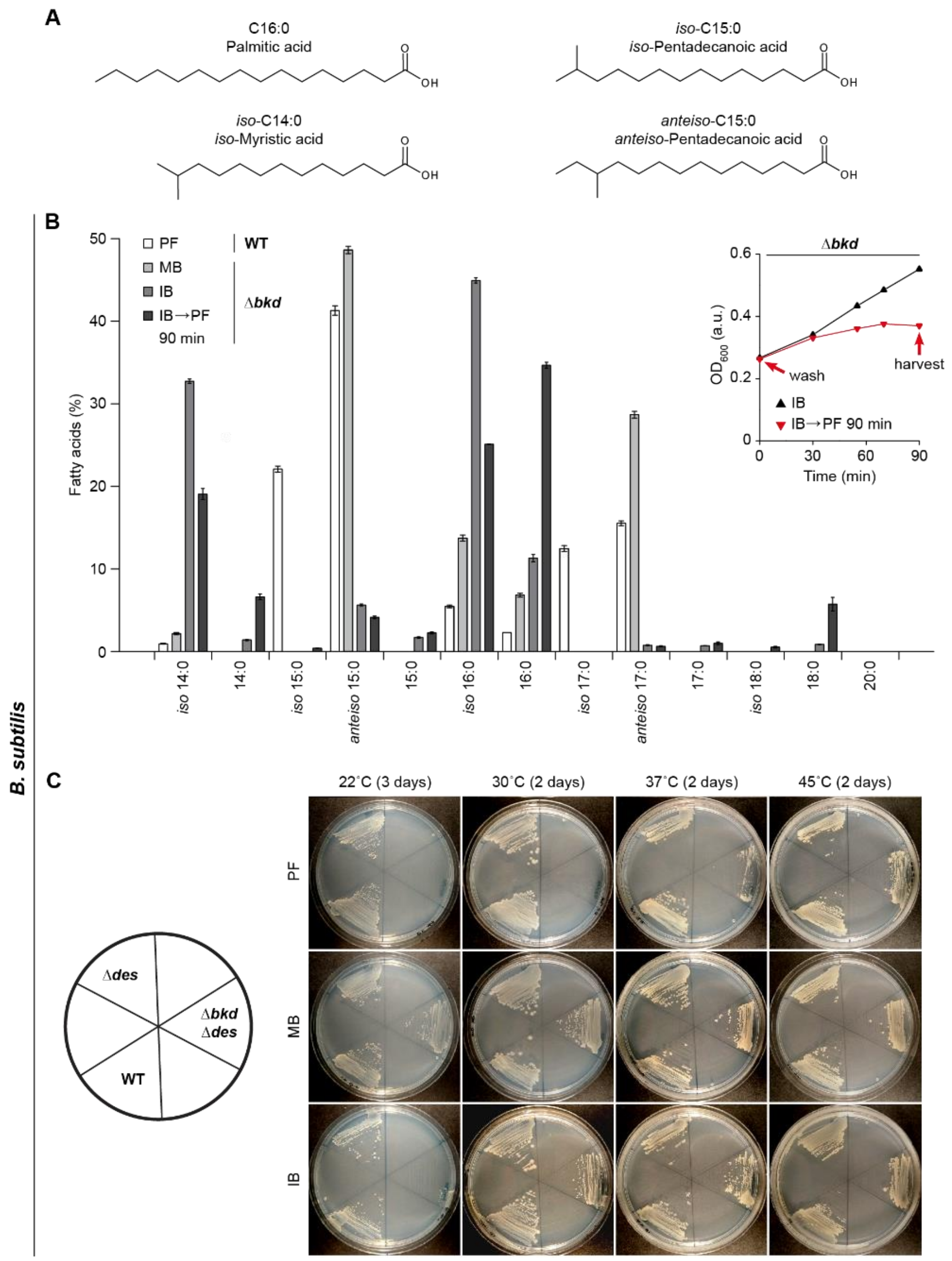

**Figure S1. Depletion of branched chain fatty acids in *B. subtilis*.** (**A**) Structures of fatty acids typical for lipids in the plasma membrane of *B. subtilis* comprising BCFAs as fluidity-promoting and SFAs as fluidity-reducing fatty acids. Whereas IB is the precursor for *iso*-fatty acids with even numbers of C atoms such as *iso*-C14:0 or *iso*-C16:0, MB is used for the synthesis of *anteiso*-C15:0 and *anteiso*-C17:0. (**B**) Detailed fatty acid composition of *B. subtilis* wild type cells grown in the absence of fatty acid precursors, and cells of the fatty acid precursor auxotroph strain Δ*bkd* grown in the presence of MB, IB or grown precursor-free (PF) for 90 min (IB→PF). The determination was carried out through GC-MS of fatty acid methyl esters. The insert depicts the corresponding growth behaviour of the Δ*bkd* strain upon pre-culturing in the presence of IB, followed by wash and resuspension in either IB-supplemented or precursor-free (IB→PF 90 min) medium. The time point of cell harvest for the lipid analyses is indicated. (**C**) Temperature-dependent growth behaviour of *B. subtilis* wild type cells, a strain deficient for the lipid desaturase Des (Δ*des*) and the fatty acid precursor auxotroph *B. subtilis* strain carrying deletions of both *des* and the *bkd* operon (Δ*bkd* Δ*des;* named "Δ*bkd*" for simplicity throughout the text) on solid growth medium supplemented either with MB, IB or grown precursor-free. Plates were incubated at temperatures of 45°C, 37°C and 30°C for 2 days, and 22°C for 3 days. **Data information**: (**B**) The histogram depicts means and SD of biological triplicates for each strain and temperature condition. (**C**) Experiments are representative of three independent repeats. (**B**) *B. subtilis* 168, HS527; (**C**) *B. subtilis* 168, KS20, HS527.

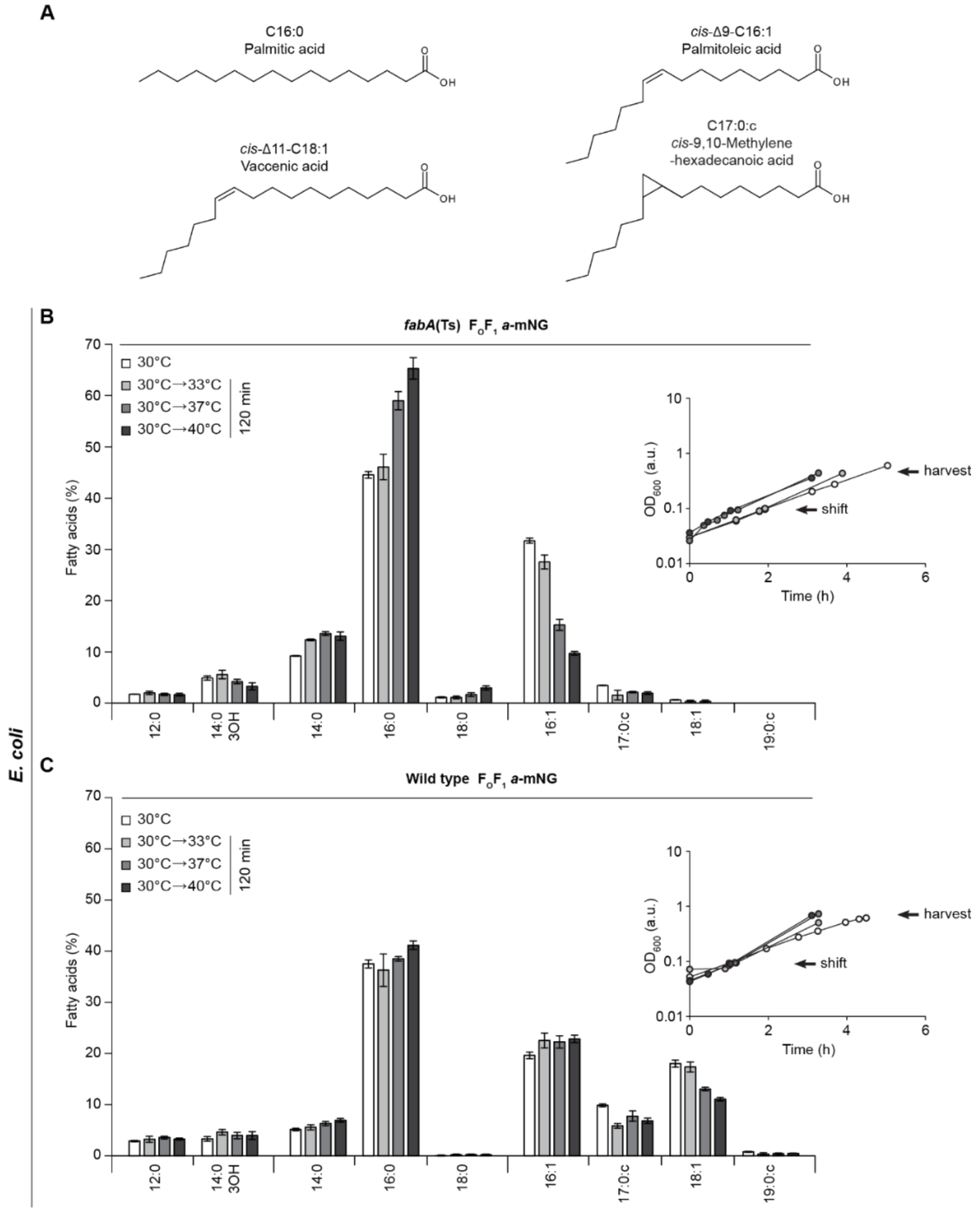

**Figure S2. Depletion of unsaturated fatty acids in *E. coli.*** (**A**) Structures of fatty acids typical for lipids in the plasma membrane of *E. coli* comprising UFAs as fluidity-promoting and SFAs as fluidity-reducing fatty acids. Phospholipids contain as main fatty acids C16:0 and C16:1, while the amount of C18:1 is decreased at higher temperatures. During oxidative or acid stress as well as in the stationary growth phase, UFAs are converted into cyclopropane fatty acids (CFA) such as C17:0:c or C19:0:c to retain the *cis* double-bond configuration. (**B**) Detailed fatty acid composition of the temperature-sensitive *E. coli* strain *fabA*(Ts) (*fabF fabA*(Ts); named "*fabA*(Ts)" for simplicity throughout the text) grown at 30°C, or grown at 30°C followed by a shift to 33°C, 37°C or 40°C for 120 min. (**C**) Detailed fatty acid composition of *E. coli* wild type cells grown as described in panel B for *fabA*(Ts). (**B**, **C**) Inserts show the corresponding growth curves. The time point of the temperature shift as well as the cell harvest for the lipid analyses is indicated. Fatty acids C12:0 and C14:0-3OH were not included into the summarised fatty acid profiles shown in Figure 1E due to their sole presence in lipopolysaccharide molecules. The determination was carried out through GC-MS of fatty acid methyl esters. **Data information**: (**B**, **C**) The histograms depict means and SD of biological triplicates for each strain and temperature condition. Strains used: (**B**) *E. coli* MG4; (**C**) *E. coli* MG1, (strains Y-Mel and UC1098, respectively, additionally encoding fluorescent ATP synthase ($F_OF_1$ *a*-mNG)).

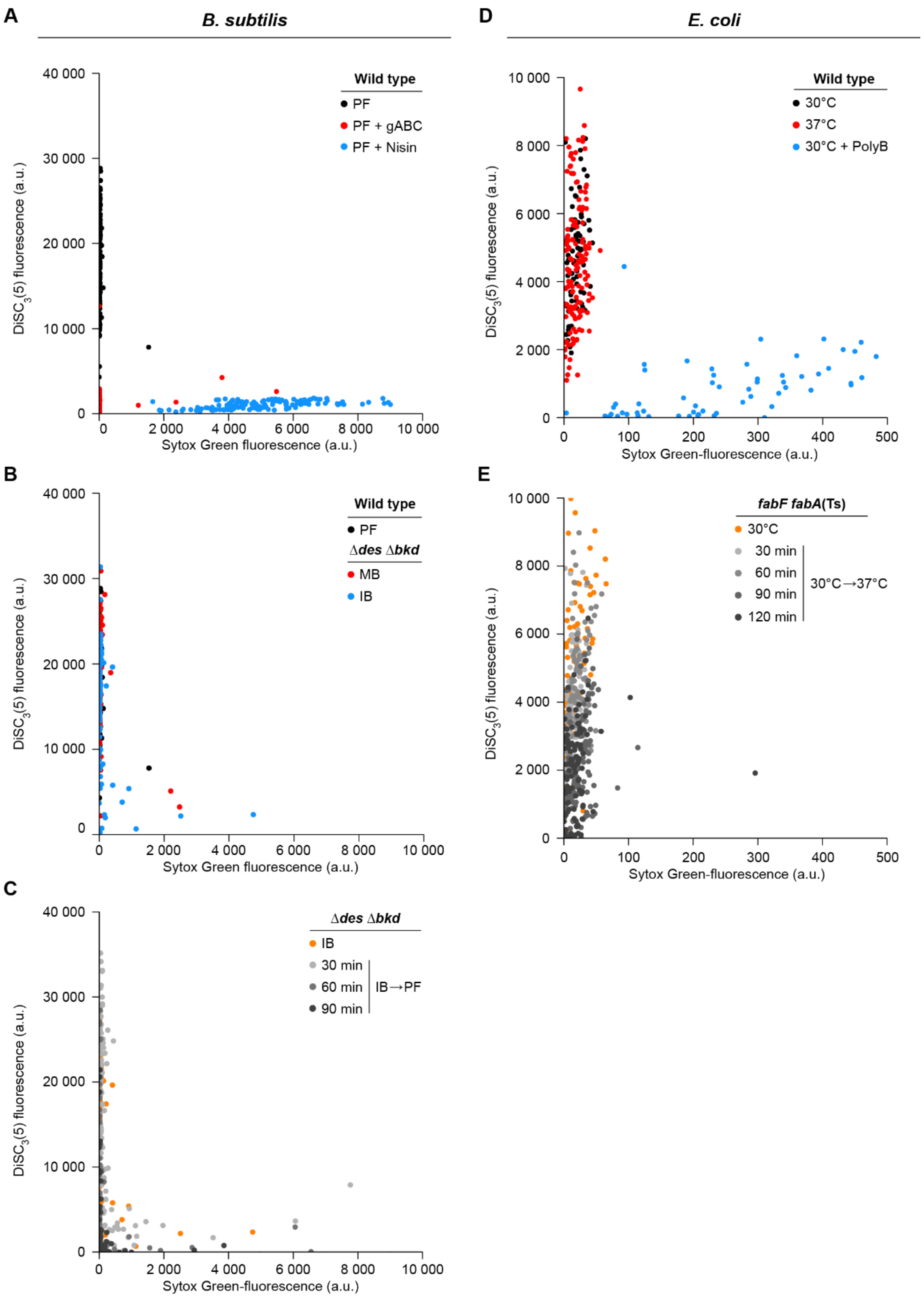

**Figure S3. Cross-correlation between membrane depolarisation and membrane permeabilisation.** (**A**) Correlation between membrane depolarisation and permeabilisation in *B. subtilis* wild type cells co-stained with the membrane potential-sensitive dye DISC3(5) and the membrane permeability indicator Sytox Green, respectively. Depicted are the quantified cellular fluorescence signals for untreated wild type cells grown in the absence of fatty acid precursors, and the same cells incubated with gramicidin ABC (gABC), a membrane-depolarising but not permeabilising antimicrobial peptide, or with permeabilising, pore-forming lantibiotic nisin. (**B**) Correlation between membrane depolarisation and permeabilisation in *B. subtilis* fatty acid precursor auxotroph strain Δ*bkd* grown in the presence of fatty acid precursor MB or IB and co-labelled with $DISC_3(5)$ and Sytox Green. (**C**) Correlation between membrane depolarisation and permeabilisation in *B. subtilis* Δ*bkd* grown in IB, washed by centrifugation, further incubated in the absence of fatty acid precursors for the time points indicated and co-labelled with $DISC_3(5)$ and Sytox Green. (**D**) Correlation between membrane depolarisation and permeabilisation in *E. coli* wild type cells grown at 30°C or 37°C and co-labelled with $DISC_3(5)$ and Sytox Green. As a control, wild type cells grown at 30°C were incubated in the presence of the pore-forming antibiotic Polymyxin B. (**E**) Correlation between membrane depolarisation and permeabilisation in *E. coli fabA*(Ts) cells grown at 30°C and transferred to the non-permissive temperature of 37°C for the time points indicated, and co-stained with $DISC_3(5)$ and Sytox Green. **Data information**: (**A**-**E**) Data shown here represent the same dataset (n= 100-142) used for generating the graphs shown in Figures 3B and 3D, but were additionally analysed for the Sytox Green fluorescent signal and its correlation with corresponding $DISC_3(5)$ fluorescence signals. Strains used: (**A**-**C**) *B. subtilis* 168, HS527; (**D**, **E**) *E. coli* Y-Mel, UC1098.

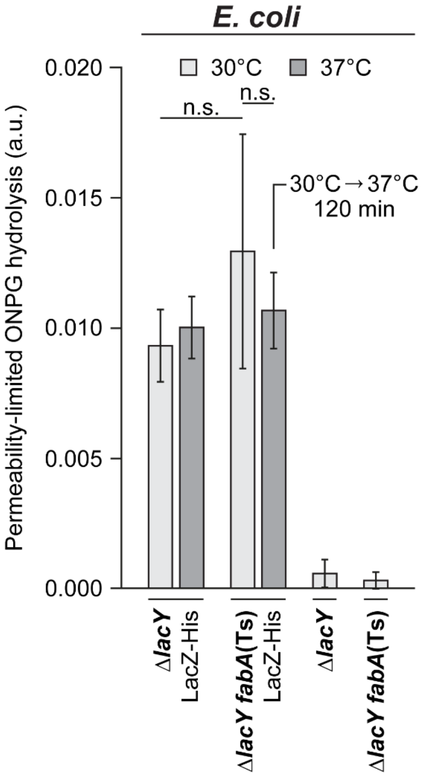

**Figure S4. Very low membrane fluidity in *E. coli* does not trigger membrane permeabilisation for *ortho*-nitrophenyl β-D-galactopyranoside (ONPG).** The membrane permeability of ONPG was assessed in strains deficient for the active uptake system LacY simultaneously expressing *lacZ* from a plasmid-encoded leaky *Ptac* promoter (without addition of the inducer IPTG). The graph depicts ONPG hydrolysis rates measured for intact cells upon incubation at 30°C and 37°C. In case of the temperature-sensitive FabA(Ts) strain, growth at the non-permissive temperature of 37°C was limited to 120 min. As controls, the ONPG conversion rates were measured in strains lacking the *lacZ*-expressing plasmids. Note the lack of significant difference in ONPG conversion rates upon strong depletion of unsaturated fatty acids (FabA(Ts) strain at 37°C for 120 min), implying the lack of detectable membrane permeabilisation due to phase separation. **Data information**: The graph depicts mean and SD of biological triplicates. The P values represent the results of unpaired, two-sided t-tests. Insignificant changes ($p>0.1$) are indicated with n.s. Strains used: *E. coli* Y-Mel.Δ*lacY*, UC1098.Δ*lacY*, Y-Mel.Δ*lacY*/pTM30.*lacZ*-His2, UC1098.Δ*lacY*/pTM30.*lacZ*-His2.

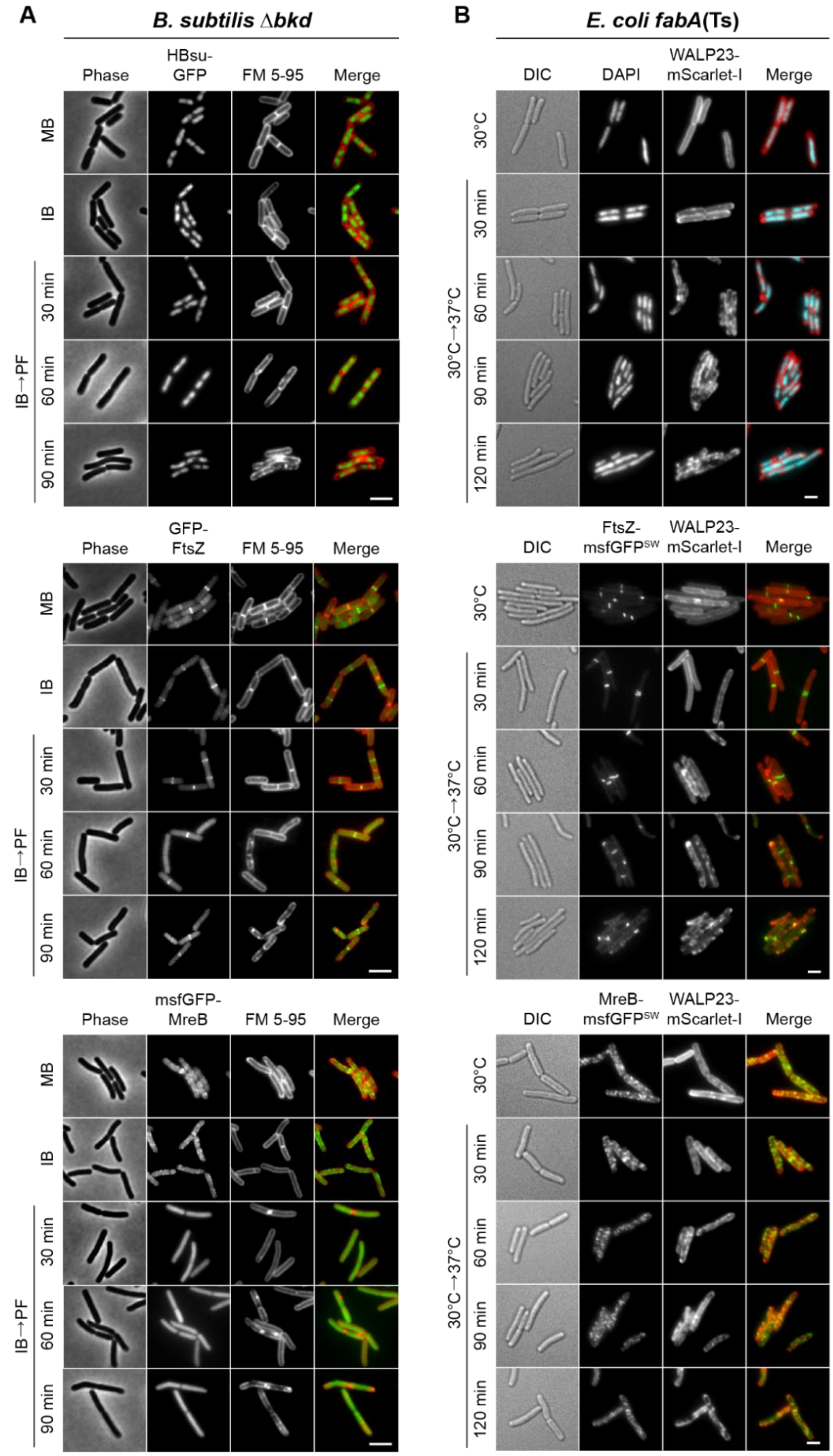
A
B. subtilis Δbkd
Phase
HBsu-GFP
FM 5-95
Merge
MB
IB
IB→PF
30 min
60 min
90 min
GFP-FtsZ
msfGFP-MreB
B
E. coli fabA(Ts)
DIC
DAPI
WALP23-mScarlet-I
30°C
30°C→37°C
120 min
FtsZ-msfGFPSW
MreB-msfGFPSW

**Figure S5. Consequences of low membrane fluidity on cell morphogenesis.** (**A**) Detailed phase contrast and fluorescence images of *B. subtilis* fatty acid precursor auxotroph strain Δ*bkd* stained with the membrane dye FM 5-95 and expressing GFP fusions to the sequence-unspecific DNA-binding protein HBsu (top), cell division protein FtsZ (middle) or cell elongation protein MreB (bottom). Depicted are cells grown in the presence of the precursor IB or MB, and cells grown with IB, washed followed by precursor-free incubation (IB→PF) for the time points indicated. (**B**) Detailed differential interference contrast (DIC) and fluorescence images of *E. coli fabA*(Ts) cells, incubated with dye FM 5-95 (staining the outer membrane), were labelled in addition with DAPI for DNA (top), or expressed GFP fusions to FtsZ (middle) and MreB (bottom), respectively. Depicted are cells grown at the permissive growth temperature (30°C) and upon growth shift to the non-permissive temperature (37°C) for the time points indicated. (**A**, **B**) The panels provide additional examples and time points for the images shown in Figure 4. **Data information**: Experiments are representative of biological triplicates. (**A**) Scale bar 3 µm. (**B**) Scale bar 2 µm. Strains used: (**A**) *B. subtilis* HS541, HS548, HS549; (**B**) *E. coli* UC1098, BHH100, BHH101.

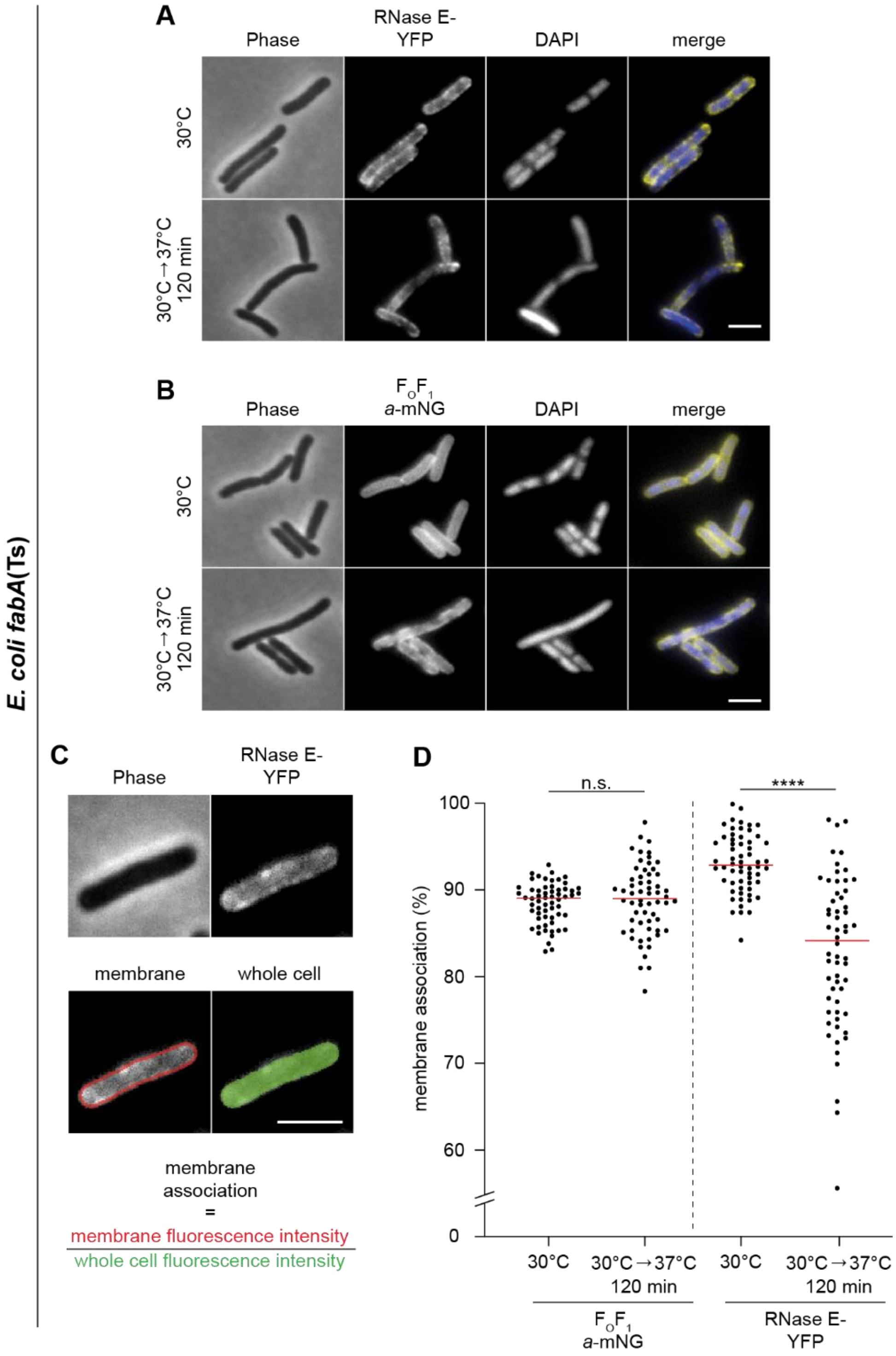

**Figure S6. Very low membrane fluidity triggers partial dissociation of RNase E from the membrane in *E. coli fabA*(Ts).** (**A**, **B**) Phase contrast and fluorescence images of *E. coli fabA*(Ts) strain expressing (A) RNase E-YFP or (B) $F_OF_1$ *a*-mNG. Cells were grown in LB at 30°C to an $OD_{600}$ of 0.3, transferred to the non-permissive temperature of 37°C for 120 min followed by labelling with DAPI and fluorescence microscopy. Note the increasing cytoplasmic localisation of RNase E-YFP upon depletion of the membrane for UFA, which coincides with decondensation of the nucleoid (compare Figure 4B). (**C**) Quantification of membrane association of RNase E. The degree of membrane association was quantified by automated detection of cells using phase contrast images, defining a 3-pixel wide band around the periphery of the cell, and measuring the relative membrane association as a ratio between the mean peripheral fluorescence signal and the mean fluorescence of the whole cell. (**D**) Relative membrane association of $F_OF_1$ *a*-mNG and RNase E-YFP at 30°C and 37°C for 120 min in individual cells (n=60). Red lines indicate the median. **Data information**: (**A**-**D**) Experiments are representative of independent biological duplicates. (**D**) Red lines indicate the median. P values represent the results of unpaired, two-sided t-tests. Significance was assumed with **** $p < 0.0001$, *** $p < 0.001$, ** $p < 0.01$, * $p < 0.05$, n.s., not significant. (**A**-**C**) Scale bar: 3 µm. Strains used: (**A**, **C**, **D**) *E. coli* UC1098/pVK207; (B, **D**) *E. coli* MG4.

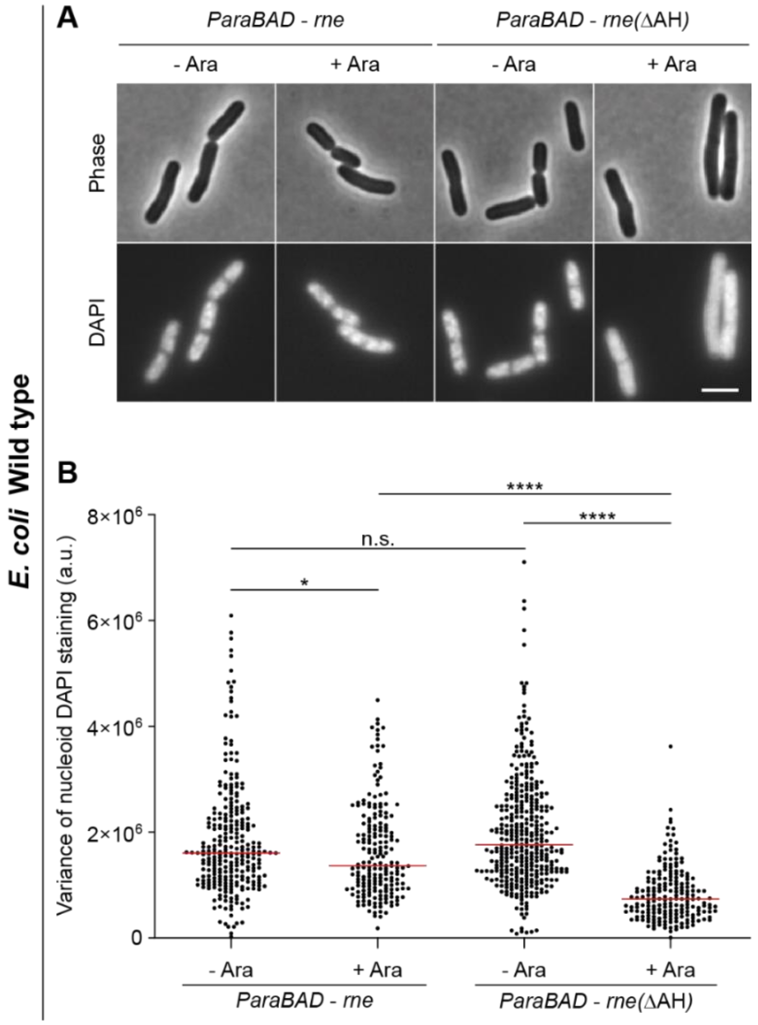

**Figure S7. Expression of cytoplasmic RNase E is sufficient to trigger decondensation of the nucleoid.** (**A**) Phase contrast and fluorescence images of *E. coli* WT cells expressing plasmid-encoded full-length membrane-associated RNase E (Rne) and a corresponding construct encoding RNase E that lacks the membrane-binding amphipathic helix (RneΔAH), respectively. The cells depicted were grown in LB medium at 37°C to an $OD_{600}$ of 0.3 followed by induction of *rne* with 0.2% (w/v) arabinose (Ara) for 60 min, labelling with DAPI for 15 min and fluorescence microscopy. Note the decondensation of the nucleoid observed upon expression of cytoplasmically located RneΔAH, but not in the presence of the native membrane-associated RNase E. (**B**) Variance of nucleoid staining. The degree of nucleoid condensation was assessed by analysing the variance of DAPI fluorescence within the cell (n=199-372). In this type of analysis, a more homogeneous fluorescence signal such as that caused by nucleoid decondensation results in a lower variance of the per pixel fluorescence intensity. **Data information**: (**A**, **B**) The experiments are representative of biological duplicates. (**B**) Red lines indicate the median. P values represent results of unpaired, two-sided t-tests. Significance was assumed with **** $p < 0.0001$, *** $p < 0.001$, ** $p < 0.01$, * $p < 0.05$, n.s., not significant. (**A**) Scale bar: 3 µm. Strains used: (**A**, **B**) *E. coli* MG1655/pJG130, MG1655/pJG131.

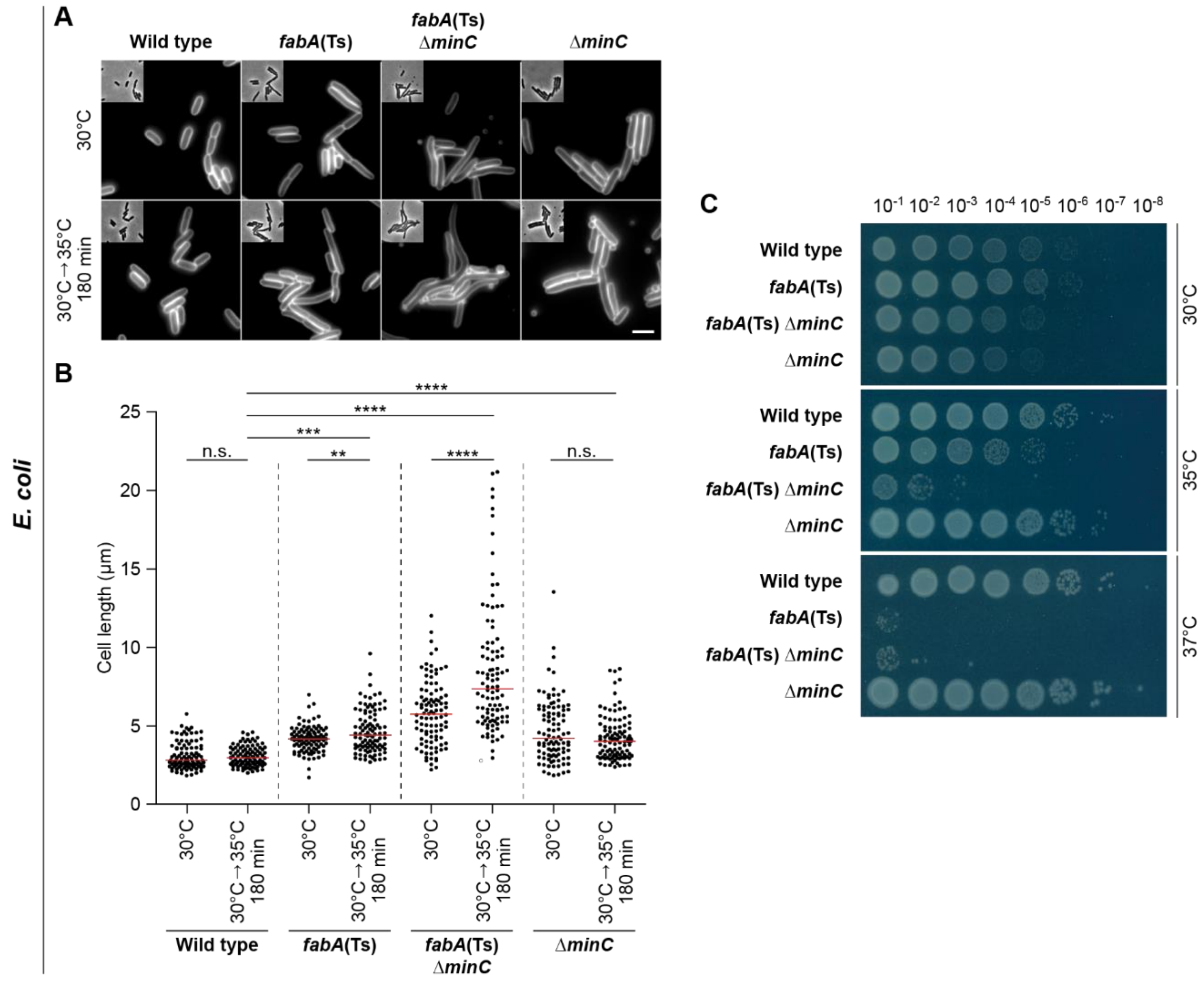

**Figure S8. Destabilisation of *E. coli* divisome by deletion of the division regulator *minC* triggers hypersensitivity towards low membrane fluidity.** (**A**) Images of *E. coli* wild type, *fabA*(Ts), *fabA*(Ts) Δ*minC and* Δ*minC cells* grown either at the permissive temperature (30°C) or for 180 min at 35°C, which is non-permissive for the *fabA*(Ts) Δ*minC* strain. Cells were stained with the outer membrane dye FM 5-95 prior to microscopy. Note the strong cell elongation of the *fabA*(Ts) Δ*minC* strain upon incubation at 35°C, which is indicative of a severe cell division defect. (**B**) Quantification of cell length for cells (n=100) depicted in panel A. (**C**) Viability of the strains depicted in panel **A** upon incubation on agar plates in M9-glucose minimal medium overnight at different temperatures. The serial dilutions and spot assays were carried out with pre-cultures grown at 30°C to mid-log growth phase. Note the temperature hypersensitivity and loss of viability of *fabA*(Ts) Δ*minC*-strain at 35°C, which indicates that the cell division process has become the limiting factor in tolerance towards low membrane fluidity in this strain. **Data information**: (**A**-**C**) The experiments are representative of biological triplicates. (**B**) Red lines indicate the median, while the P values represent results of unpaired, two-sided t-tests. Significance was assumed with **** $p < 0.0001$, *** $p < 0.001$, ** $p < 0.01$, * $p < 0.05$, n.s., not significant. (**A**) Scale bar: 3 µm. Strains used: (**A**-**C**) *E. coli* Y-Mel, UC1098, UC1098.Δ*minC*, JW1165-1.

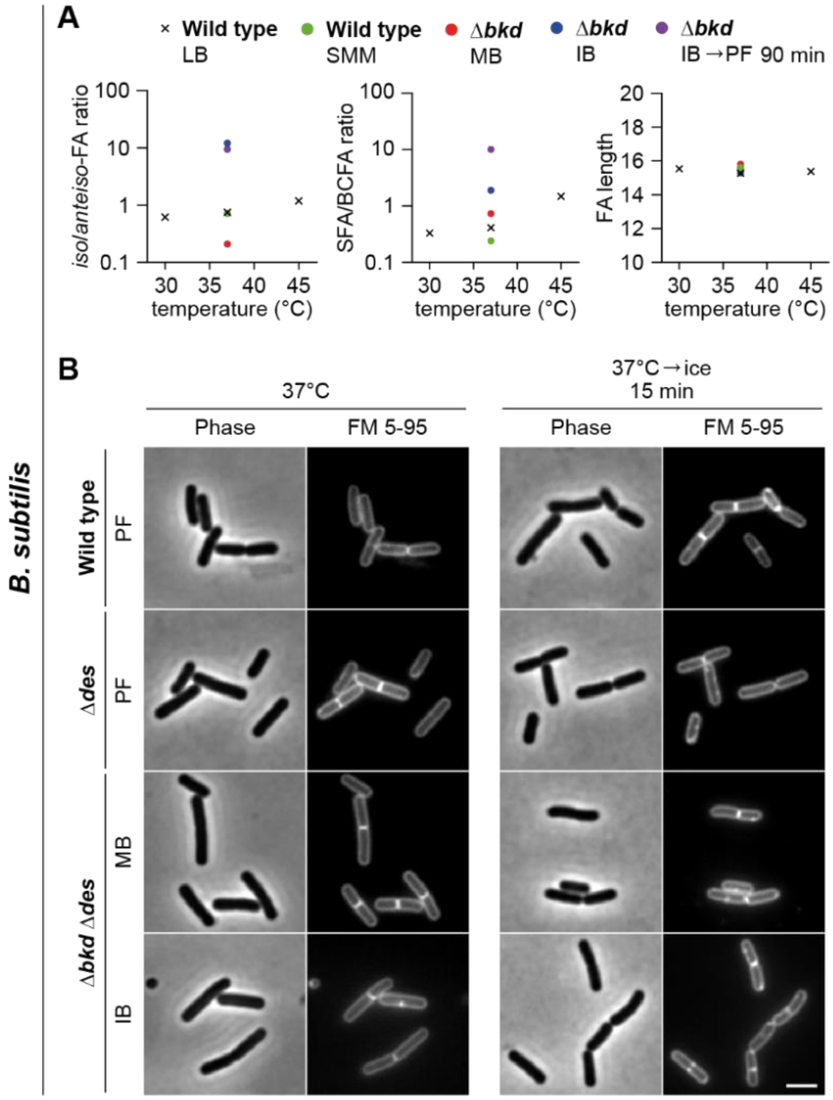

**Figure S9. Changes in *B. subtilis* fatty acid profile required to induce lipid phase separation are rather extreme.** (**A**) Comparison of changes in fatty acid (FA) composition observed for *B. subtilis* fatty acid precursor-auxotrophic Δ*bkd* strain upon changes in precursor availability, and *B. subtilis* wild type strain upon normal homeoviscous adaptation to different growth temperatures. Depicted are changes in the *iso/anteiso*-FA ratio (left panel), which is the main mechanism of membrane temperature adaptation in *B. subtilis*, changes in the ratio between SFA and BCFA (middle panel), and changes in the fatty acid chain length (right panel). The profiles were obtained for WT cells grown in LB medium or Spizizen minimal medium (SMM) at 30°C, 37°C and 45°C, respectively, and for the Δ*bkd* strain upon supplementation with 2-methyl butyric acid (MB), isobutyric acid (IB) or grown precursor-free (PF) for 90 min. Note that this is the same dataset as shown in Figures 1 and S1 for Δ*bkd.* While the temperature-dependent adaptation of *B. subtilis* fatty acid profile is observed both in terms of *iso/anteiso*-FA and SFA/BCFA ratio, the changes are minor compared to the more substantial changes induced by fatty acid precursor availability. Also note that, despite the large changes in the fatty acid types, the homeostasis of the fatty acid chain length remains unaffected. (**B**) Consistent with the large shifts in both *iso/anteiso*-FA and SFA/BCFA, required to induce large scale *in vivo* lipid phase separation, cold shock carried out by incubation on ice for 15 min is only capable for inducing minor membrane irregularities (compare with Figure 4A). The images depict phase contrast and fluorescent FM 5-95-stained wild type and ∆*des* cells, together with ∆*bkd* ∆*des* cells supplemented with MB or IB. The cells were grown at 37°C (left panels) followed by incubation on ice for 15 min (right panels). **Data information**: (**A**, **B**) The experiments are representative of biological duplicates. (**B**) Scale bar: 3 µm. Strains used: (**A**) *B. subtilis* 168, HS526; (**B**) *B. subtilis* 168, KS20, HS527.

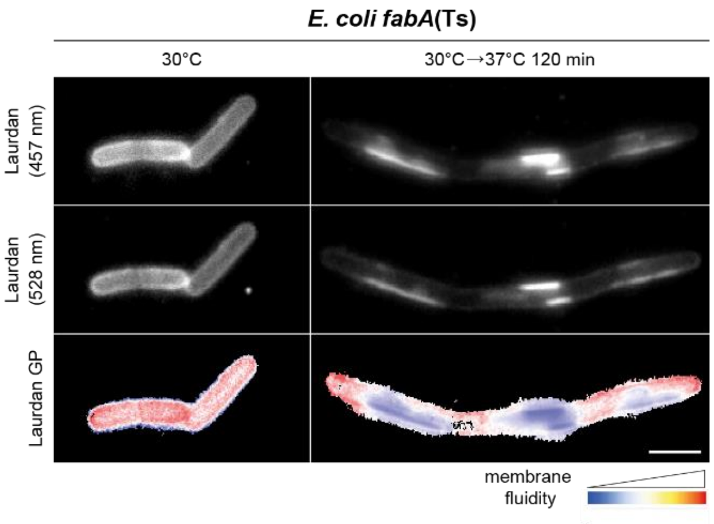

**Figure S10. Consequences of low membrane fluidity on membrane homogeneity.** Fluorescence images of *E. coli fabA*(Ts) cells grown in LB medium at permissive 30°C or shifted to the non-permissive temperature of 37°C for 120 min. Cells were stained with membrane fluidity-sensitive dye Laurdan and imaged at 457 nm, 528 nm and as the corresponding colour-coded Laurdan GP map. Data information: The experiment is representative of biological triplicates. Scale bar, 3 µm. Strain used: *E. coli* UC1098.

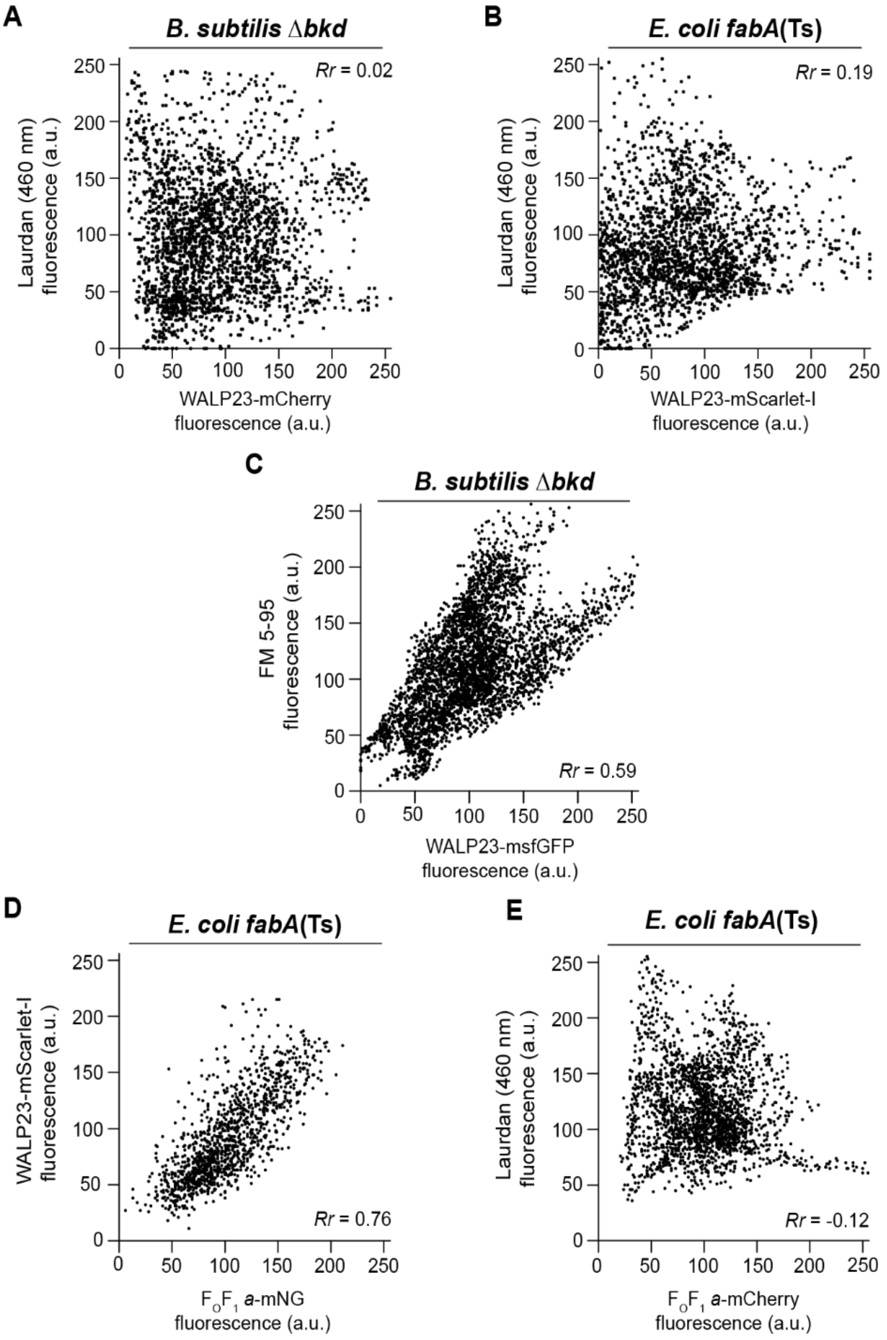

**Figure S11. Co-localisation correlation of the fluorescent dyes and marker proteins used in phase-separated membranes of *B. subtilis* and *E. coli.*** (**A**) Pixel-by-pixel intensity correlation between WALP23-mCherry and Laurdan (460 nm) fluorescent signals in *B. subtilis* Δ*bkd* cells grown for 90 min in the absence of fatty acid precursor (IB→PF) as shown in Figure 5D. The low Pearson's correlation coefficient (*Rr*) indicates lack of co-localisation. (**B**) Pixel-by-pixel intensity correlation between WALP23-mScarlet-I and Laurdan (460 nm) fluorescent signals in *E. coli fabA*(Ts) cells grown at 30°C and shifted for 120 min to the non-permissive temperature of 37°C as shown in Figure 5E. The low Pearson's correlation coefficient (*Rr*) again indicates lack of co-localisation. (**C**) Pixel-by-pixel intensity correlation between WALP23-msfGFP and FM 5-95 fluorescent signals in *B. subtilis* Δ*bkd* cells grown for 90 min in the absence of fatty acid precursor (IB→PF) as shown in Figure S12. The high *Rr* value indicates significant co-localisation. (**D**) Pixel-by-pixel intensity correlation between WALP23-mScarlet-I and $F_oF_1$ *a*-mNG in *E. coli fabA*(Ts) cells grown at 30°C and shifted for 120 min to the non-permissive temperature of 40°C as shown in Figure 6B. The high *Rr* value indicates significant co-localisation. (**E**) Pixel-by-pixel intensity correlation between $F_OF_1$ *a*-mCherry and Laurdan (460 nm) fluorescent signals in *E. coli fabA*(Ts) cells grown at 30°C and shifted for 120 min to the non-permissive temperature of 37°C as shown in Figure 6C. The low *Rr* value indicates lack of co-localisation. **Data information**: (**A**-**E**) Experiments are representative for three independent repeats. Strains used: (**A**) *B. subtilis* HS547; (**B**) *E. coli* UC1098/pBH501; (**C**) *B. subtilis* HS552; (**D**) *E. coli* MG4/pBH501; (**E**) *E. coli* LF6.red.

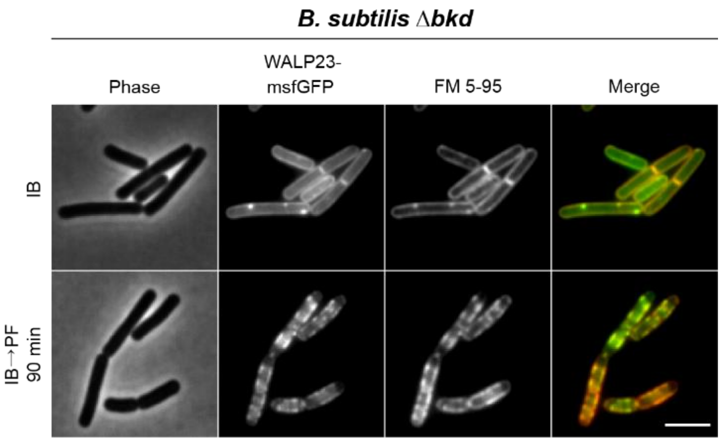

**Figure S12. Co-localisation of membrane dye FM 5-95 and transmembrane marker peptide WALP23 in phase-separated membranes of *B. subtilis.*** Phase contrast and fluorescence images of fatty acid precursor auxotroph *B. subtilis* strain Δ*bkd* grown in the presence of IB and in the absence of precursor for 90 min (IB→PF). Depicted are cells expressing WALP23-msfGFP, which were labelled with fluorescent dye FM 5-95. For fluorescence intensity correlations, see Figure S11C. Data information: The experiment is representative for three independent repeats. Strain used: *B. subtilis* HS552.

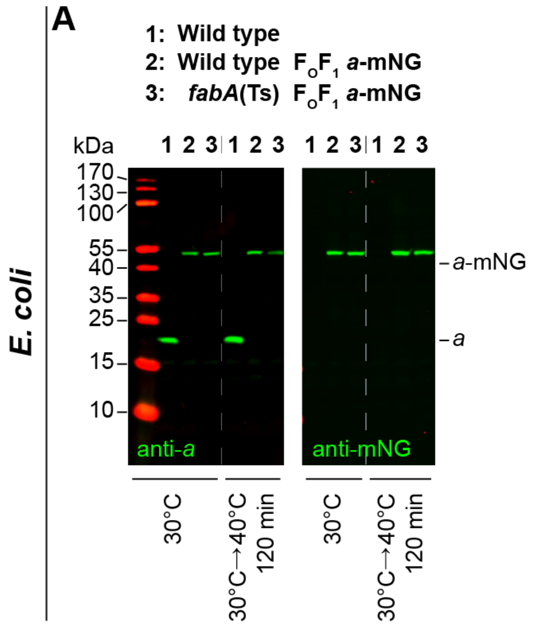

**Figure S13. Stability and activity of mNG-labelled $F_OF_1$ ATP synthase in *E. coli.*** (**A**) Stability of $F_OF_1$ *a*-mNG tested by Western blotting with antibodies specific for membrane-integral subunit $F_O$-*a* (GDH 14-5C6) or mNG (32F6) using inverted membrane vesicles prepared from *E. coli* wild type cells and from *fabA*(Ts) cells, respectively, both expressing $F_OF_1$ *a*-mNG. Cells were grown either at 30°C or grown at 30°C followed by transfer to 40°C for 120 min. (**B**) Functionality of $F_OF_1$ *a*-mNG verified by ATPase activities measured in inverted membrane vesicles in the absence or presence of $F_OF_1$ inhibitor *N, N*'-dicyclohexylcarbodiimide (DCCD). **Data information**: (**A**) The experiment is representative for technical triplicates. (**B**) The diagram depicts mean and SD of technical triplicates for each strain and condition. P values represent the results of unpaired, two-sided t-tests. Insignificant changes (p>0.1) are indicated with n.s., not significant. (**A**) *E. coli* Y-Mel, MG1, MG4; (**B**) *E. coli* MG1, MG4.

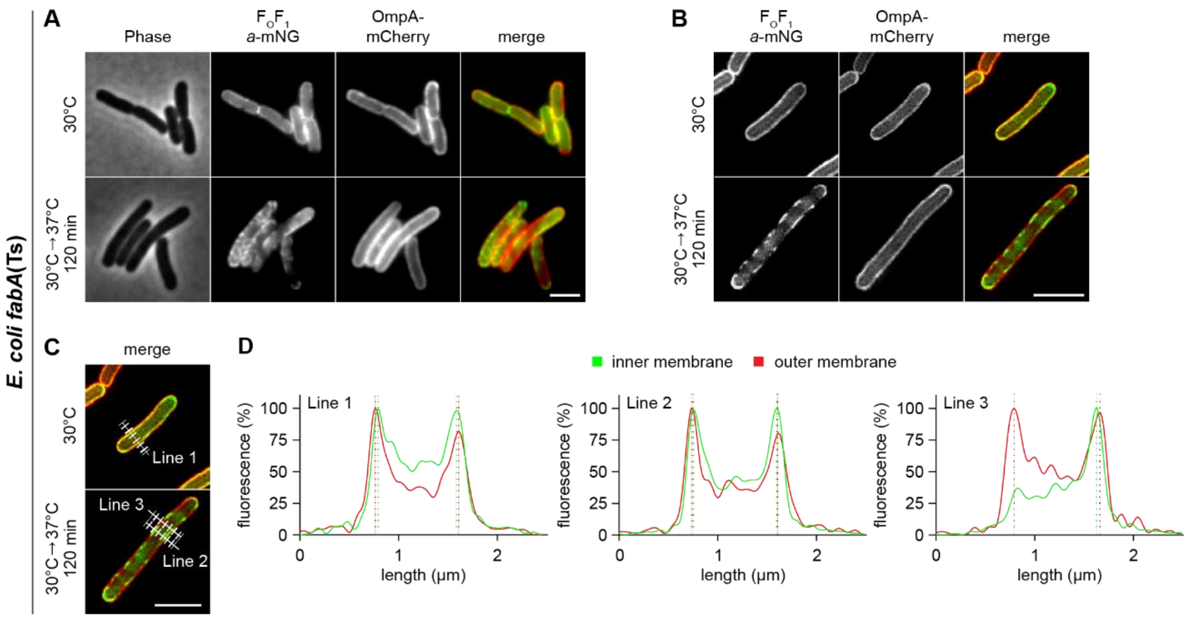

**Figure S14. Protein segregation induced by very low membrane fluidity is limited to the inner cytoplasmic membrane in *E. coli*.** (**A**) Membrane protein segregation was monitored in *fabA*(Ts) cells expressing as inner membrane marker $F_OF_1$ *a*-mNG and as outer membrane marker OmpA-mCherry. Widefield microscopy images depict phase contrast, fluorescence and overlay images for cells grown at permissive 30°C or at non-permissive 37°C for 120 min in M9-glucose minimal medium. Note the transition from disperse localisation into a segregated pattern in case of the inner membrane-localised $F_OF_1$ *a*-mNG at 37°C, while the pattern of the outer membrane-localised OmpA-mCherry remains homogeneous at both growth temperatures. (**B**) Super-resolution 2D-SIM (structured illumination microscopy) images of cells expressing $F_OF_1$ *a*-mNG and OmpA-mCherry at both growth temperatures. (**C**) Localisation and orientation of 5-pixel wide lines used to analyse fluorescence intensity profiles depicted in panel **D**. (**D**) Fluorescence intensity line scans across the cells imaged with 2D-SIM microscopy. Note the small, but detectable outward shift between the inner membrane marker $F_OF_1$ *a*-mNG and outer membrane marker OmpA-mCherry. **Data information**: (A-**D**) Experiments are representative of biological triplicates. (**A**-**C**) Scale bar: 3 µm. Strain used: (**A**-**D**) *E. coli* MG4/pGI10.

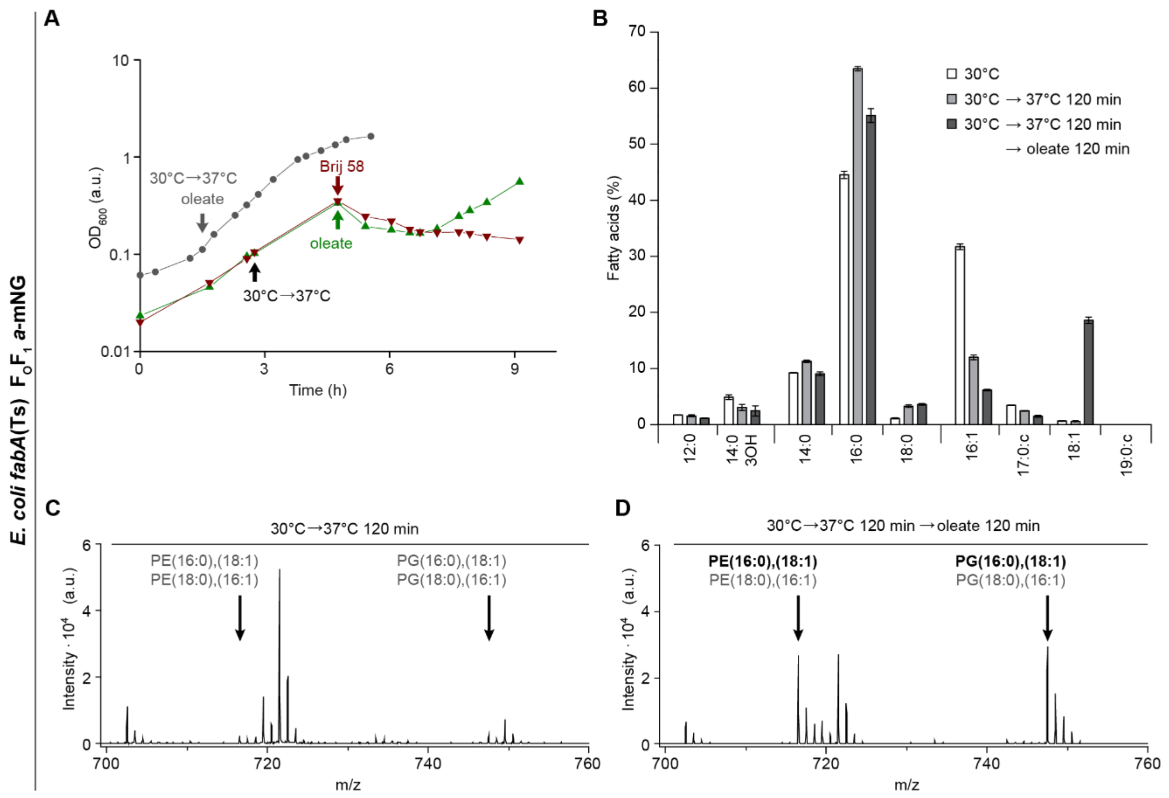

**Figure S15. Recovery of phase-separated plasma membranes of *E. coli fabA*(Ts) by lipid-incorporation of exogenous UFA oleate (*cis*-Δ9-C18:1).** (**A**) Growth curves depicting the growth inhibition of *E. coli fabA*(Ts) encoding $F_OF_1$ *a*-mNG upon transfer to the non-permissive temperature of 37°C, which can be rescued by addition of UFA oleate dissolved in Brij 58 (green triangles), but not by Brij 58 alone (dark red triangles). Supplementing the medium with oleate immediately upon shift to the non-permissive temperature prevents growth arrest (gray circles). (**B**) Detailed fatty acid profiles of $F_OF_1$ *a*-mNG-expressing *E. coli fabA*(Ts) upon growth at the permissive temperature of 30°C, upon depletion of UFA by a shift to non-permissive 37°C for 120 min and upon recovery by subsequent supplementation with oleate for additional 120 min at 37°C. (**C**) MALDI-TOF MS (matrix-assisted laser desorption ionization – time of flight mass spectrometry) spectra of lipids extracted from *fabA*(Ts) cells grown at 30°C and shifted to non-permissive 37°C for 120 min. (**D**) MALDI-TOF MS spectra of lipids extracted from *fabA*(Ts) cells grown at 30°C, shifted to 37°C for 120 min and subsequently supplemented with oleate for 120 min at 37°C. The intensities of *m/z*-values 716.5 and 747.5, only present in minute amounts after the 120 min-shift to 37°C (panel **C**), showed a strong increase after growth in the presence of oleate for 120 min. Both masses were identified by standard lipids as well as fragmentation of the mother ions (MALDI-TOF/TOF analyses) to be mainly PE(16:0),(18:1) and PG(16:0),(18:1), respectively, confirming the incorporation of oleate into phospholipids of *E. coli fabA*(Ts). **Data information**: (**A**-**D**) Results shown are representative for biological triplicates. (**B**) The histogram depicts mean and SD of biological triplicates. Strain used: (**A**-**D**) *E. coli* MG4.

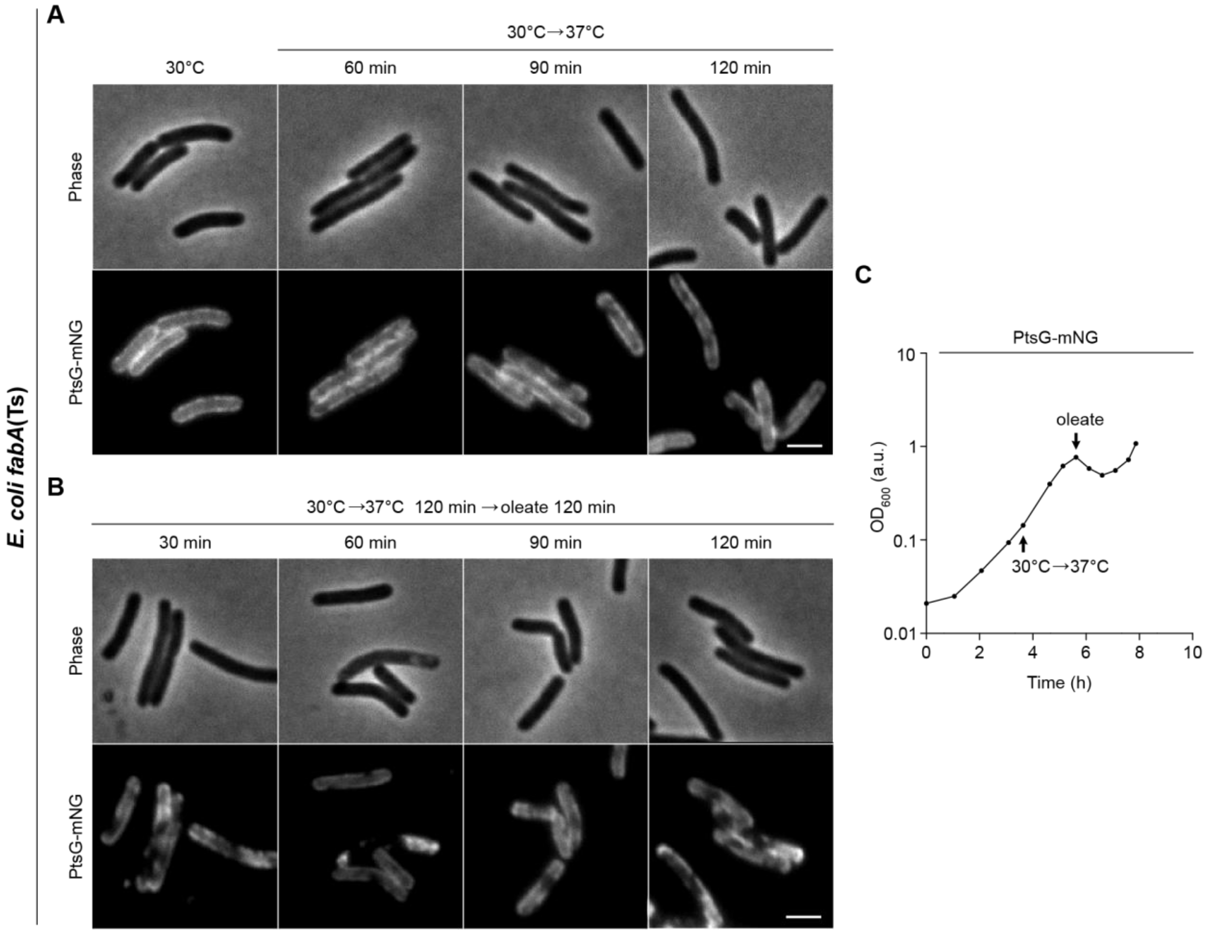

**Figure S16. Segregation of glucose permease (PtsG) into the fluid phase of phase-separated plasma membranes of *E. coli fabA*(Ts) and its re-localisation in the presence of UFA oleate.** (**A**) Phase and fluorescence images of *E. coli fabA*(Ts) cells chromosomally expressing mNG-labelled glucose permease PtsG. Depicted are cells grown at 30°C, and those shifted to the non-permissive temperature 37°C for the time points indicated, leading to PtsG-GFP partitioning within the membrane. (**B**) Phase and fluorescence images of the same strain as depicted in panel **A**, but imaged after supplementation of the growth medium with oleate at 37°C for further 120 min, leading to a reversion of the protein segregation observed in panel **A**. (**C**) Corresponding growth curve of *E. coli fabA*(Ts) expressing PtsG-mNG with the temperature shift (30°C→37°C) and the supplementation with oleate as indicated. **Data information**: (**A**-**C**) Results shown are representative for biological triplicates. (**A**, **B**) Scale bar: 2 µm. Strain used: (**A**-**C**) *E. coli* UC1098.PtsG-mNG.

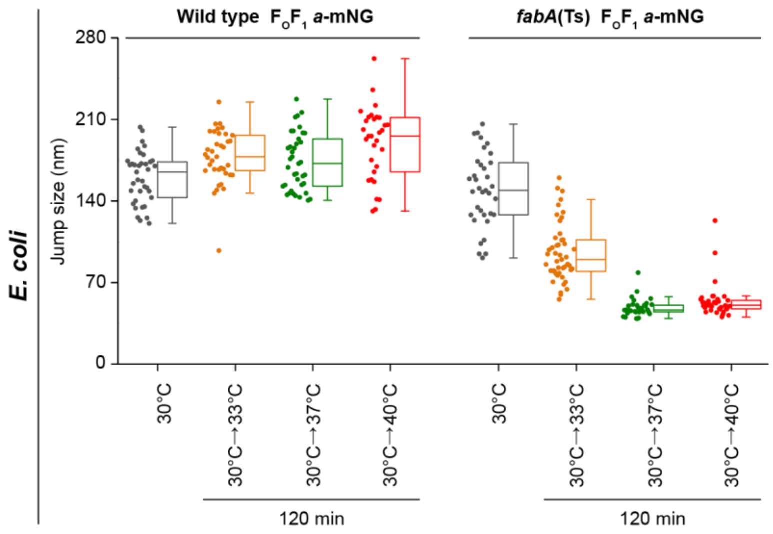

**Figure S17. Reduced lateral mobility of $F_OF_1$ *a*-mNG after UFA depletion in *E. coli fabA*(Ts).** Analysis of cell-to-cell heterogeneity in median jump sizes of $F_OF_1$ *a*-mNG. The box plot depicts the distribution of all trajectories determined for individual cells of *E. coli* wild type and *fabA*(Ts) at the growth temperatures indicated. Data information: The analysis was carried out with cells (n=31-47) from 3-5 biological replicates for each condition and strain, using 76$\pm$28 trajectories with $\geq$5 consecutive frames per cell. The interquartile range defines 50% of the data including the corresponding median, while the whiskers represent the borders of 25% and 75%, respectively, of the cells. Strains used: *E. coli* MG1, MG4.

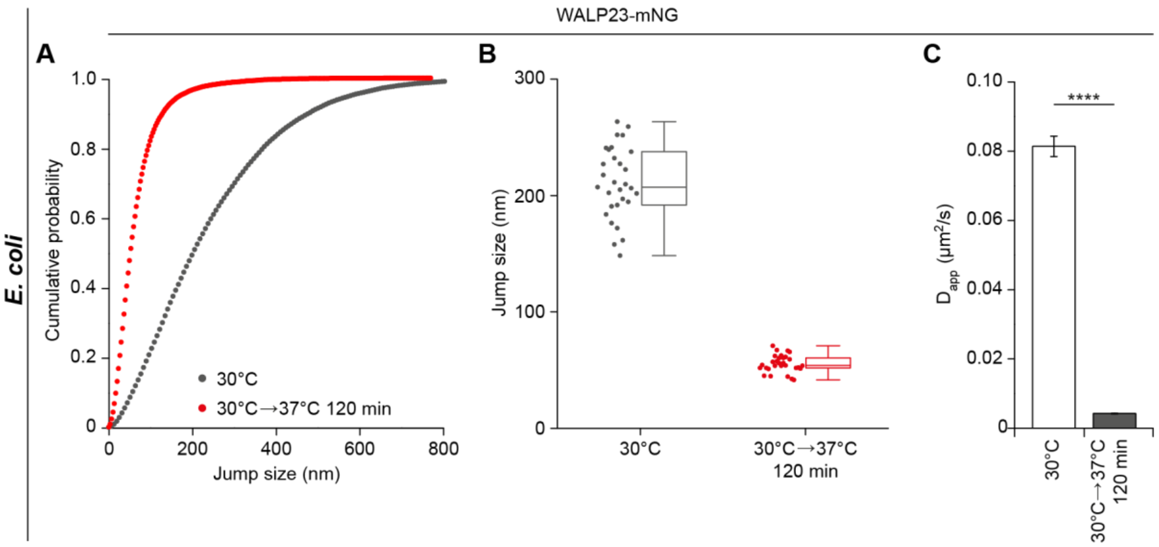

**Figure S18. Reduced lateral mobility of transmembrane peptide WALP23 after UFA depletion in *E. coli fabA*(Ts).** (**A**) Cumulative probability plot of jump sizes of mNG-labelled WALP23 in *fabA*(Ts) cells grown at permissive 30°C or shifted to the non-permissive temperature of 37°C for 120 min. (*B*) Analysis of the cell-to-cell heterogeneity in median jump sizes of WALP23-mNG. The box plot depicts the distribution of all trajectories determined for *E. coli* wild type and *fabA*(Ts) cells at the growth temperatures indicated. (C) Corresponding apparent diffusion coefficients ($D_{app}$) of WALP23-mNG calculated from trajectories analysed in panel **A**. **Data information**: (**A**) Trajectories with $\geq$5 consecutive frames, respectively, were pooled from biological triplicates and used for analysis (n=3305 (30°C); n=2801 (30°C→37°C)). (**B**) The analysis was carried out with cells (n=30) from biological triplicates for each condition. The interquartile range defines 50% of the data including the corresponding median, while the whiskers represent the borders of 25% and 75%, respectively, of the cells. (**C**) The histogram depicts mean and SD from biological triplicates, together with P values of a two-sided Wilcoxon rank sum test. Significance was assumed with **** p < 0.0001, *** p < 0.001, ** p < 0.01, * p < 0.05, n.s., not significant.  Strain used: (**A**-**C**) *E. coli* UC1098/pBH500.

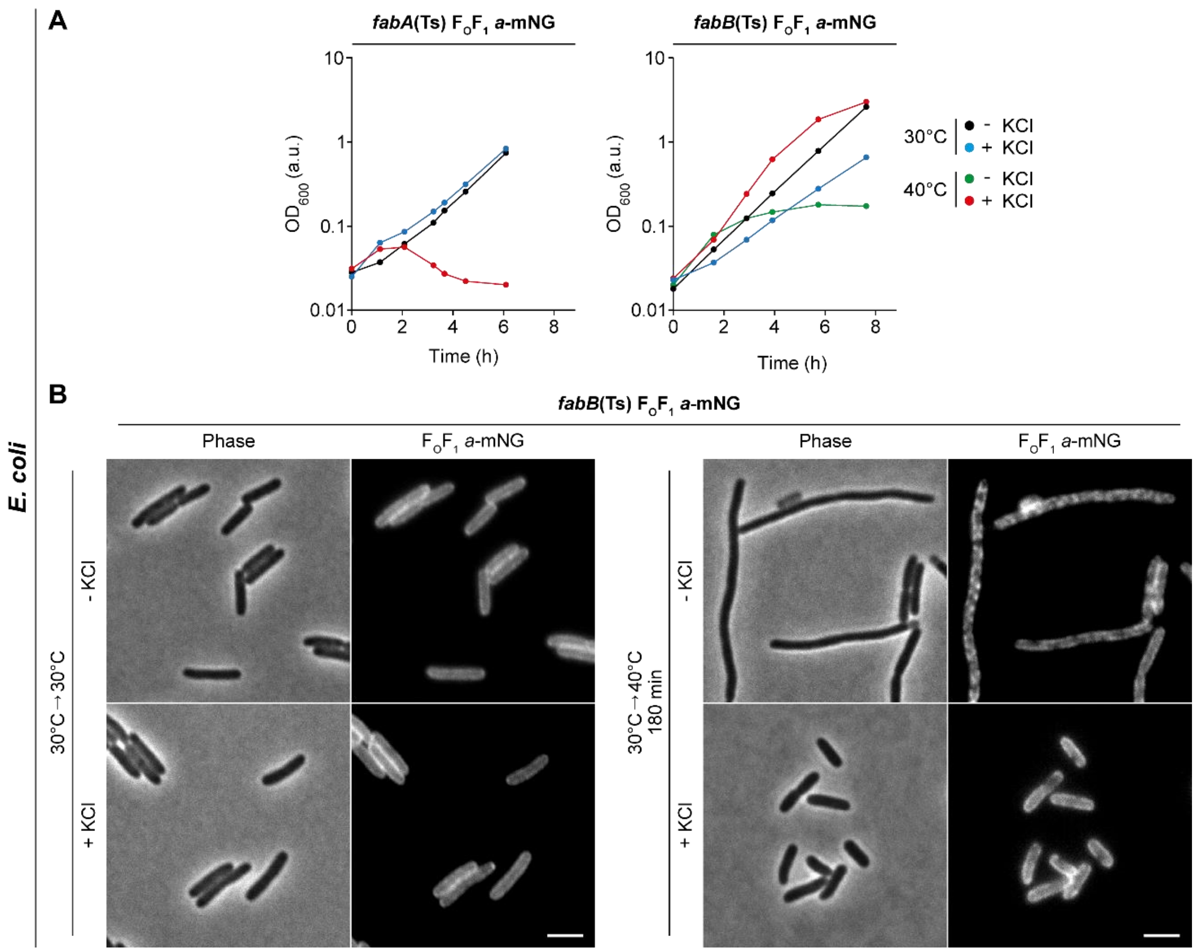

**Figure S19. Osmotic stabilisation of *E. coli fabB15*(Ts) enables cell growth at non-permissive temperature and prevents partially the partitioning of $F_OF_1$ *a*-mNG.** (**A**) Comparison of temperature-dependent growth behaviour of *E. coli* strains *fabA*(Ts) (left panel) and *fabB15*(Ts) (*fabF fabB15*(Ts); named "*fabB15*(Ts)" for simplicity throughout the text) (right panel) using KCl for osmotic stabilisation. Cells were grown in M9-glucose minimal medium in the absence or presence of 2% (w/v) KCl using for inoculation cultures grown overnight correspondingly at 30°C. Both strains grew comparably at 30°C. At non-permissive 40°C, *fabB15*(Ts) was able to grow in the presence of 2% KCl, while *fabA*(Ts) could not and cells started to lyse after 120 min. The inability of the *fabA*(Ts) strain UC1098 to retain growth through osmotic stabilisation is a well-known observation. From 17 tested UFA auxotrophic *E. coli* strains only 11 could maintain cell growth by osmotic stabilisation (Akamatsu, 1974; Broekman & Steenbakkers, 1973). This demonstrates that growth at non-permissive temperature in a high-osmotic medium is not a common characteristic of mutants impaired in the biosynthesis of unsaturated fatty acids. (**B**) Phase and fluorescence images of *E. coli fabB15*(Ts) cells chromosomally expressing $F_OF_1$ *a*-mNG. At 30°C cells exhibited a homogeneous distribution of $F_OF_1$ *a*-mNG independent of KCl supplementation. Cells shifted to 40°C for 180 min without KCl supplementation developed significant partitioning of $F_OF_1$ *a*-mNG, which was largely surpassed by KCl. Also note the clear increase in cell length of *fabB15*(Ts) observed without KCl supplementation at 40°C indicating severe cell division defects. **Data information**: (**A**, **B**) Experiments are representative of three independent repeats. (**B**) Scale bar: 2 µm. Strains used: (**A**) *E. coli* MG1, BHH87; (**B**) *E. coli* BHH87.

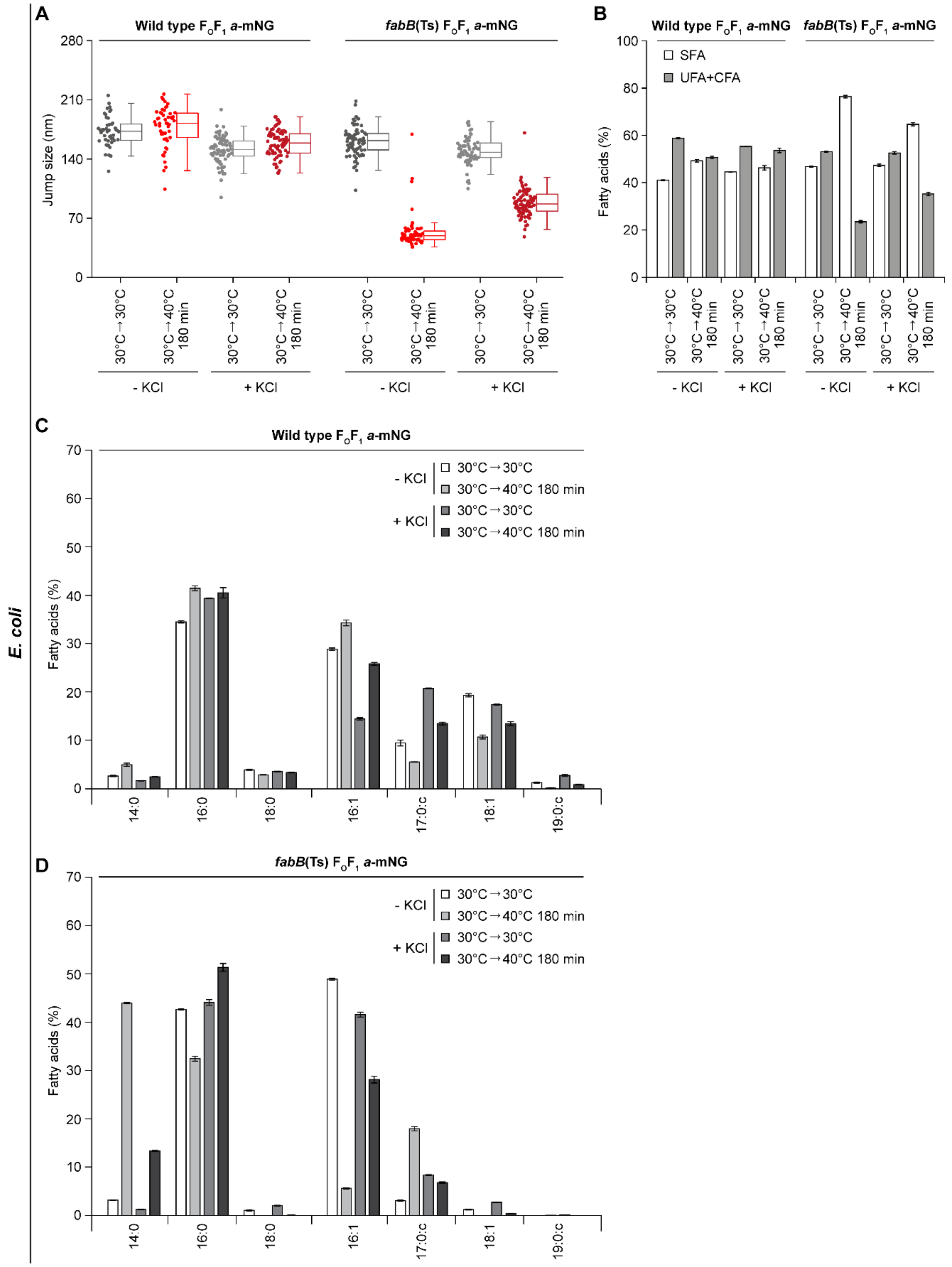

**Figure S20. Osmotic stabilisation of lateral diffusion of $F_OF_1$ *a*-mNG in UFA-depleted membranes of *E. coli fabB15*(Ts).** (**A**) Lateral mobility of $F_OF_1$ *a*-mNG upon osmotic KCl stabilisation. Distribution of median jump sizes of $F_OF_1$ *a*-mNG in different cells. The box plot depicts the distribution of all trajectories determined for *E. coli* WT and *fabB15*(Ts) cells at growth temperatures and KCl supplementation as indicated. Supplementation of M9-glucose minimal medium with 2% (w/v) KCl resulted in a slight decrease in lateral mobility of $F_OF_1$ *a*-mNG in WT cells grown at 30°C or grown at 40°C for 180 min. The same was observed for *fabB15*(Ts) cells grown at permissive 30°C. Cells of *fabB15*(Ts) grown at non-permissive 40°C for 180 min in the presence of 2% KCl maintained intermediate lateral diffusion rates for $F_OF_1$ *a*-mNG (app. 60% compared to *fabB15*(Ts) cells grown at 30°C without KCl), whereas at 40°C for 180 min without KCl supplementation, a more substantial reduction of lateral displacement was observed, comparable to that monitored for *fabA*(Ts) (see Figures 7 and S17). (**B**) Fatty acid composition (SFA versus UFA/CFA ratio) of *E. coli* WT and *fabB15*(Ts) cells. The same cell cultures as for imaging in panel **A** were analysed. *E. coli* WT cells at both temperatures and *fabB15*(Ts) cells grown at 30°C were essentially unaffected in their fatty acid composition from osmotic changes in the medium. Cells of *fabB15*(Ts) grown at 40°C for 180 min showed a strong decrease in UFA in the absence of additional KCl, comparable to *fabA*(Ts) (see Figure 1E), while in the presence of KCl the amount of UFA+CFA remained at intermediate levels. (**C**, **D**) Detailed fatty acid composition of *E. coli* WT cells (**C**) and *E. coli fabB15*(Ts) (**D**) grown as described in panel **A**. Note the high content of C14:0 in the absence of KCl at 40°C, which may be an adaptive response to fluidise the membrane upon low UFA conditions; an observation also described for a different *fabB* strain (Budin *et al*, 2018). In high-osmotic medium, the higher content of UFA present may be due to an osmotic stabilisation of the temperature-sensitive enzyme carrying the substitution A329V located within the interior of each monomer of the homodimeric enzyme (Olsen *et al*, 1999). Fatty acid compositions were calculated from lipid species analyses determined by MALDI-TOF/TOF MS. **Data information**: (**A**) The analysis was carried out for n=47-88 cells from biological triplicates for each condition and strain, using 100±25 trajectories with ≥5 consecutive frames per cell. The interquartile range defines 50% of the data including the corresponding median, while the whiskers represent the borders of 25% and 75%, respectively. (**B**-**D**) Graphs depict mean and SD of biological triplicates. Strains used: (**A**, **B**) *E. coli* MG1, BHH87; (**C**) *E. coli* MG1; (**D**) *E. coli* BHH87.

**Table S1. Strains. Xyl, xylose.**

| Strain | Relevant Genotype | Induction | Source/Reference |
|---|---|---|---|
| *B. subtilis* 168 | *trpC2* | - | (Barbe *et al*, 2009) |
| *B. subtilis* HB5134 | *des::spc* | - | (Hachmann *et al*, 2009) |
| *B. subtilis* LC5 | *des::kan* | - | (Altabe et al, 2003) |
| *B. subtilis* KS20 | *des::spc* | - | This work |
| *B. subtilis* RM113 | *spoVD::cat Pxyl-murE sepFT11M bkd::ery* | - | (Mercier *et al*, 2012) |
| *B. subtilis* HS526 | *bkd::ery* | - | This work |
| *B. subtilis* HS527 | *bkd::ery des::spc* | - | This work |
| *B. subtilis* HS550 | *bkd::ery des::kan* | - | This work |
| *B. subtilis* JWV042 | *amyE::cat Phbs-hbs-gfp* | - | (Strahl & Hamoen, 2010) |
| *B. subtilis* HS541 | *amyE::cat Phbs-hbs-gfp bkd:ery des::spc* | - | This work |
| *B. subtilis* 2020 | *amyE::spc Pxyl-gfp-ftsZ* | - | (Stokes *et al*, 2005) |
| *B. subtilis* HS548 | *amyE::spc Pxyl-gfp-ftsZ bkd::ery des::kan* | 0.3% Xyl | This work |
| *B. subtilis* KS69 | *amyE::spec Pxyl-msfGFP-MreB* | - | (Scheinpflug *et al*, 2017) |
| *B. subtilis* HS549 | *amyE::spec Pxyl-msfGFP-MreB bkd::ery des::kan* | 1% Xyl | This work |
| *B. subtilis* HS547 | *amyE::spec Pxyl-WALP23-mCherry bkd::ery des::kan* | 1% Xyl | This work |
| *B. subtilis* JG054 | *amyE::spec Pxyl-WALP23-msfGFP* | - | This work |
| *B. subtilis* HS552 | *amyE::spec Pxyl-WALP23-msfGFP bkd::ery des::kan* | 1% Xyl | This work |
| *E. coli* Y-Mel | $\lambda^+$ $F^+$ *mel-1 supF58* (wild type) | - | (Rickenberg & Lester, 1955) |
| *E. coli* MG1655 | $\lambda^+$ $F^-$ *ilvG- rfb-50 rph-1* (wild type) | - | (Blattner *et al*, 1997) |
| *E. coli* AB1 | $F^-$ *fhuA22 fabB15*(Ts) *fabF200 zcf-229*::Tn*10 gyrA220*($Nal^R$) *rpsL146*($Sm^R$) Δ*atpIBE*::FRT-*kan*-FRT ($Kan^R$) | - | This work |
| *E. coli* BHH87 | $F^-$ *fhuA22 fabB15*(Ts) *fabF200 zcf-229*::Tn*10 gyrA220*($Nal^R$) *rpsL146*(Sm) *atpB-mNeonGreen* | - | This work |
| *E. coli* BHH100 | $F^+$ $Sm^R$ *fabA*(Ts) *fabF* $Crc^-$ *ftsZ*$^{55}$*-msfGFP-*$^{56}$*ftsZ* ($Cm^R$) | - | This work |
| *E. coli* BHH101 | $F^+$ $Sm^R$ *fabA*(Ts) *fabF* $Crc^-$ *mreB*$^{226}$*-msfGFP-*$^{227}$*mreB csrD-mreB::*FRT-*kan*-FRT ($Kan^R$) | - | This work |
| *E. coli* CY288 | F- *fhuA22 fabB15*(Ts) *fabF200 zcf-229*::Tn*10 gyrA220*($Nal^R$) *rpsL146*($Sm^R$) | | (Garwin *et al*, 1980) |
| *E. coli* EB4 | $\lambda^+$ $F^+$ *mel-1 supF58* Δ*atpI'BE'::kan* | - | (Renz *et al*, 2015) |
| *E. coli* EB8.1 | $\lambda^+$ $F^+$ *mel-1 supF58 atpB-mCherry* | - | This work |
| *E. coli* JW0334-1 | $\lambda^-$ $F^-$ *rph-1 hsdR514* Δ*lacY784::*FRT-*kan*-FRT Δ*(araD-araB)567* Δ*lacZ4787*(::*rrnB-3*) Δ*(rhaD-rhaB)568* ($Kan^R$) | - | (Baba *et al*, 2006) |
| *E. coli* JW1087-2 | $\lambda^-$ $F^-$ *rph-1 hsdR514* Δ*ptsG763::*FRT-*kan*-FRT Δ*(araD-araB)567* Δ*lacZ4787*(::*rrnB-3*) Δ*(rhaD-rhaB)568* ($Kan^R$) | - | (Baba *et al*, 2006) |
| *E. coli* JW1165-1 | $\lambda^-$ $F^-$ *rph-1 hsdR514* Δ*minC765::*FRT-*kan*-FRT Δ(*araD-araB*)567 Δ*lacZ4787*(::*rrnB-3*) Δ(*rhaD-rhaB*)568 ($Kan^R$) | - | (Baba *et al*, 2006) |
| *E. coli* KC555 | $\lambda^-$ $F^-$ *rph-1 ftsZ*$^{55}$*-msfGFP-*$^{56}$*ftsZ* ($Cm^R$) | - | (Yang *et al*, 2017) |
| *E. coli* LF4 | $F^+$ $Sm^R$ *fabA*(Ts) *fabF* $Crc^-$ Δ*atpIBE::FRT-kan-FRT* ($Kan^R$) | - | This work |

| | | | |
|---|---|---|---|
| *E. coli* LF6.red | $F^+$ $Sm^R$ *fabA*(Ts) *fabF* $Crc^-$ *atpB-mCherry* | - | This work |
| *E. coli* MG1 | $\lambda^+$ $F^+$ *mel-1 supF58 atpB-mNeonGreen* | - | This work |
| *E. coli* MG4 | $F^+$ $Sm^R$ *fabA*(Ts) *fabF* $Crc^-$ *atpB-mNeonGreen* | - | This work |
| *E. coli* MG1655 mreB-msfGFP | $\lambda^-$ $F^-$ *rph-1* $mreB^{226}$*-msfGFP-*$^{227}$*mreB* *csrD-mreB*::FRT-*kan*-FRT ($Kan^R$) | - | (Ursell *et al*, 2014) |
| *E. coli* UC1098 | $F^+$ $Sm^R$ *fabA*(Ts) *fabF* $Crc^-$ | - | (Cronan *et al*, 1972) |
| *E. coli* UC1098.PtsG-mNG | $F^+$ $Sm^R$ *fabA*(Ts) *fabF* $Crc^-$ *ptsG-mNeonGreen* | - | This work |
| *E. coli* UC1098.Δ*ptsG* | $F^+$ $Sm^R$ *fabA*(Ts) *fabF* $Crc^-$ Δ*ptsG763::FRT-kan-FRT* ($Kan^R$) | - | This work |
| *E. coli* UC1098.Δ*lacY* | $F^+$ $Sm^R$ *fabA*(Ts) *fabF* Δ*lacY784*::FRT-*kan*-FRT ($Kan^R$) | - | This work |
| *E. coli* UC1098.*ΔminC* | $F^+$ $Sm^R$ *fabA(Ts) fabF* $Crc^-$ Δ*minC765*::FRT-*kan*-FRT ($Kan^R$) | - | This work |
| *E. coli* Y-Mel.Δ*lacY* | $\lambda^+$ $F^+$ *mel-1 supF58 ΔlacY784::*FRT-*kan*-FRT ($Kan^R$) | - | This work |

**Table S2. Oligonucleotides.** Deoxynucleotides corresponding to the genes encoding fluorescent proteins are in italics; restriction sites are underlined.

| Oligo | Sequence (5‘ → 3‘) |
|---|---|
| 1 | GCA ACT TCT CCA ATG ATC TGA AG |
| 2 | GCA TAA CCG ATA CCG ACG ATC GG |
| 3 | CCG ATC GTC GGT ATC GGT TAT GCG |
| 4 | *ATC CTC CTC GCC CTT GCT CAC CAT* TCT AGA TGC CGG AGT TGG AGC CG |
| 5 | CGG CTC CAA CTC CGG CAT CTA GA*A TGG TGA GCA AGG GCG AGG AGG AT* |
| 6 | GTC TCC CCA ACG TCT TAC GGA TTA *CTT GTA CAG CTC GTC CAT GCC* |
| 7 | *GGC ATG GAC GAG CTG TAC AAG* TAA TCC GTA AGA CGT TGG GGA GAC |
| 8 | GTC AAT AAC CTG TTC GAC AAA ACC |
| 9 | GAA TTC TCA TGT TTG ACA GC |
| 10 | *GTT ATC CTC CTC GCC CTT GCT CAC CAT* GGA TCC ATG ATC TTC AGA CGC |
| 11 | GCG TCT GAA GAT CAT GGA TCC *ATG GTG AGC AAG GGC GAG GAG GAT AAC* |
| 12 | CGT AGT AGT GTT GGT AAA *TTA CTT GTA CAG CTC GTC CAT GCC C* |
| 13 | *CGG CAT GGA CGA GCT GTA CAA G*TA ATT TAC CAA CAC TAC TAC G |
| 14 | CGC TGG CCT TTG CAA GGT CAA GGT CCT TAT GTG CTC ATT CTG CGG |
| 15 | GGA GAT TCC TAG GAT GGC TTG GTG G |
| 16 | *GTT ATC CTC CTC GCC CTT GCT CAC CAT* TCC TGA GCC GCT TCC TGA CGC C |
| 17 | *GAT TAC GGC CTC TCC TTT GGA GAC CAT* TCC TGA GCC GCT TCC TGA CGC C |
| 18 | GGC GTC AGG AAG CGG CTC AGG A*AT GGT GAG CAA GGG CGA GGA GGA TAA C* |
| 19 | GCT CTA GAA CTA GTG GAT CTG AAG TCT GGA CAT *TTA CTT GTA CAG CTC GTC CAT GCC C* |
| 20 | GGC GTC AGG AAG CGG CTC AGG A*AT GGT CTC CAA AGG AGA GGC CGT AAT C* |
| 21 | GCT CTA GAA CTA GTG GAT CTG AAG TCT GGA CAT *TTA TTT ATA CAG CTC ATC CAT ACC AC* |
| 22 | CAT TCC TGA GCC GCT TC |
| 23 | GAC TTC AGA TCC ACT AGT TCT AGA G |
| 24 | CTA GAA CTA GTG GAT CTG AAG TC |
| 25 | AGC GGC TCA GGA ATG AGC AAA GGA GAA GAA CTT TTC |
| 26 | GTA CCC GGG GAT CCT C |
| 27 | CAT GGT GAA TTC CTC CTG C |
| 28 | GCA GGA GGA ATT CAC CAT GAA AAG AAT GTT AAT CAA CGC AAC |
| 29 | AGG ATC CCC GGG TAC TTA CTC AAC AGG TTG CGG AC |
| 30 | TTC TTC ACC ACC GCT TGC AGG AGC TGC TGG TG |
| 31 | AGC GGT GGT GAA GAA ACC AAA C |

**Table S3. Plasmids.** IPTG, isopropyl-β-D-thiogalactopyranosid; Ara, arabinose

| Plasmid | Relevant Genotype | Induction | Source |
|---|---|---|---|
| pBH4 | *ori*-ColE1 *bla* Δ*rop* ($Ap^R$) (pBR322) *Patp3-atpBEFHAGDC* | - | This work |
| pBH189 | *ori*-ColE1 *bla* Δ*rop* ($Ap^R$) (pBR322) *Patp3-atpBEFHAGDC atpB-mNG* | - | This work |
| pBH500 | *ori*-ColE1 *bla spc* ($Ap^R$ $Sp^R$) (pSG1154) *amyE*::*Pxyl-WALP23-mNeonGreen* | - | This work |
| pBH501 | *ori*-ColE1 *bla spc* ($Ap^R$ $Sp^R$) (pSG1154) *amyE::Pxyl-WALP23-mScarlet-I* | - | This work |
| pBLP2 | *ori*-ColE1 *bla* ($Ap^R$) (pBAD24) *araC ParaBAD-ptsG-GFP* | - | (Kosfeld & Jahreis, 2012) |
| pBWU13 | *ori*-ColE1 *bla* ($Ap^R$) (pBR322) *Patp3-atpBEFHAGDC* | - | (Moriyama *et al*, 1991) |
| pEB21.2 | *ori*-ColE1 *bla* ($Ap^R$) (pBR322) *Patp3-atpBEFHAGDC atpB-mCherry* | - | This work |
| pGI10 | *ori*-ColE1 *bla* ($Ap^R$) *Ptrc-ompA-(LEDPPAEF)-mCherry* | 0.1 mM IPTG | (Verhoeven *et al*, 2013) |
| pJG130 | *ori*-ColE1 *bla* ($Ap^R$) (pBAD322) *ParaBAD-rne* | 0.2% Ara | This work |
| pJG131 | *ori*-ColE1 *bla* ($Ap^R$) (pBAD322) *ParaBAD-rne(*ΔAH*)* | 0.2% Ara | This work |
| pKD4 | *ori* ColE1 *bla FRT-kan-FRT* ($Ap^R$, $Kan^R$) (pANTSγ) | - | (Datsenko & Wanner, 2000) |
| pKD46 | *ori-R101 repA101*(Ts) *bla* ($Ap^R$) *araC ParaBAD-gam-beta-exo* | 0.2% Ara | (Datsenko & Wanner, 2000) |
| pL030 | *ori*-ColE1 *bla spc* ($Ap^R$ $Sp^R$) (pSG1154) *amyE::Pxyl-WALP23-mCherry* | - | S. Lee |
| pNCS-mNeonGreen | *ori*-ColE1 *bla* ($Ap^R$) (pNCS) *T7-tag* $His_6$*-mNG* (constitutive) | - | (Shaner *et al*, 2013) |
| pQW58 | *ori*-ColE1 *bla* ($Ap^R$) *$lacI^q$ Plac-mCherry* | | (Galli & Gerdes, 2010) |
| pSD166 | *ori*-p15A *bla* ($Ap^R$) (pACYC177) *Patp3-atpBEFHAGDC atpB-EGFP* | - | (Düser *et al*, 2008) |
| pSG1154 | *ori*-ColE1 *bla spc amyE*'-'*amyE* ($Ap^R$ $Spec^R$) | - | (Lewis & Marston, 1999) |
| pSTK3 | *ori*-ColE1 *bla* ($Ap^R$) (pBR322) *Patp3-atpBEFHAGDC atpF(C21A)* | - | (Brandt *et al*, 2013) |
| pTM30.LacZ-His2 | *ori*-ColE1 *bla* ($Ap^R$) *Ptac-OlacZ-lacZ-His2* | - | K. Jahreis |
| pVK207 | *ori*-pSC101 *spc* ($Sp^R$) *Prne-rne-yfp* | - | (Khemici *et al*, 2008) |

### Movie legends

**Movie S1 (separate file). Time lapse microscopy of fatty acid precursor auxotroph *B. subtilis* Δ*bkd* strain depleted for branched chain fatty acids.** Cells labelled with membrane dye FM 5-95 were grown in the presence of precursor IB, washed precursor-free (PF) and transferred to time lapse slides prepared with PF medium or IB-supplemented medium. **Data information**: Cells were imaged for 250 min at 5 min intervals and 350 ms exposure time per frame. The movie frame rate is 5 frames per second. Scale bar, 3 µm. Strain used: *B. subtilis* HS527.

**Movie S2 (separate file). Time lapse microscopy of the thermosensitive *E. coli fabA*(Ts) strain depleted for unsaturated fatty acids.** Cells chromosomally expressing mNG-labelled ATP synthase ($F_OF_1$ *a*-mNG) were grown at 30°C, transferred to time lapse slides with the same medium and grown at permissive 30°C or at non-permissive 40°C for depletion of UFA. **Data information**: Cells were imaged for 150 min in 5 min intervals and 50 ms exposure time per frame. The movie frame rate is 5 frames per second. Deconvolution (only for the merge frames) was performed using SoftWoRx software. Scale bar, 2 µm. Strain used: MG4.

**Movie S3 (separate file). Trajectory maps of *in vivo* single molecule tracking of mNG-labelled ATP synthase ($F_OF_1$ *a*-mNG) in *E. coli* wild type cells.** Cells chromosomally expressing $F_OF_1$ *a*-mNG were grown at 30°C or grown at 30°C and shifted to 33°C, 37°C or 40°C for 120 min as indicated. **Data information**: The movie shows sequential frames with a frame binning of 2 and 15 frames per second. All trajectories of $F_OF_1$ *a*-mNG complexes with ≥5 consecutive frames are shown. Scale bar, 1 µm. Strain used: *E. coli* MG1.

**Movie S4 (separate file). Trajectory maps of *in vivo* single molecule tracking of mNG-labelled ATP synthase ($F_OF_1$ *a*-mNG) in thermosensitive *E. coli fabA*(Ts) after depletion of unsaturated fatty acids.** Cells chromosomally expressing $F_OF_1$ *a*-mNG were grown at 30°C or grown at 30°C and shifted to 33°C, 37°C, or 40°C for 120 min as indicated. **Data information**: The movie shows sequential frames with a frame binning of 2 and 15 frames per second. All trajectories of $F_OF_1$ *a*-mNG complexes with ≥5 consecutive frames are shown. Scale bar, 1 µm. Strain used: *E. coli* MG4.
